# Implementing N-terminomics and machine learning to probe *in vivo* Nt-arginylation

**DOI:** 10.1101/2025.04.09.646507

**Authors:** Shinyeong Ju, Laxman Nawale, Seonjeong Lee, Jung Gi Kim, Hankyul Lee, Narae Park, Dong Hyun Kim, Hyunjoo Cha-Molstad, Cheolju Lee

## Abstract

N-terminal arginylation (Nt-arginylation) serves as a protein degradation signal in both the ubiquitin-proteasome system and the autophagy-lysosomal pathway. However, the scarcity of arginylated proteins in cells and limitations in current identification methods have hindered progress in this field. In this study, we developed a novel integrated approach that combines N-terminomics (N-terminomic enrichment, LC-MS/MS, and tandem database search) with advanced machine learning-based filtering strategies to successfully identify *in vivo* Nt-arginylation with unprecedented sensitivity and specificity. By utilizing Arg-starting peptides from missed cleavage products as physicochemical proxies for ATE1-mediated Nt-arginylation, we trained a transfer learning-based model to predict MS2 spectra and retention times of candidate peptides. Additionally, near-isobaric modifications were filtered by statistically analyzing mass error deviations in MS2 fragment ions. Using this approach, we identified 134 Nt-arginylation sites in thapsigargin-treated HeLa cells, revealing a significant increase of Nt-arginylome under unfolded protein response (UPR) stress. Arginylated proteins originate from various organelles, including ER, nucleus and mitochondria. Notably, arginylation frequently occurred at sites processed by caspases or where signal peptides had been cleaved. Several proteins identified in our arginylome study were validated for their interaction with the R-catcher, an Nt-Arginylation bait derived from the p62 ZZ domain. Temporal profiling of N-terminal arginylation sites post-stress induction via targeted proteomics revealed a sequential response pattern. The UPR signal transducer ATF4 exhibited the most rapid increase, followed by N-terminal arginylation of caspase-3 substrates, and subsequently by arginylation at signal peptide cleavage sites among endoplasmic reticulum proteins. Our novel machine learning-based filtering methodologies enable the discovery of rare post-translational modifications by implementing specialized filtering strategies tailored to the unique physicochemical properties of terminal modifications, with potential applications in biomarker discovery, drug target identification, and elucidation of disease-specific protein regulation mechanism.

## Main

Protein Nt-arginylation is mediated by ATE1 (Arginyl-tRNA-protein transferase 1)^1^ for degradation through the ubiquitin-proteasome system as a pivotal component of the Arg/N-degron pathway^2^. Nt-arginylation not only promotes the degradation of short-lived protein fragments but also triggers autophagic processes in a concentration-dependent manner^3, 4^. Disruption of ATE1 leads to defects in cardiovascular development and angiogenesis^5^, neurodegeneration including Parkinson’s and Alzheimer’s disease^6, 7^, and carcinogenesis^8^. The ability to identify Nt-arginylation is thus a key to understanding the mechanism behind aberrant proteostasis. While affinity-based methods proved powerful for this purpose, direct evidence of modification crucially relied on mass spectrometry (MS)-based analysis^9–13^. Approaches to detect Nt-arginylation by profiling at the protein level often lack positional specifics and direct evidence of modification^11, 12^. To our knowledge, activity-based arginylation profiling which measures mass shifts of Nt-arginylation by ATE1 *in vitro* is the only method that has addressed these challenges^14^.

Identification of Nt-arginylation in a global profiling at the peptide level without any utilization of positional proteomics needs heavy integration of biological materials and separation methods^15^. Positional proteomic techniques such as COFRADIC^15^ and TAILS^16^ have been developed to meet the need to study N-terminal modifications. Such methods would also be suitable for Nt-arginylation since the modification occurs at the N-terminal site of polypeptides after cleavage by intracellular protease or aminopeptidase^15–17^. The identification of Nt-arginylation by only mass shift in MS risks generating false positives because of mass ambiguities associated with other unknown N-terminal modifications and amino acid combinations with similar masses^18^. Recent advancements in machine learning (ML) for predicting MS characteristics, particularly retention time (RT)^19^ and fragment spectra^20^, enhance peptide identification rates and enable assessment of the validity of modified peptides. The integration of ML models into transformer architectures has improved performance and facilitated transfer learning, reducing the size of required training data significantly through the fine-tuning of pre-existing models^21–23^.

Here, we introduce ML for stringent filtering of N-terminomics MS data generated to profile *in vivo* Nt-arginylation. Identifications of Nt-arginylation were assessed by analyzing fragment spectra, RT, and fragment mass errors, comparing them to predicted values obtained using prediction models derived from transfer learning of pre-trained models. The N-terminomics MS data from ER stress-induced HeLa cells were refined to 134 Nt-arginylation sites through false discovery rate (FDR) control and statistical analysis. The Nt-arginylome was further validated using a p62-ZZ domain-derived bait called R-catcher, which has an affinity for arginylated proteins. Temporal changes in the Nt-arginylome following ER stress induction were monitored by parallel reaction monitoring MS (PRM-MS). Our approach significantly enhances understanding of Nt-arginylation substrates by reducing common mis-annotations through the application of ML algorithms.

## Result

### A tandem database search is necessary, but not enough for Nt-arginylation identification

To discover Nt-arginylome by mass spectrometry-based proteomics, we applied our established method, iNrich to human HeLa cells for N-terminal peptide enrichment and LC-MS/MS analysis^24^. The MS data were searched twice sequentially against the human protein database. The first search was designed to remove MS spectra matching peptides with Nt-arginine residues (Arg-starting peptides) that arise when consecutive cleavage sites (for example, -XKRX-in trypsin cleavage) are cleaved at the first site but not the second during sample processing rather than true post-translational arginylation events and the second search on the remaining unassigned spectra for identification of Nt-arginylation as a posttranslational modification (PTM)^25, 26^. Before LC-MS/MS, we treated the cells with an ER-stress inducer, thapsigargin (TG) and a proteasome inhibitor MG132 to enhance *in vivo* Nt-arginylation (Fig. 1a). We identified 392 putative Nt-arginylated peptides and 1,217 corresponding PSMs (Supplementary Data 1). The protein N-termini identified in both consecutive searches showed consistently reproducible counts across replicates and treatments. By contrast, the putative Nt-arginylated peptides yielded varied outcomes depending on TG treatment, demonstrating that Nt-arginylation is specifically responsive to ER stress conditions and consistent with its rarity among protein N-termini (Supplementary Fig. 1). Although many Arg-starting peptides were removed in the first round of tandem database search, we found that many remained in the list as a result of missed proteolytic cleavage rather than Nt-arginylation. The amino acid residues from P3 to P1 (n^th^ residue in protein sequence before the arginylated site) showed high usage of arginine (Supplementary Fig. 2a, b)^27^. For P1=R peptides, logo analysis revealed a preference for lysine and arginine at P2, suggesting potential mis-annotation of missed trypsin cleavage product as Nt-arginylation (Supplementary Fig. 2c). Even after removing 53 peptides with P1=R and P2P1 = GV or VG, which are mass ambiguities of Nt-arginylation, arginine still remained the most prevalent amino acid in positions P3 to P2 (Supplementary Fig. 2d-f)^18^. Nonetheless, we also observed high levels of aspartate in P4 and P1, reminiscent of the caspase cleavage site DXXD motif, suggesting the possibility of genuine Nt-arginylation events^28, 29^. Our results highlight that the Nt-arginylome, as defined by mass deviation only, is influenced by mis-annotations related to various post-translational modifications and amino acid combinations^18^. Thus, a thorough filtering process is required for accurately identifying authentic Nt-arginylated peptides and reducing false positives.

**Figure 1.**
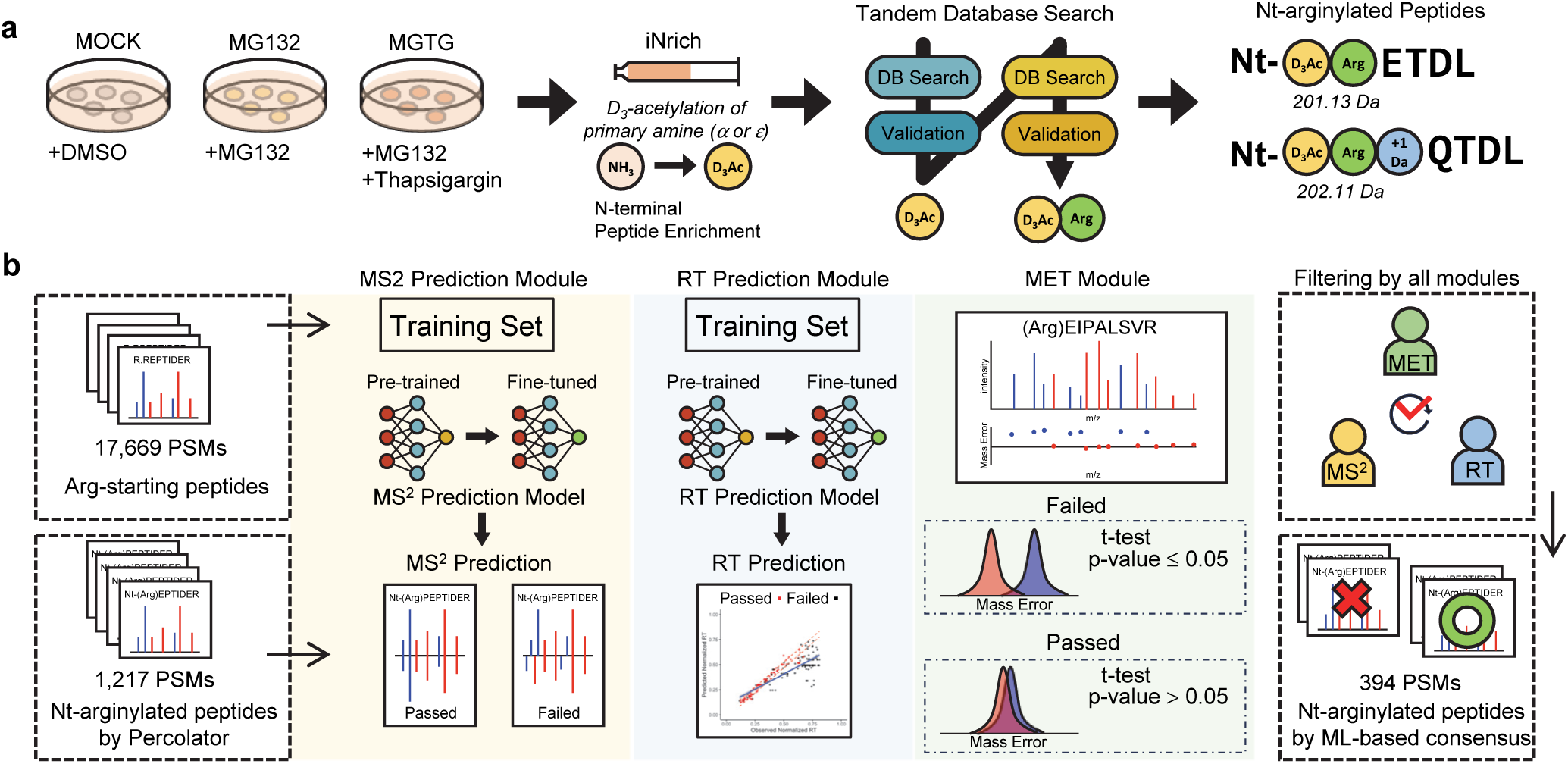
Strategy for the profiling of *in vivo* Nt-arginylation by N-terminomics and ML-based filtering a, Profiling of Nt-arginylome in ER stress-induced HeLa cells. Cells are treated with a proteasome inhibitor (MG132) or MG132 and thapsigargin (MGTG), on which N-terminomics is performed using iNrich method. Mass spectra searched are tried for matching to protein sequences in a two-stage database search method. During the N-terminal peptide enrichment process, peptides with free Nt-amine (α and ε-amine) are labeled with D3-acetyl (D3Ac). **b,** Schematic representation of ML-based filtering of mass spectra for putative Nt-arginylated peptides. Spectra of the Arg-starting peptides derived from missed cleavage of consecutive tryptic sites and their RT were used as training material for the fine-tuning of MS2 prediction model and RT prediction model. Spectra identified as Nt-arginylation in the database search are examined to determine if their fragment spectra or RT align with the predictions made by each model. In the MET filtering step, a null hypothesis that there is no difference in mass error distributions for the two sets of fragment ions, specifically b-ions and y-ions, is examined. The PSMs that have all three ML-based filtering consensuses are further analyzed as high confidence Nt-arginylome.

### Trainable features of Nt-arginylation: peptides containing arginine at the N-terminus generated by missed trypsin cleavage

We hypothesized that Arg-starting peptides produced by missed trypsin cleavage (Supplementary Fig. 3a) could serve as proxies for understanding the characteristics of *in vivo* Nt-arginylated peptides due to their identical chemistry. In this regard, we evaluated the physicochemical characteristics of the Arg-starting peptides, positing that Nt-arginylation would have a pronounced positive effect on peptide hydrophilicity in LC and generation of b-ion fragments in MS/MS, given that the functional group of arginine has the highest pKa among conventional amino acids (Supplementary Fig. 3b). With 17,669 PSMs of Arg-starting peptides obtained from a conventional database search (see details in Method) on the same LC-MS/MS dataset, we observed the following characteristics: i) high b-ion fragment intensities (average area difference in Arg-starting peptides: 25.7%; non-Arg-starting peptides: 45.8%) (Supplementary Fig. 3c, d), ii) relatively low RT distribution than non-Arg-starting peptides (normalized Δt = 0.186) (Supplementary Fig. 3e, Supplementary Data 2). Based on these attributes, a prediction model trained with the mass spectra of Arg-starting peptides is valuable to discover bona fide Nt-arginylome. However, constructing such a model would require at least millions of mass spectra, as suggested by previous research^30^. To resolve this matter, we leveraged a transfer learning strategy with a pre-trained MS2 prediction model, which is part of the recently launched AlphaPeptDeep algorithm, a large language model (LLM) dedicated to proteomics employing transformer layers^23^. By fine-tuning the LLM via transfer learning using the spectra of Arg-starting peptides, we predicted the MS2 spectra and RT of the putative Nt-arginylated peptides identified from the tandem database search, and compared them to the experimental data. In addition, we employed a statistical method based on the mass errors of MS2 fragment ions to verify the accuracy of identification for Nt-arginylated peptides (Fig. 1b, Supplementary Fig. 3f).

### Spectrum filters using ML-based prediction modules and a statistical test

#### MS2 prediction module

To assess the practicality of improving a pre-trained model via transfer learning with LLM, we fine-tuned the pre-trained tryptic peptide MS2 model of the AlphaPeptDeep Python package. The fine-tuning process used all available spectra we acquired (N=311,547 for a trypsin model and N=251,969 for a chymotrypsin model), including 17,669 PSMs of Arg-starting peptides. We also built additional MS2 models by conventional training from scratch or by transfer-learning with the same spectra except for Arg-starting peptides. We then evaluated the performance of each model by calculating Pearson’s correlation coefficient (PCC) between the predicted and the observed fragment intensities (Fig. 2a, Supplementary Data 3) of Arg-starting peptides (N=9,569 in trypsin and N=8,100 in chymotrypsin). From the fine-tuned MS2 prediction model, 87.8% of Arg-starting peptide spectra had a PCC of at least 0.9 (PCC90) while only 56.4% of PCC90 was obtained by the pre-trained MS2 model and 75.9% by “from scratch” model. MS2 prediction performance of the fine-tuned MS2 model is comparable to the result from the reference of AlphaPeptDeep^23^. The other models revealed a strong predictive accuracy for y-ions, but they did not achieve the same results for b-ions. The significance of including Arg-starting peptide spectra is evident, as the fine-tuned model without these peptides failed to exceed the performance of the pre-trained model and even showed reduced predictability for b-ions. On the other hand, a chymotryptic peptide prediction model using a transfer learning of trypsin MS2 model with chymotryptic peptides showed moderately increased prediction performance as PCC90 increased by 9.1% compared to the model without transfer learning (from scratch model) (Supplementary Fig. 4a, b).

**Figure 2.**
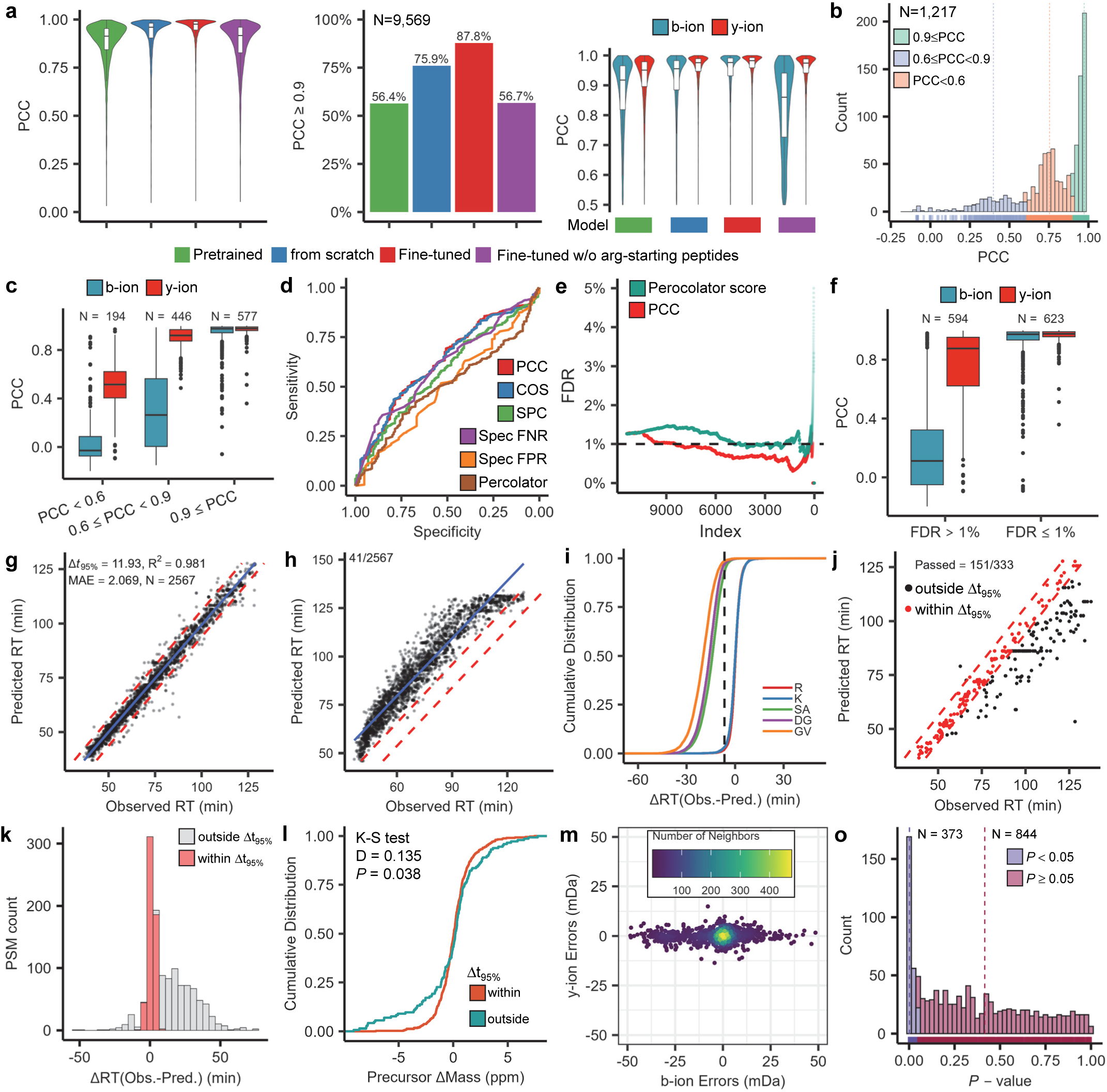
Development of ML-based filtering a, Assessment of prediction accuracy for MS2 spectra with varying training methods. Plotted are the distribution of Pearson’s correlation coefficient (PCC) of PSMs for Arg-starting peptides according to prediction models (left), percentage of PSMs with PCC≥0.9 (center) and fragment ion species-specific PCC (right). PCC is determined through the comparison of fragment ion intensities between the predicted and measured spectra. **b,** PCCs obtained by comparing the PSMs searched as Nt-arginylated peptides to the predicted spectra generated using the fine-tuned MS2 prediction model. Bin size: 0.025. **c,** Boxplots that compare ion species-specific PCC values for b-ion and y-ion according to the PCC score group of Fig 2b. Boxplots show the median, Q1 (25%), Q3 (75%) and whiskers at Q ± 1.5 (Q3–Q1). **d,** Discriminatory power of similarity measures between true positive and false positive Nt-arginylation PSMs using receiver-operating characteristic (ROC) curves. The PSMs were generated using a decoy database specialized for Nt-arginylation search. PCC, Pearson’s correlation coefficient; COS, cosine similarity; SPC, Spearman’s correlation coefficient; spec_FNR, spectral false negative rate; spec_FPR, spectral false positive rate; percolator, percolator score. **e,** Cumulative false discovery rate by order of percolator score and PCC similarity score. A PSM with a lower index has a higher score. **f,** Distribution of ion species specific PCC values for b-ion and y-ion divided into two groups based on the PCC score of Fig 2e corresponding to 1% FDR. Boxplots show the median, Q1 (25%), Q3 (75%) and whiskers at Q ± 1.5 (Q3–Q1). **g,** RT of Arg-starting peptides predicted by RT prediction model versus observed RT. Blue line, fitted linear regression; red dashed line, Δt_95%_ region. **h,** Predicted RT of peptides with Arg replaced by Gly-Val in Arg-starting peptides of Fig 2g. Only 41 out of 2,567 PSMs remain within the Δt_95%_ RT interval. **i**, Deviation between the predicted and observed RT of Arg-starting peptides in which Arg is replaced by other types of amino acid/dipeptides. The distributions are expressed as cumulative fractions. Dashed line, RT at Δt_95%_. **j,** Predicted versus observed RT of the putative Nt-arginylated peptides obtained by tandem database search. Red dashed lines indicate the Δt_95%_ region of the RT model. **k,** A histogram of Nt-arginylation peptides as a function of RT deviation between observed and predicted. **l,** Distribution of precursor mass errors that fit (within) and do not fit (outside) the RT model. A two-sample Kolmogorov– Smirnov (K–S) test shows that the two distributions are different. D, distance statistic; P, *P*-value. **m,** Average mass errors of b- and y-fragment ions in each PSMs of the putative Nt-arginylated peptides. **o,** A histogram of Nt-arginylation peptides as a function of *P*-values obtained from Student’s t-test comparing b-ion errors and y-ion errors. The PSMs are divided into two groups based on *P*-values. Dashed line, median of each group.

To evaluate the predicted spectra using the fine-tuned MS2 model, we compared the measured mass spectra of the well-known Nt-arginylation proteins with the predicted spectra, i.e., CALR|18, P4HB|18, and FBLN1|30 (denoted as gene name|arginylation site). The prediction accuracy was significantly high with average PCC of 0.954±0.117 for the CALR|18 (36 PSMs), 0.931±0.081 for the P4HB|18 (37 PSMs), and 0.960±0.038 for the FBLN1|30 (12 PSMs), while the PCC values were 0.891±0.121, 0.915±0.094, and 0.938±0.061, respectively, when predicted with pre-trained model (Supplementary Fig. 4c-e, Supplementary Data 1). The median PCC was 0.878 for all 1,217 PSMs including 15 chymotryptic PSMs of putative Nt-arginylated peptides (Supplementary Fig. 4f). We categorized the prediction results into three groups based on the PCC values: high (PCC ≥ 0.9), moderate (0.9 > PCC ≥ 0.6) and low (PCC < 0.6) (Fig. 2b). In the high group, the median PCC of b-ions and y-ions were 0.976 and 0.978, respectively, compared to the moderate group with 0.264 and 0.920 and the low group with 0.051 and 0.510 for b-ions and y-ions, respectively (Fig. 2c). The gathered data illustrate that database search alone relies primarily on y-ions of tryptic peptides, while b-ion signals are required for sufficient confirmation of Nt-arginylation.

Next, we aimed to determine the cut-off for the PCC score for maximizing sensitivity and specificity in identifying authentic Nt-arginylated peptides. In pursuit of this goal, we constructed a decoy database by altering the protein sequences, wherein consecutive arginine residues were consolidated into singular arginine (Supplementary Fig. 5a). The application of the decoy database causes Arg-starting peptides to become Nt-arginylated peptides and thus allows for the distinction of true positives from false positives by determining whether the peptides are originated from the altered location of the decoy database or not. With the decoy database, 11,407 PSMs containing Nt-arginylation modification were identified, with 144 of them being deemed as false (Supplementary Data 4). There was no significant difference (*p*-value = 0.94) in the Percolator scores of the database search output between the true and false positives (Supplementary Fig. 5b). ROC analysis indicated that the Percolator method had unreliable performance, with an AUROC value of 0.498. In contrast, PCC values of the MS2 prediction-based rescoring method demonstrated improved performance, achieving an AUROC of 0.624 (Fig. 2d, Supplementary Fig. 5c). Several other metrics reflecting the similarity between predicted and observed spectra also showed higher AUC values compared to the Percolator score, with PCC achieving the highest performance. PCC outperformed the Percolator score in controlling the FDR. The cumulative FDR obtained by sorting the Percolator scores was always higher than the FDR obtained by PCC scores, and the deviation was particularly large in spectra with high scores. (Fig. 2e). We chose a cut-off value of 0.869 for PCC in the trypsin dataset, which achieved a cumulative FDR of 1%. The median PCC for b-ions and y-ions were 0.973 and 0.976, respectively (Fig. 2f). For the chymotrypsin dataset, we set a PCC threshold at 0.9, a decision driven by the limited number of 15 PSMs associated with Nt-arginylated peptides. Using this PCC cutoff, 623 out of 1,217 PSMs of Nt-arginylated peptides were accepted.

#### RT prediction module

Arg-starting peptides exhibited another distinct feature, earlier elution than ordinary tryptic peptides during reverse-phased LC (Supplementary Fig. 3e). For all sequential LC-MS/MS runs performed on fractionated samples originating from a single sample, a total of 12 RT models were generated, one per run^23^. Comparison of the observed RT and the predicted RT for Arg-starting peptides in the 12 fine-tuned RT models showed a significant level of predictive accuracy with R^2^ values spanning from a low of 0.955 to a high of 0.980. (Fig. 2g, Supplementary Fig. 6). We used these RT models as filters to determine the absence or presence of arginylation modification of the putative Nt-arginylated peptides.

We first evaluated the RT models by comparing the RT changes when the Nt-modification of Arg-starting peptides was replaced with GV, a dipeptide that is identical in mass to Nt-arginylation but is not basic^18^. The average rise in normalized RT upon GV substitution was 0.105, which is more than the average value of 0.0475 for Δ*t*_95%_, a 95% confidence level derived from the linear model (Fig. 2h). Only 232 out of 17,669 PSMs within 95% confidence of prediction (Supplementary Fig. 7). In addition to GV, the RT model reliably distinguished other substitutions, such as SA and DG, except lysine (K) which exhibits basicity like arginine (Fig. 2i). This demonstrates that the RT models developed through transfer learning of Arg-starting peptides can effectively distinguish Nt-arginylation modification from hydrophobic or neutral modifications with similar masses.

Based on the results showing that the RT model generated from Arg-starting peptides accurately predicted the elution times of Nt-arginylated peptides, we established a filtering criterion at the 95% confidence level (Δ*t*_95%_) of the linear regression (Supplementary Fig. 8). Nt-arginylated peptides falling within the prediction interval were classified as positives (Fig. 2j). Among 1,217 PSMs, 544 passed the filter. Intriguingly, most PSMs outside the 95% confidence interval exhibited greater-than-predicted increases in RT (Fig. 2k) suggesting that unknown modifications initially mistaken for Nt-arginylation during the database search are less basic than arginine (Supplementary Fig. 3e) which is a plausible outcome. Moreover, the PSMs that passed the RT filter had smaller mass deviation than those that did not pass (Fig. 2l), suggesting that the RT prediction module worked appropriately.

#### Mass error test (MET) module

We introduced an additional evaluation module that leverages systematic variations in intrinsic mass measurement inaccuracies caused by mis-annotations^31^. This module is based on the hypothesis that mass spectrometers produce equivalent measurement errors for fragment ions regardless of whether the ion is a b-ion or a y-ion. However, when we compared the error distributions between b-ions and y-ions across 1,217 putative Nt-arginylated PSMs, the average error for b-ions was -1.37±14.2 mDa, while for y-ions, it was 0.0383±8.93 mDa (Supplementary Fig. 9a, b). This imbalance between b-ion and y-ion errors was consistent at the spectrum level (Fig. 2m). These findings suggest that b-ion mass errors likely stemmed from mis-annotation of the N-terminal modification. Indeed, we have proposed that the observed error discrepancy could serve as a confidence metric for assessing the correctness of N-terminal modifications, which we termed the MET^32^.

We used a two-tailed Student’s t-test to compare the b-ion error distribution with the y-ion error distribution, and discarded spectra with *P*-values less than 0.05 (Fig. 2o). MS2 and RT prediction modules could also remove several spectra with heterogeneous mass errors (Supplementary Fig. 9c, d). Nevertheless, we observed significant discrepancies between b- and y-ion mass error distributions in many PSMs having agreement with the MS2 and RT prediction modules (Supplementary Fig. 9e-h). These findings suggest that MET can effectively eliminate mis-annotations of other modifications with physicochemical properties similar to Nt-arginylation. Using the MET module, we narrowed down the 1,217 PSMs to 844.

#### Performance of ML-based filtering

Of 1,217 PSMs identified as Nt-arginylated in database search, 394 PSMs were retained after being processed by ML-based filtering (Fig. 3a). The proportion of remaining PSMs after filtering under each experimental condition varied across the three conditions. In DMSO treated control (MOCK), only 16.1% PSMs remained (Fig. 3b) while 50.8% for MG132 treated samples (MG132) and 65.3% for MG132 and TG treated samples (MGTG) were kept. In contrast, the number of discarded PSMs was similar: 252, 246, and 317 PSMs from MOCK, MG123, and MGTG, respectively. This indicates that the results of ML-based filtering reflect the biological disturbances we intentionally introduced and remove randomly appearing false positives.

**Figure 3.**
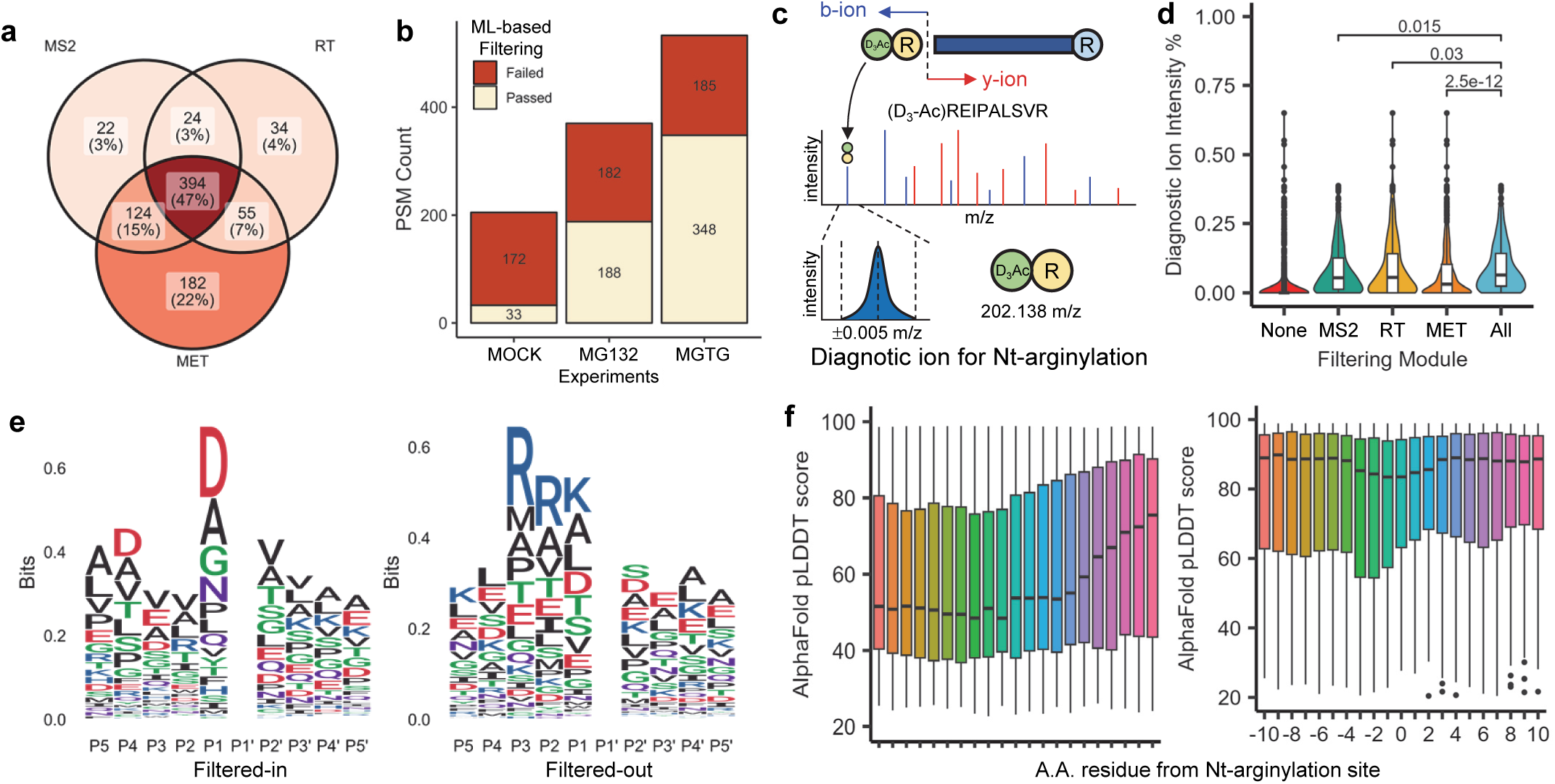
Validation of ML-based filtering a, A Venn diagram showing the PSMs of Nt-arginylated peptides that pass each ML-based filtering module. **b,** Number of PSMs for Nt-arginylated peptides that pass all three ML-based filtering modules. **c,** Schematic for annotating the diagnostic ion of Nt-arginylated peptide. **d,** Proportion of the diagnostic ion intensity compared to the total ion intensity. The x-axis represents the PSMs that pass the indicated prediction module. Two-tailed Student’s t-test *P*-values are shown. **e,** A logo plot showing Nt-arginylation sites that pass (Filtered-in) or fail to pass (Filtered-out) the ML-based filters. The logo for P1’, which is made of only four amino acids, D/E/N/Q, is not displayed. **f,** The pLDDT score distributions determined by AlphaFold for the peptide residue positions of Nt-arginylation sites that pass (left) and fail (right) ML-based filtering. The position value of 0 denotes the residue where Nt-arginylation occurs.

The effectiveness of ML-based filtering could also be verified by frequent observation of fragment ion of the modification itself. When comparing MS2 spectra of an Arg-starting peptide with that of the same peptide without Arg, the b1-ion of D_3_-acetylated Arg is generally observed at 202.138±0.005 m/z (Fig. 3c and Supplementary Fig. 10a, b). The b1 ion was observed in 15,665 out of 17,669 PSMs of Arg-starting peptide at the 1% intensity cutoff and their median relative intensity was 16.9% in trypsin and 11.7% in chymotrypsin datasets (Supplementary Fig. 10c). Chi-squared test showed that the Arg-starting peptide were significantly enriched with b1 ion (*P*-value <2.2×10^-16^) (Supplementary Fig. 10d). The b1-like diagnostic ion would also be observed at the same m/z in the MS2 spectrum of Nt-arginylated peptide. The 394 PSMs that passed all three modules had higher median relative intensity of diagnostic ion (6.48%) than any other PSM groups that failed at least one of the three modules. The PSMs that failed in all modules had the lowest median, 0% (Fig. 3d). This suggests that not only the MS2 module but also two other modules select PSMs of peptides with similar properties to Arg-starting peptides and the synergy of all three modules strengthens this tendency.

Next, we revisited sequence preference of P5-P5’ sites of Nt-arginylation after ML-based filtering. Comparing the logo analysis results before and after filtering, PSMs with arginine at P3 and P2 sites were mostly removed (Fig. 3e). Instead, aspartic acids were observed frequently at P4 and P1 sites which is analogous to the DXXD caspase motifs^28^. In concordance, DAU analysis also showed that arginine is categorized as one of the least preferable residues in P5-P1 sites (Supplementary Fig. 11).

A previous research has demonstrated that the structural context of functionally relevant PTMs is concentrated within the intrinsically disordered region when assessing the predicted protein structure^33^. Remarkably, structure prediction scores from AlphaFold database on Nt-arginylation sites of 394 PSMs showed low average pLDDT (58.5 ± 23.0) while the sites of PSMs excluded by ML-based filtering exhibited high average pLDDT (75.1±21.0) with significant difference (*P* = 1.1×10^-11^) (Fig. 3f, Supplementary Data 1). The result is consistent with the notion that PTMs occurs in the disordered region of proteins. Notably, the pLDDT scores of the residues in the N-terminal direction from Nt-arginylation sites remained similarly low, which is not observed in the C-terminal direction. Taken together, the comparative analyses show that ML-based filtering is effective for screening mass spectra with Nt-arginylation modification.

### Characterization of ER-stress induced Nt-arginylome

From 394 Nt-arginylation PSMs that passed ML-based filters, 134 Nt-arginylation sites were annotated (Fig. 4a, Supplementary Table 1, Supplementary Data 5). In order to comprehend the biological roles of annotated Nt-arginylome in response to ER stress, we conducted Reactome pathway enrichment analysis using the list of Nt-arginylated proteins (Supplementary Data 6). The UPR pathway stood out as the most significantly enriched pathway (Fig. 4b). The impact of TG was evident as the gene set enrichment analysis (GSEA) revealed distinct pathways connected to protein metabolism and stress responses in the comparison of Nt-arginylation sites between MGTG and MG132 (Supplementary Fig. 12a). Gene ontology (GO) overrepresentation analysis (ORA) showed that the highest enriched ontologies were GO terms related to cellular component (GOCC), such as focal adhesion, cell-substrate junction, and ER lumen (Supplementary Fig. 12b). A supplementary analysis with GOCC terms reveals that proteins with Nt-arginylation are found in a broad spectrum of subcellular organelles, including ER, nucleus and mitochondria (Supplementary Fig. 12c, d). The protein-protein interaction analysis using STRING database revealed that the arginylome proteins are functionally closely connected with each other even though they are located in various subcellular organelles such as nucleus, cytoplasm, extracellular space, and endoplasmic reticulum (Supplementary Fig. 12e). MCL clustering analysis reveals that essential cytoskeletal proteins, such as beta actin (ACTB), filamin A (FLNA) and moesin (MSN), are at the middle of proteins of ER lumen and cytoplasmic ribonucleoprotein granule (Supplementary Fig. 12f). This suggests a homeostatic role of ATE1, as previously demonstrated by the interaction of Nt-arginylated calreticulin with stress granules^34^. Of note, the observation of Nt-arginylated proteins in MOCK hints a mechanism where Nt-arginylation may not induce protein degradation. These Nt-arginylated proteins also remained stable in MG132 and MGTG (Supplementary Fig. 13).

**Figure 4.**
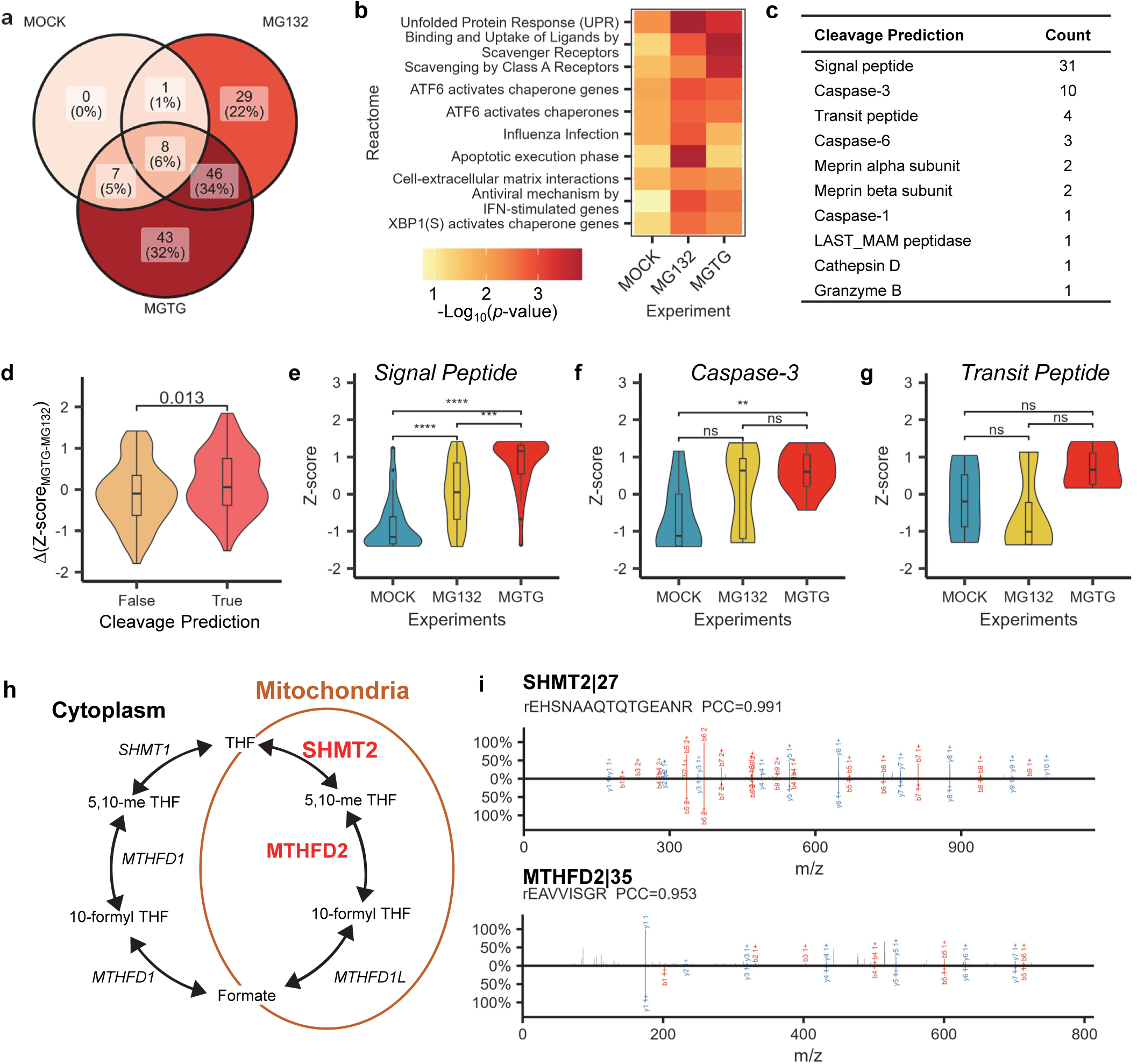
Characteristics of the Nt-arginylome under ER stress a, A Venn diagram showing the proportion of Nt-arginylation sites that were identified in each experimental condition. **b,** Reactome pathway enrichment analysis of the Nt-arginylated proteins. **c,** Number of Nt-arginylation sites with the motif of specified proteases. Cleavage sites were derived from the predictive output of SignalP, TransitP and Procleave. **d,** Quantitative comparison of Nt-arginylation sites, categorized by the absence or presence of a predicted cleavage site motif, with Δ(Z-score) obtained by comparing MGTG and MG132. Z-scores for each experiment were obtained from the MS1 label-free quantification result of all identified peptides regardless of Nt-arginylation. **e-g,** Quantitative distribution of Nt-arginylation sites predicted as cleavage sites for signal peptides (**e**), caspase-3 substrates (**f**), and transit peptides (**g**). Z scores were obtained as in Fig. 4d. T-test *P*-values are: \*\**P* <0.01, \*\*\**P*<0.001, \*\*\*\**P*<0.0001. ns, not significant. **h,** One-carbon metabolism pathway. The proteins marked in red are those where Nt-arginylation sites at the transit peptide cleavage site were discovered. **i**, Spectra mirror plots for the Nt-arginylation sites of Fig. 4h. The observed mass spectrum is shown in the upper plane and the corresponding predicted spectrum is in the bottom plane. Nt-arginylation is indicated by ‘r’ at the beginning of the peptide sequence and the protein name is denoted as (gene name)|(Nt-arginylation site position).

The preference of Nt-arginylation sites for less ordered structure can be elucidated by their explicit association with protease cleavage sites^4^. With signal^35^, transit^36^ and protease site predictions^37^, we have found 56 Nt-arginylation sites that displayed a high likelihood of being protease cleavage sites (Fig. 4c). The most abundant cleavage sites we observed were signal peptide, transit peptide, and caspase-3 substrates. Logo analysis on these sites indicated that the presence of arginine in the P2 position is strongly associated with signal peptides and transit peptides (Supplementary Fig. 14a, b), which is mostly in agreement with the previously reported amino acid frequency of signal and transit peptides^36^. The results for Nt-arginylation sites lacking known protease motifs or predicted to be caspase-3 substrates showed no significant presence of arginine (Supplementary Fig. 14c, d). Importantly, we had previously hypothesized that Nt-arginylation sites having arginine at P2 were from mis-annotations. Nevertheless, our data unequivocally demonstrate clear link between Nt-arginylation and protease cleavage.

Quantitation of the Nt-arginylome demonstrates increased level of Nt-arginylation at sites cleaved by proteases upon MGTG treatment (Fig. 4d). It was evident that arginylation increased at the N-terminal site newly exposed after the signal peptide is cleaved off, or at the N-terminal region newly exposed in the substrate following caspase-3 cleavage. (Fig. 4e, f). Mitochondrial proteins containing transit peptides also appeared to increase, although this was not statistically significant (Fig. 4g). Even when all mitochondrial proteins were included, the increase was not as pronounced as with other organelle proteins (Supplementary Fig. 15). We identified six Nt-arginylated mitochondrial proteins, four of which were at the transit peptide cleavage site. Interestingly, two proteins SHMT2|27 and MTHFD2|36 out of the four are primarily involved in folate pathway, known as one-carbon metabolism (Fig. 4h, i)^38^. Since these proteins are essential for cancer cell survival and proliferation, the discovery of Nt-arginylation of these proteins may provide a novel tool for controlling these potential therapeutic targets^39^.

### Validation of Nt-arginylome using R-Catcher pulldown assay

Nt-arginylation of the identified arginylated proteins were validated using an R-catcher pulldown assay^11, 25^. The R-catcher is derived from the ZZ domain of p62/sequestosome-1, known to have binding affinity toward arginylated protein/peptide. We expected that at least some, if not all, Nt-arginylated proteins would have binding affinity to R-Catcher, Therefore, we arbitrarily selected 12 proteins out of the Nt-arginylome list, cloned and transfected the corresponding genes into HeLa cells. After treating cells with MG132 and TG (MGTG), pull-down assays were performed using R-catcher beads. The results were visualized by western blot (Fig. 5a). As a control, we also used a D129A mutant R-catcher that lacks binding affinity for arginylated peptides. Eight out of the 12 candidate proteins exhibited significant and selective binding to the wild-type R-catcher (Fig. 5b, c). The same phenomenon was observed when both Nt-arginylated HSPA5 and CALR, which are known to bind to p62/sequestosome-1 upon arginylation, were used as positive controls, proving proper working of the assay. We then performed competition assays by pre-charging the R-catcher with an RA dipeptide. All proteins except PDIA3 and CALU lost binding affinity for the R-catcher in the presence of the RA dipeptide but not in the presence of the control AR dipeptide, confirming the specificity of the interaction (Fig. 5d). Nt-arginlyation of these 6 proteins by ATE1 was confirmed by comparing the R-Catcher pulldown assays performed using ATE1 wild-type (ATE1+/+) and knockout (ATE1-/-) MEF cells. The interaction was seen only in wild-type MEF cells not in ATE1 KO cells (Fig. 5e). Taken together, our results suggest that arginylated proteins discovered through MS are substrates of ATE1-mediated arginylation *in vivo*. Furthermore, our *in vitro* R-catcher pull-down assay demonstrates that a significant subset of these proteins can interact with p62, suggesting their potential involvement in autophagy pathways following arginylation. Additionally, western blot validation confirms that our ML-based filtering approach effectively enhances the identification accuracy of N-terminally arginylated peptide spectra.

**Figure 5.**
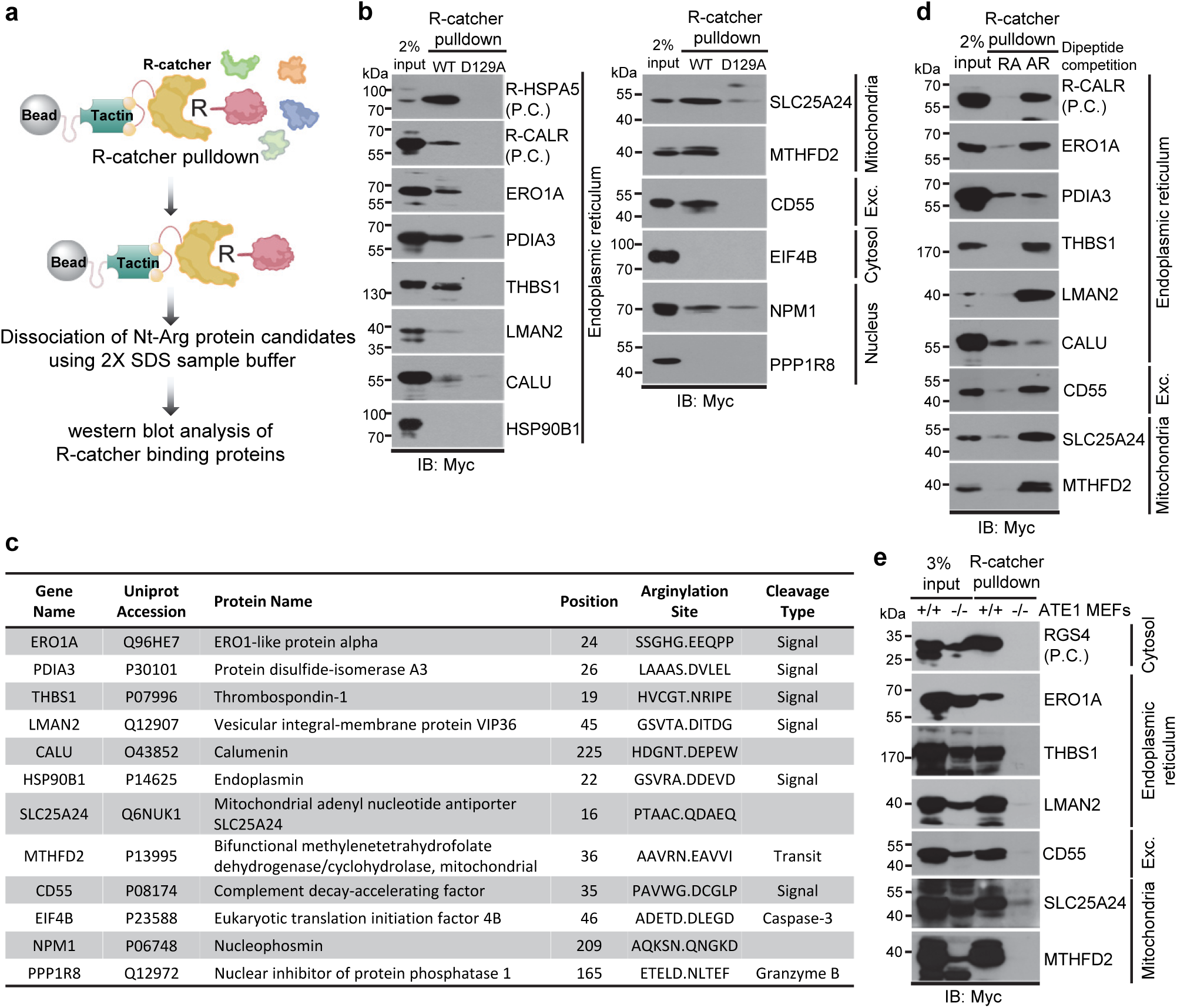
Validation of Nt-arginylation candidate proteins using R-catcher pulldown assay a, A schematic representation describing the R-catcher pulldown assay for validation of Nt-arginylation candidate proteins. **b,** R-catcher pulldown assay of Nt-arginylation candidate proteins in HeLa cells treated with MG132 10 μM plus thapsigargin 50 nM. HSPA5 and CALR are positive controls showing that the assay system works properly. **c,** List of proteins tested for R-catcher pulldown assay. Position: residue number in the protein sequence where Nt-arginylation occurs; Arginylation site: 10 amino acid sequence surrounding the Nt-arginylation site, spanning 5 residues on each side; Cleavage type: proteases specified by the motif of Nt-arginylation sites. **d,** Dipeptide competition R-catcher pulldown assay to validate the binding is Nt-arginylation-dependent. HeLa cell lysates stimulated with MGTG were incubated with a purified R-catcher in the presence of 25 mM dipeptides, RA or AR. **e,** R-catcher pulldown assay to validate ATE1-dependent arginylation of candidate proteins using ATE1 +/+, -/- mouse embryonic fibroblasts (MEFs) cell. P.C.: positive control.

### Temporal changes of Nt-arginylation in response to ER stress

We next monitored temporal changes of Nt-arginylation in response to ER stress. After treating HeLa cells with MGTG, arginylated proteins and the corresponding unmodified proteins were detected by western blot and MS at regular intervals. Antibodies for studying PTMs are generally limited, and even for Nt-arginylated proteins, only a few antibodies were available. For proteins HSPA5, CALR and P4HB, we used antibodies directed to the arginylated form. We attempted PRM-MS to detect Nt-arginylated proteins whose antibodies were not available (Fig. 6a). We aimed to monitor 21 Nt-arginylation sites with their corresponding unmodified sites primarily associated with UPR-related biological processes, apoptosis, autophagy, and caspase cascade (Supplementary Data 7). In addition, regardless of arginylation, 6 peptides derived from 6 proteins that have previously served as markers for the UPR and the advancement of subsequent processes were included. Fourteen synthetic peptides (ISTDs) exhibiting a broad range of RTs were utilized as standards to evaluate system performance.

**Figure 6.**
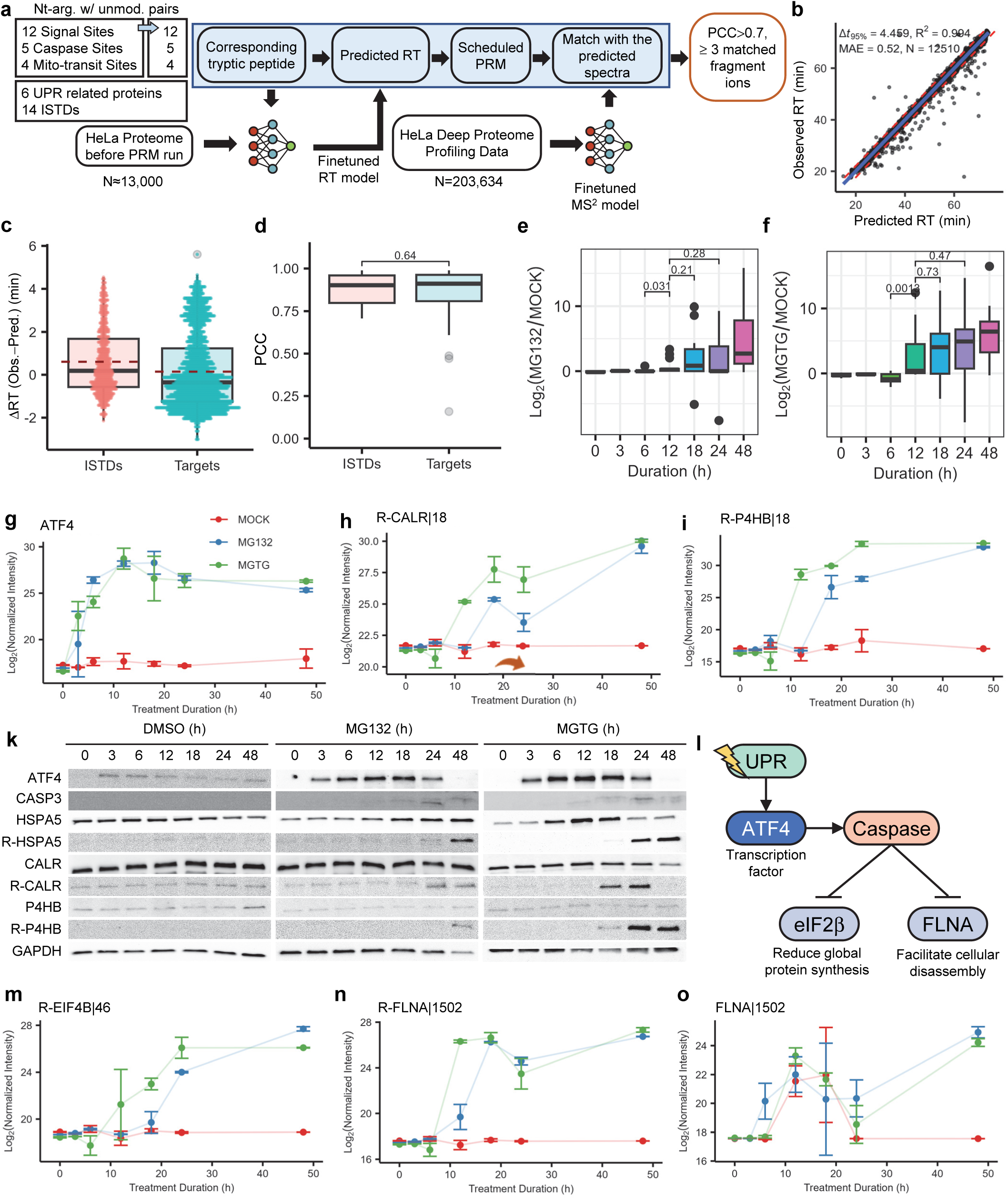
Temporal change in Nt-arginylation during ER stress monitored by parallel reaction monitoring mass spectrometry a, A schematic representation of parallel reaction monitoring mass spectrometry (PRM-MS) performed in this study. For each Nt-arginylation target (Nt-arg.), the corresponding unmodified (unmod.) peptides were also monitored. **b,** Observed versus predicted RT of tryptic peptides of HeLa digest. **c-d,** Box plots comparing retention time deviation (**c**) and PCC similarity value (**d**) between predicted and observed MS spectra for synthetic ISTD peptides and target peptides. **e-f,** Log_2_ fold changes of N-arginylation sites comparing MG132 to MOCK (**e**) and MGTG to MOCK (**f**). A two-tailed Student’s t-test was used to analyze the results. **g-i,** Normalized intensities of target peptides as a function of drug treatment duration. Shown are quantitation results for a tryptic peptide of ATF4 (**g**), Nt-arginylated peptide starting at the 18^th^ residue of CALR (**h**), and Nt-arginylated peptide starting at the 18^th^ residue of P4HB (**i**). **k**, Immunoblot analysis of ATF4, activated CASP3, HSPA5, CALR and P4HB during ER stress. R-HSPA5, R-CALR, and R-P4HB represent Nt-arginylated forms of HSPA5, CALR and P4HB, respectively. **l**, Schematics of UPR leading into caspase-3 activation and cleavage of its substrates. **m-o**, Normalized intensities of substrate peptides at the caspase cleavage sites as a function of drug treatment duration. EIF4B|46 with Nt-arginylation (**m**), FLNA|15 02 with Nt-arginylation (**n**), and FLNA|1502 without Nt-arginylation (**o**).

In PRM-MS, whole cell lysate was digested with trypsin and analyzed in a single run of LC-MS acquiring only the MS2 spectra of the target peptides repeatedly in a predefined time duration. Since the N-terminal peptide was not enriched beforehand, the peptide we monitored was not the same as that we found in the profiling experiment: lysine was left unacetylated thus susceptible to tryptic cleavage. For example, if Arg-DEPEWVKTER is the peptide detected in the profiling experiment., Arg-DEPEWVK was attempted to be monitored in the PRM-MS. For this, we had to predict theoretical MS2 spectra and RT for the new peptides by ML. Tryptic digests of HeLa cell lysate were analyzed shortly before the PRM experiment with the same LC condition, and the peptide-RT data matrix was fine-tuned by transfer-learning. After testing several gradient conditions, we selected a 24%B gradient that allowed even distribution of multiple target peptides across the entire LC running time (Supplementary Fig. 16). The fine-tuned RT prediction model gave R^2^ of 0.994 and an MAE (mean absolute error) of 0.52 min (Fig. 6b, Supplementary Data 8). To validate PRM spectra, a fine-tuned MS2 model using 203,634 PSMs of HeLa tryptic peptides was prepared (Fig. 6a). The performance of the fine-tuned MS2 model was measured as an average PCC of 0.957 (Supplementary Fig. 17). We further optimized PRM parameters, i.e., acquisition time and isolation width (Supplementary Figs. 18 and 19).

We then performed 42 PRM-MS runs, monitoring the target peptides and ISTDs in duplicated samples of three different conditions, MGTG, MG and Mock, collected at seven-time points from 0 to 48 hours after ER stress activation. Each PRM-MS run was evaluated with the 14 ISTDs included therein, which maintained ΔRT of no more than 4 min to allow 6 min of monitoring time duration around the predicted RT. The average ΔRT and MS2 PCC of ISTDs were 0.610±1.537 min and 0.876±0.098, respectively, across all 42 PRM-MS runs (Supplementary Fig. 20a, b). The values for the target peptides were 0.144±1.738 min and 0.854±0.166, respectively (Fig. 6c). No significant difference was observed between ISTDs and the targets in the MS2 similarity score (Fig. 6d). For further analysis, we chose only the MS2 scans with cosine similarity>0.7 (ref. ^40^) and utilized the three fragments with the highest intensity for quantification (Supplementary Fig. 20c).

Of 21 Nt-arginylation sites and 21 unmodified sites, 15 arginylated and 11 unmodified sites could be detected (Supplementary Fig. 21, Supplementary Data 9). The PRM-MS result displayed a notable increase of Nt-arginylation in almost all sites upon MG132 and MGTG treatments, with MGTG showing a more pronounced increase than MG132 in line with the predictions made from LFQ analysis of Nt-arginylome (Fig. 6e, f, Supplementary Fig. 22 and 23). By contrast, the corresponding unmodified sites were insensitive to induced ER stress (Supplementary Fig. 24). The increase in arginylation began later than the increase in the amount of ATF4 protein, a UPR transcription factor^41^. ATF4 demonstrated its initial rise at 3 hours post-treatment (Fig. 6g) while arginylation of CALR|18 and P4HB|18 increased at 12 hours post-treatment in MGTG and 18 hours in MG132 (Fig. 6h, i). Immunoblot analysis using the antibodies to Nt-arginylated CALR and Nt-arginylated P4HB showed similar temporal change to those observed by PRM-MS (Fig. 6k). Furthermore, HSPA5, a well-known substrate of ATE1^42^, also showed an increase in Nt-arginylation at a similar time period, indicating that our experimental condition was appropriate to profile ER stress-induced arginylome.

Other notable Nt-arginylation sites were those located at predicted caspase cleavage sites. We monitored four sites by PRM-MS, two of which are in EIF4B^43^ and FLNA^44^. These two sites (EIF4B|46 and FLNA|1502) have previously been shown to be cleaved by caspases but this is the first time we have seen them following UPR induction (Fig. 6l). The induction of UPR led to a notable rise in Nt-arginylation after 12 hours in both MG132 and MGTG treatments relative to MOCK and stabilizing at 12 hours in the MGTG condition and this increase preceded the alteration seen at signal or transit peptide cleavage sites (Fig. 6m, n, Supplementary Fig. 24). Immunoblotting using cleaved caspase-3-specific antibody demonstrated that these temporal changes were consistent with caspase-3 activation (Fig. 6k). In contrast, the unmodified form of FNLA|1502 was detected in small amounts making the change unclear. This implies that FLNA is immediately Nt-arginylated once cleaved by caspase-3 (Fig. 6o). The PRM-MS analysis results confirmed that the targets of caspase-3 are indeed subject to modification via Nt-arginylation in the context of the UPR.

## Discussion

Recent advancements of proteomic technologies reach significant success on even a scale of a single cell. Yet, PTM proteomics, like phosphorylation, needs signal-boosting techniques such as PTM enrichment^45^ and refence boost^46^. Thus, PTM studies are often resource-intensive and laborious. Researching Nt-arginylation serves as an additional illustration of the constraints in studying PTMs. Due to its lower prevalence compared to modifications such as acetylation and phosphorylation and lack of Nt-arginylation specific affinity purification method, the detection of Nt-arginylation leads to numerous incorrect identifications stemming from the weak signals in fragment spectra and poor reproducibility. It is evident from our findings that a mere 32% of Nt-arginylation PSMs met the criteria set by our rigorous filtering strategy. PTM proteomics studies, because of this, need the pursuit of strict specificity which can often lead to reduced sensitivity^47^. Our method introduces more flexible thresholds that preserve specificity yet enhance overall sensitivity. These new stringent filtering criteria for terminal PTMs will greatly enable future studies, allowing the community to not only address the complexity of Arg/N-degron pathway but also survey other terminal PTMs, such as methylation, lipidation, and ubiquitination.

Our study focused on the identification of Nt-arginylation sites, which exhibit a significant inclination towards positive charges, resulting in a higher likelihood of ionization and detection during MS analysis. These characteristics, when utilized alongside N-terminal peptide enrichment, have the potential to generate considerable synergistic benefits for constructing ML-based filtering modules. The N-terminomics method we applied involving blocking primary amines resulted in N-terminal peptides having positive charges only at their C-termini. Consequently, fragment spectra should exhibit predominantly y-ion species in LC-MS analysis. In contrast, fragment spectra from Nt-arginylated peptide have distinguishable intensities of b-ion fragments. We have shown that this substantial discrepancy between Nt-arginylated and non-arginylated peptides greatly enhances the performance of the fragment spectrum prediction model. Similarly, this charge deprivation at the N-terminal amine also creates advantages in setting up the RT prediction model. Indeed, many peptides that were searched as Nt-arginylated peptides but discordant to the RT model tended to have positive RT bias suggesting a mis-annotation from more hydrophobic modifications or amino acid combinations. The enrichment method we used is not to enrich arginylated peptides, but to enrich N-terminal peptides and then analyze them by MS to find Nt-arginylated peptides among them. As a stark difference between the peptide with modification-of-interest like Nt-arginylation and background peptides aids in establishing distinct exclusion criteria, we envision that an imperfect enrichment method like iNrich that generates background peptides with contrasting characteristics for modification-of-interest will provide clearer criteria to gain genuine targets.

We compared our Nt-arginylome with that recently discovered by Lin and colleagues using an ATE1-dependent reaction system reconstituted *in vitro*^14^. A key contrast between the two studies is that the Nt-arginylation sites we have discovered are indicative of the *in vivo* context. Our methodology enables us to focus on investigating the *in vivo* effects of ATE1 under specific stress conditions, allowing us to pinpoint Nt-arginylation sites associated with protease substrates that were degraded or not activated in the control group. Among the 229 Nt-arginylation sites discovered by Lin et al and the 134 Nt-arginylation sites discovered in this study, 10 sites were discovered in both. Even excluding the stress-induced MGTG results, there were seven overlaps (Supplementary Fig. 26a-b). In terms of protein, 10 proteins were shared between 119 proteins from our study and 161 proteins from Lin et al (Supplementary Fig. 26c). Interestingly, 5 of 10 shared sites are originated from signal peptide cleavage and 4 are from mitochondrial transit peptide cleavage (Supplementary Fig. 26d). The derivation of common Nt-arginylation sites from their natural biological state provides insight into the discrepancies caused by methodological differences. We detected five Nt-arginylation sites that other studies have confirmed as substrates for caspases whereas the sites reported by Lin et al. did not (Supplementary Fig. 26e)^48^. The caspase substrates that underwent Nt-arginylation exhibited elevated levels following treatment with thapsigargin, which trigger stress related to unfolded proteins. Notably, all caspase substrates, except β-actin (ACTB|12), were absent in the results from the MOCK treatment. Since there is a crucial connection between Nt-arginylation and caspase substrates, our method delivers a detailed insight into the role of *in vivo* Nt-arginylation ^49^.

Nt-arginylome we found in this study represents substrates of ATE1 *in vivo*. Investigation of proteins that are potential substrates of the pathway involving cleavage and degradation could be facilitated by *in vivo* method highlighted by Nt-arginylation sites of caspase cleavage sites. In particular, Nt-arginylation of mitochondrial proteins with cleaved transit peptides were key enzymes in mitochondrial folate metabolism and revealed the evidence for the connections between UPR and folate metabolism. A research has shown that the control of translation mediated by the UPR leads to the enhanced expression of enzymes involved in mitochondrial folate metabolism^50^. The conflicting relationship between the involvement of mitochondrial folate metabolism and the predominantly degradative function of Nt-arginylation, a key ligand in the Arg/N-degron pathway, remains perplexing. A recent study demonstrated the mechanism behind the autophagic degradation of cytosolic mitochondrial DNA is significantly associated with ATE1^51^. We speculate that targeting of ATE1 towards mitochondrial enzymes is associated with the removal of mis-localized mitochondrial components. Notably, the mis-localized mitochondrial enzyme involved in folate metabolism exhibited dysregulated metabolic activity^52^. In relation to the evolutionary origin of ATE1 from the mitochondrial microbial ancestor^53^, Nt-arginylation may play a crucial role as the control between the cell and mitochondria.

The temporal quantifications by PRM-MS clarified the dynamics of relative Nt-arginylation under UPR stress. The upregulation of ATF4, UPR stress effector, is expected to be the earliest phenotype while the activation of caspase-3 is the latest under sustained presence of stress that could lead to pro-apoptotic events^54^. Nt-arginylation has a pivotal function over autophagy-apoptosis balance in UPR stress elaborately explained in studies of Nt-arginylated HSPA5 (R-HSPA5)^4, 55^. Interestingly, we found that caspase-3 activation and hence appearance of its cleaved substrates underwent much earlier inception than the appearance of R-HSPA5. Integrating previous research on the anti-apoptotic role of Nt-arginylation, these early Nt-arginylation of caspase-3 targets might be also anti-apoptotic flux as a result of adapting to UPR stress. Our data further demonstrate that Nt-arginylation of caspase-3 targets is relatively rapid compared to proteins with a signal peptide, indicating that potential pro-apoptotic components are more susceptible to this modification than the proteins with cleaved signal peptides, which are likely to be newly synthesized. This indicates that proteins located within organelles might experience a postponed interaction with ATE1, whereas the caspase substrates predominantly found in the cytosol undergo simultaneous Nt-arginylation following their cleavage.

In this study, we introduced ML-based filtering to profile Nt-arginylation *in vivo*. Our findings led to a more comprehensive map of Nt-arginylated proteins during UPR. The best way for Nt-arginylome profiling would be to develop a method to directly enrich the PTM from cells or clinical samples. While such a method has not yet been developed, our method using N-terminomics and highly stringent filtering could be an effective stopgap. We envision that our method contributes to a better understanding of how Nt-arginylation affects protein regulation during various stress conditions including autophagy and apoptosis.

## Methods

### Cell culture

HeLa was grown in DMEM (Gibco, Rockville, MD, USA) medium supplemented with 10% FBS (Gibco) and 1% penicillin/streptomycin (Gibco). The HeLa cells were sourced from ATCC. Cultures were maintained in an atmosphere of 5% CO2 and 95% air in a humidified incubator at 37 °C. Cells were grown to >90% confluence. For ER stress experiments, HeLa cells were treated with 10 µM MG-132 and/or 0.1 µM TG for 24 hours. For TG chase experiments, HeLa cells plated in 6-well plates at about 90% confluency were treated with DMSO, MG132, or MG132/TG for defined lengths of time (0, 3, 6, 12, 18, 24, and 48 h). Cells were harvested by trypsinization, washed thrice with ice-cold PBS (phosphate-buffered saline, pH 7.4; Gibco), and resuspended in an appropriate lysis buffer.

### Cell lysis and protein digestion

For the N-terminome experiment by iNrich, cells were lysed in iNrich lysis buffer (0.2 M EPPS, pH 8.0, 6 M guanidine, 20 mM TCEP, 80 mM 2-chloroacetamide) containing 1× HALT protease inhibitor cocktail (Thermo Scientific). Lysate was boiled for 10 min at 600 rpm and 95 °C, disrupted with ultrasonication (BranSonic 400B), and cleared for 10 min at 10,000 g and 4 °C. Protein concentrations were determined with Pierce BCA Protein Assay Kit (Thermo Scientific). Proteins were precipitated by adding 8× volumes of acetone and 1× volume of methanol to the lysate and incubating overnight at −80 °C. Precipitates were washed twice with methanol and dried briefly.

For PRM-MS and global proteomics, samples were prepared differently. Cells were lysed in 8 M urea in 50 mM Tris-HCl, pH 8.0 containing 1× HALT protease inhibitor cocktail (Thermo Scientific), and disrupted with BranSonic 400B sonifier. The lysate was cleared for 10 min at 10,000 g and 4 °C. Proteins in the lysate were reduced (5 mM DTT, 45 min at 25 °C and 600 rpm), alkylated (20 mM 2-chloroacetamide, 45 min at 25 °C and 600 rpm), and then diluted to bring the urea concentration to <0.8 M using 50 mM Tris-HCl, pH 8.0. Digestion was performed by adding trypsin (Promega, 1:50 enzyme-to-substrate ratio) and incubating overnight at 25 °C and 600 rpm. Digests were acidified to pH < 3 by addition of trifluoroacetic acid (TFA) to 0.5% and were desalted using HLB solid-phase extraction (SPE) cartridges (Waters; wash solvent: 0.1% TFA; elution solvent: 0.1% FA in 50 % acetonitrile (ACN)). Eluates were dried by vacuum centrifugation and stored at −20 °C.

### Enrichment of N-terminal peptides

The precipitated protein sample (1 mg) was reconstituted to 4 mg/mL in 0.25 mL reaction buffer (6 M guanidine in 0.2 M EPPS, pH 8.0). Enrichment of N-terminal peptides was carried out as described previously^24^. Briefly, proteins in the sample were labeled with 200 mM D_6_-acetic anhydride and 200 mM pyridine for 2 hours at 25 °C with end-over-end rotation. Labeled proteins were digested with trypsin or chymotrypsin (Promega, 1:50 enzyme-to-substrate ratio) overnight at 25 °C with end-over-end rotation. The peptides were loaded onto the HLB SPE column (Waters). Depletion of internal peptides was performed by adding 330 mg of NHS-activated agarose dry resin and incubating for 2 hours at 25 °C with end-over-end rotation. The unbound N-terminal peptides were transferred to the stationary phase of the SPE column by drawing under a controlled vacuum. SPE-bound peptides were washed for 20 mL of 0.1% TFA and were eluted using 1 mL of 0.1% FA in 50% ACN. N-terminal peptides were dried by vacuum centrifugation and stored at −20 °C.

### Peptide fractionation by basic reversed-phase liquid chromatography (bRPLC)

For basic reversed-phase liquid chromatography (bRPLC) of N-terminal peptide samples, 100 μg peptides were reconstituted in bRPLC solvent A (10 mM ammonium formate, pH 10) and loaded onto an XBridge BEH C18 RPLC column, 130Å, 3.5 μm (4.6 × 250 mm) and coupled to a 1290 UHPLC system (Agilent). Samples were washed using solvent A for 10 min at 0.5 mL/min and subsequently eluted applying a two-step gradient from 0 to 40% bRPLC solvent B (10 mM ammonium formate, pH 10 in 90% ACN) in 38.5 min, to 70% B in 14 min, and holding at 70% B for 10 min. A total of 168 fractions (0.5 min each) were collected, and then every 12th fraction was pooled to create 12 fractions for N-terminomics or every 24th fraction to create 24 fractions for global proteomics. The pooled fractions were dried and stored at −20 °C until LC-MS analysis.

### LC-MS/MS of N-terminal peptide samples

LC-MS measurements of N-terminal peptide samples were performed with an Ultimate 3000 RSLCnano system coupled to a Q-Exactive mass spectrometer (Thermo Fisher Scientific). bRPLC fractionated peptide samples were reconstituted in 5 μL of 0.1% FA in 2% ACN. 2 µL of samples were injected onto a PepMap 100 trap column (75 µm × 20 mm, Thermo Fisher Scientific), washed with 0.1% FA in 2% ACN for 10 min at a flow rate of 5 µL/min and subsequently transferred to an EASY-Spray PepMap RSLC, 2 µm analytical column (75 µm × 500 mm, Thermo Fisher Scientific). Peptides were separated at 300 nL/min using a 110 min linear gradient from 2.5 to 37.5% LC solvent B (0.1% FA in 80% ACN) in LC solvent A (0.1% FA). MS1 spectra were recorded in the Orbitrap from 400 to 1800 m/z at a resolution of 70,000 and using an automatic gain control (AGC) target value of 1e6 charges and a maximum injection time (maxIT) of 30 ms. Up to 12 of the most abundant precursors (topN) were selected for HCD fragmentation at 27% normalized collision energy (NCE). MS^2^ spectra were acquired at 17,500 resolution using an isolation window of 2.0 m/z, an AGC target value of 5e4 charges, and a maxIT of 120 ms. The dynamic exclusion was set to 30 s.

### Protein sequence databases

A UniProt human reference protein database (Release 2023_02) with common contaminants was used throughout the study. We also constructed specialized decoy databases from this database for FDR estimation of Nt-arginylation search. In case that trypsin was used for protein digestion, any consecutive arginine residues in the original database were consolidated in a single arginine. In our experimental workflow, lysine was modified before digestion and therefore the residue was left untouched in the database. In the case of chymotrypsin, any arginine (or arginines) that immediately follows phenylalanine, leucine, methionine, tryptophan, or tyrosine was removed.

### Database search of mass spectral data

Proteome Discoverer v2.4 with its built-in search engine SequestHT was used to identify and quantify Nt-arginylated peptides. Fragment mass spectra were searched against the protein database in two stages (‘tandem database search’). Mass spectra that failed to pass Percolator validation in the first stage search were collected and used as input data for the second stage search. Search parameters for the first and the second searches were identical except for the types of modifications included. Parameters for both stages were ±10 ppm for precursor tolerance, ±0.05 Da for fragment ion tolerance, up to 2 missed cleavages for the trypsin dataset, and up to 5 missed cleavages for chymotrypsin dataset, fixed modification of carbamidomethylation of cysteine and D_3_−acetylation of lysine, variable modification of methionine oxidation, and cleavage specificity at carboxy-terminal end. Parameters included only in the first stage were N-terminal acetylation and N-terminal D_3_−acetylation as variable modifications. Parameters for the second stage search were pyro-glutamation at N-terminal glutamate, D_3_-acetyl-arginylation (+201.1305 Da) at N-terminal aspartate and glutamate, and D_3_-acetyl-arginylation-deamidation (+202.1145 Da) at N-terminal asparagine and glutamine as variable modification. Minora feature detector module was used for LFQ. Parameters for consensus workflow regarding LFQ: Precursor abundance, area; internal normalization for experimental bias correction, and total peptide amount. For mass spectra of global proteomics, parameters of D_3_−acetylation modifications were removed. Unless stated otherwise, Proteome Discoverer’s default parameters were applied.

When collecting MS2 spectra of Arg-starting peptides resulting from missed cleavage and using decoy databases to estimate FDR of Nt-arginylation modification, we performed database search in a single step (‘conventional database search’). The same parameters as for tandem database search were applied except the following: cleavage specificity to both ends; N-terminal acetylation, N-terminal D_3_-acetylation, and pyro-glutamylation at N-terminal glutamate as variable modifications.

### Construction of MS2 prediction models

AlphaPeptDeep v1.1.5 was used to build MS^2^ prediction models from the training dataset containing PSMs processed with Proteome Discoverer. Four fragment types (b+, b++, y+ and y++) were used to train and predict. Unless stated otherwise, AlphaPeptDeep’s default parameters were applied. To build a “from scratch” model, the pDeep model with ModelMS2Transformer was used. The training parameters were: epoch=100 and batch_size = 100. To build a fine-tuned model, a pre-trained model (“generic”) was used with the “train_ms2_model” function. The training parameter was: epoch=50.

### Spectral similarity scoring

Similarities between two spectra (for example, observed vs. predicted spectra) were determined by calculating Pearson’s correlation coefficient (PCC), cosine similarity (COS), Spearman’s correlation coefficient (SPC), spectral FPR and spectral FNR. These similarity scores were computed using Python (v3.9.18, function calc_ms2_similarity) or R (v4.4.1, function cor). Spectral FPR and spectral FNR for each comparison were calculated as:

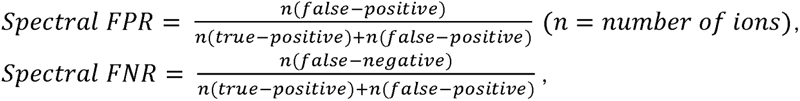

where false positive is a fragment ion appearing in the observed spectrum but not in the predicted spectrum, false negative is that appearing in the predicted spectrum but not in the observed spectrum, and true positive is that appearing in both spectra.

### FDR estimation of MS2 prediction model for detecting Nt-arginylation

FDR for detecting Nt-arginylation was estimated based on the fact that Arg-starting peptide is searched as a missed cleavage peptide in the original database, but as Nt-arginylated peptide without the first residue Arg in the decoy database. Any PSMs found in this way were treated as true positives. Nt-arginylation PSMs that mapped to sequences other than those immediately following the modified sequence were treated as false positives. By sorting the Nt-arginylated PSMs in descending order of a specific score and applying an arbitrary threshold value, we could calculate the FDR. Conversely, the threshold was set to satisfy an arbitrary FDR value. In most cases, FDR was set to be less than 0.01.

### Construction of RT prediction models

AlphaPeptDeep v1.1.5 was used to build RT prediction models from the training dataset containing PSMs processed with Proteome Discoverer. PSMs from each LC-run were used separately for the training, taking into account the degradation of analytical columns with use. The observed RTs were normalized by dividing by the time length of the LC gradient. Fine-tuning of a pre-trained model was performed with the same parameters as for MS2 except using train_rt_model function instead of the train_ms2_model function. We then collected pairs of RT predicted by the fine-tuned RT prediction model *versus* actual measured RT, performed linear regression, and calculated the RT deviation that included 95% of all data points (Δ*t*_95%_) by using R (function lm and metrics).

The fine-tuned RT prediction model thus constructed was used in the next step to predict the RT of PSM for Nt-arginylation in the search result. The RT prediction was performed in Python (function predict_rt). If the predicted RT is within Δ*t*_95%_, the PSM is considered true, otherwise it is considered false.

Each RT prediction model was only used to analyze the PSMs of Nt-arginylation identified in the same dataset used to build that model. In LC-PRM-MS experiments, a mixture of HeLa digest and PRTC (Thermo Fisher Scientific) as internal standards was analyzed by LC-MS/MS in DDA mode before every six PRM experiments. The dataset collected in this DDA mode was used to build an RT prediction model, which was used to analyze the following six PRM data.

### Mass Error Test

For each PSM of Nt-arginylation, m/z errors of fragment ions were calculated by subtracting theoretical m/z values from the measured m/z values. A two-sided Student’s t-test was performed between the m/z errors of b ions and those of y ions in R (function t.test). A PSM with a *P*-value ≥ 0.05 (that is, an insignificant *P*-value) was considered true, otherwise, it is considered false.

### Structural annotation of Nt-arginylation sites

For structural annotation of the Nt-arginylated sites, we utilized information in the UniProt knowledgebase or used bioinformatic tools such as dagLogo^27^, DeepLoc^56^, SignalP^35^, TargetP^36^, Procleave^37^ and Alphafold^57^. The dagLogo R/Bioconductor package v1.28.1 was used to analyze differential amino acid usage (DAU) of Nt-arginylated peptides. A background model was prepared from UniProt protein database (Release 2023_02), with an argument of “fisher” test type. A total of 10 amino acids from P5 to P5’ in the sequence were tested against the background model for statistical significance (function testDAU). Visualization was performed by functions of dagHeatmap for heatmap plot and dagLogo for sequence logo plot. The predicted protein localization was determined using DeepLoc v2.0. The sequences of proteins corresponding to the identified Nt-arginylation sites were extracted from the database and used for DeepLoc analysis. Each protein was assigned a localization that exhibited a probability exceeding 0.5. Proteases predicted to cleave each Nt-arginylation sites were acquired using Procleave algorithm, inputing the sequence from P4 to P4’ into the algorithm. Predictions were made for all 27 accessible proteases. pLDDT scores of AlphaFold were acquired using protti R package (v0.9.1, function fetch_alphafold_prediction) together with AlphaFold database version v4. The pLDDT score was calculated for a total of 21 amino acids, including the N-terminal 10 and the C-terminal 10 amino acids from the Nt-arginylation site.

### Functional annotation of Nt-arginylation sites

Reactome^58^, Gene Ontology^59^, and STRING^60^ were used for functional annotation of Nt-arginylation sites. Reactome, Gene Ontology, and GESA results were obtained using clusterProfiler R package v4.12.6 (function enrichPathway, enrichGO, and gsePathway, repectively)^61^. The *P*-values were corrected by Benjamini-Hochberg method^62^ and the cutoff was set to 0.05. Protein-protein interaction network was obtained from STRING with medium confidence (combined score > 0.4) and visualized using Cytoscape v3.9.1^63^.

### Plasmid construction for the arginylated protein candidates

HeLa cells, as well as wild-type (+/+) and *ATE1*^-/-^ MEFs, were cultured in Dulbecco’s Modified Eagle Medium (DMEM/high glucose; HyClone, Cat# SH30243.01) supplemented with 10% fetal bovine serum (FBS; Gibco, Cat# 16000044) in a 5% CO2 incubator at 37°C.

Total RNA was isolated using TRI Reagent (Molecular Research Center, Cat# TR 118). 2 μg of the isolated total RNA were utilized for cDNA synthesis using TOPscript™ RT DryMIX (Enzynomics, Cat# RT220). The constructs for arginylated protein candidates were generated through PCR amplification from a human cDNA library and subsequently subcloned into the pcDNA 3.1 myc/his B plasmid (Invitrogen) using specific restriction sites. The primer information used in these experiments is provided in Supplement table 2).

### R-catcher pulldown assay

Plasmids encoding the arginylated protein candidates were transiently transfected into HeLa cells using XtremeGene HP DNA transfection reagent (Roche, Cat# C756V59) according to the manufacturer’s protocol. After transfection for 24 hours, cells were treated with 10 μM MG132 and 50 nM thapsigargin, followed by incubation for 24 hours. Cells were then collected in cold phosphate-buffered saline (PBS) and centrifuged at 500 x g for 5 minutes. The cell pellets were resuspended in hypotonic buffer containing protease and phosphatase inhibitors. The resuspended cells were lysed by undergoing at least five cycles of freezing and thawing using liquid nitrogen and 37°C water bath, followed by centrifugation at 15,928 x g for 20 minutes at 4°C. We have previously described detailed protocols for the R-catcher pulldown assays^11, 64^. Briefly, 300 μL of purified R-catcher WT and mutant (D129A) proteins were conjugated with 120 μL (50% slurry) of Strep-Tactin Sepharose resin overnight at 4°C. Subsequently, 400 μg of cell lysates from MG132- and thapsigargin-treated cells were diluted in 940 μL of binding buffer (0.05% Tween 20, 10% glycerol, 0.2 M KCl, and 20 mM HEPES at pH 7.9) and mixed with 60 μL (in packed volume) of R-catcher-conjugated beads. The mixtures were gently rotated at 4°C for 3 hours. The beads were collected by centrifugation at 4,600 x g for 1 minute, washed five times with 1 mL of binding buffer at 4°C for 10 minutes, resuspended in 50 μL SDS sample buffer, and heated at 100°C for 10 minutes. The protein samples were then separated by SDS-PAGE and transferred onto a polyvinylidene difluoride (PVDF) membrane (Cytiva, Cat# 10600023) at 35 V overnight at 4°C. The protein-bound PVDF membrane was subsequently blocked with 5% skim milk in TBS-T buffer for 1 hour at room temperature. The membrane was then incubated with the primary Myc antibody overnight at 4°C, followed by a 1-hour incubation with a host-specific HRP-conjugated mouse secondary antibody at room temperature. Protein bands were visualized using an enhanced chemiluminescence (ECL) solution (Thermo Fisher Scientific, Cat# 32106) and X-ray films.

### PRM-MS

PRM-MS was performed on the same LC-MS instrument as in the profiling experiments. A 2 µg peptide sample spiked with 125 fmol of PRTC peptide standard (Thermo Scientific) was injected onto a PepMap 100 trap column (75 µm × 20 mm, Thermo Fisher Scientific), washed with 0.1% FA in 2% ACN for 10 min at a flow rate of 5 µL/min and subsequently transferred to an EASY-Spray PepMap RSLC analytical column (75 µm × 150 mm, Thermo Fisher Scientific). Peptides were separated at 300 nL/min using a 55 min linear gradient from 2.0 to 24.0% LC solvent B (0.1% FA in 80% ACN) in LC solvent A (0.1% FA). Similar MS settings as described above were used, but the MS was operated in PRM mode with the following adjustments: MS1 scans were recorded from 300 to 1500 m/z using 250 ms for maxIT. Targeted MS^2^ spectra were recorded at a resolution of 35,000 and using an AGC target value of 2e5 charges, a maxIT of 200 ms, an isolation window of 1.2 m/z, and an isolation offset of 0.4 m/z. The number of targeted precursors per cycle was set to 20. The first mass was fixed to 200 m/z. Per six PRM LC-MS analyses, 0.5 µg of HeLa digest/PRTC standard (Thermo Fisher Scientific) was injected and analyzed with DDA method and identical LC gradients to PRM-MS. The inclusion list was comprised of 65 targets including 15 PRTC, 6 UPR related proteins, 21 Nt-arginylated peptides and 21 corresponding non-Nt-arginylated peptides. The charge state of each target peptide was determined using the ‘iep’ application of EMBOSS. The PRM acquisition times for the targets were specified as -2 minutes for the start time and +4 minutes for the end time, based on the predicted RT. Out of 15 PRTCs, 14 except the most hydrophobic ones were used as internal standards (ISTD).

MS2 spectra were assigned to the targets by comparing the charge state and MS1 m/z value (±0.001 m/z). For each target, theoretical MS2 spectrum was generated using a fine-tuned MS2 prediction model and used for spectral matching within 5 ppm error. PRM spectra exhibiting PCC values greater than 0.7, which represented the minimum PCC of PRTC standards, were selected, of which the spectra having at least 4 fragments were used for subsequent analysis. The intensities for fragment ions were integrated to get target peptide intensities and then normalized by the PRTC intensities to correct for run-to-run variation. The normalization was done using the crmn^65^ R package (v0.0.21). The final LFQ intensities were acquired using MSFragger (v4.0)^66^ and IonQuant (v1.10.12) in the platform of FragPipe (v21.1). Match-between-run workflow “LFQ-MBR” was used under the default parameters except for the following: fragment ion tolerance to ±0.05 Da, arginylation (+156.1011 Da) at N-terminal aspartate and glutamate, and arginylation-deamidation (+157.0851 Da) at N-terminal asparagine and glutamine as variable modification.

### Immunoblotting

Cells were lysed using ice-cold RIPA buffer (Sigma Aldrich, R0278) supplemented with a protease inhibitor cocktail and agitated for 30 min at 4 °C. Lysates were subsequently probe-sonicated and centrifuged at 16,000 × g for 20 min at 4°C. Protein concentrations were determined using Pierce BCA Protein Assay Kit (Thermo Scientific). For western blot analysis, 1 µg of protein was separated by SDS-PAGE to detect all targets except cleaved caspase-3. For the detection of cleaved caspase-3, the same lysate was concentrated using 3 kDa molecular weight cutoff centrifugal filter (Amicon), followed by protein quantification with BCA assay, and 5 µg of protein was separated by SDS-PAGE. After completion of the electrophoresis, proteins were transferred to a PVDF membrane at 100 V for 1 h. The membrane was blocked with 5% skim milk in TBS-T (20 mM Tris, 150 mM NaCl, and 0.1% Tween 20, pH 7.5) for 1 h at room temperature, followed by overnight incubation at 4°C with the appropriate primary antibodies, diluted in a phosphate buffered saline (PBS) solution containing 1% bovine serum albumin (BSA) and 0.02% sodium azide. After incubation, the membranes were washed with TBS-T three times and treated with the rabbit IgG-HRP secondary antibodies (1:1,000,000 dilution in 5% skim milk) for 1 h. The membranes were washed with TBS-T and visualized with the ECL chemiluminescent substrate (Thermo Fisher Scientific, A38555). Subsequently, the PVDF membranes were stripped with West Ez Stripping Buffer (GenDEPOT, S2100-050) following the manufacturer’s protocol and re-probed to detect multiple proteins of interest (POIs) and a loading control on the same blot.

The antibodies used are as follow: rabbit polyclonal anti-HSPA5 (3183, 1:5000, Cell Signaling Technology), rabbit polyclonal anti-R-HSPA5 (ABS2103, 1:5,000, Sigma-Aldrich), rabbit polyclonal anti-CRT (1:2,000, courtesy of Dr. Yong Tae Kwon), rabbit polyclonal anti-R-CRT (1:2,000, courtesy of Dr. Yong Tae Kwon), rabbit polyclonal anti-PDI (1:1,000, courtesy of Dr. Yong Tae Kwon), rabbit polyclonal anti-R-PDI (1:2,000, courtesy of Dr. Kwon Yong Tae), rabbit monoclonal anti-ATF4 (11851, 1:2,000, Cell Signaling Technology), rabbit polyclonal anti-caspase-3, cleaved form (9661, 1:500, Cell Signaling Technology), rabbit polyclonal anti-GAPDH (ab9485, 1:10,000 abcam), and goat anti-rabbit IgG-HRP (17492, 1:100,000, Cell Signaling Technology).

## Data availability

The MS proteomics data reported here were deposited to the ProteomeXchange Consortium via the PRIDE partner repository with the dataset identifiers PXD058868 and PXD058872. The data was also deposited in KPOP (Korea ProteOme rePository, https://kbds.re.kr/KPOP) with the accession ID KAP241007. All data are available in the main text and the supporting information. Further details are also available upon reasonable request.

## Code availability

The code for the ML-based filtering and data analyses is available at https://github.com/syju1984/Nt-arginylationFiltering.

## Acknowledgements

This work was supported by grants from the National Research Foundation of Korea (RS-2023-00279134, RS-2022-NR068428, RS-2024-00444177), a grant from the National Research Council of Science & Technology (GTL24021-000), a KIST intramural program (Grand Challenge), and the KRIBB Research Initiative Program (KGM1062413).

## Author contributions

S.J. and C.L. conceptualized the study. S.J. and C.L. developed the methodology. S.J. wrote the algorithm. L.N., S.L. and J.G.K. performed immunoblot validation. S.L., and H.L. performed targeted proteomics. D.H.K. and H.C.M. provided resources. S.J., L.N., S.L., J.G.K., H.L., N.P., H.C.M. and C.L. curated the data. All authors wrote the paper. C.L. supervised the project and was the project administrator. C.L. acquired funding.

