## Supplementary Information for "Implementing N-terminomics and machine learning to probe *in vivo* Nt-arginylation"


### Table of Contents


| | Pages |
|---|---|
| Supplementary Figure 1 | 3 |
| Supplementary Figure 2 | 4 |
| Supplementary Figure 3 | 5 |
| Supplementary Figure 4 | 6 |
| Supplementary Figure 5 | 7 |
| Supplementary Figure 6 | 8 |
| Supplementary Figure 7 | 9 |

| | |
|---|---|
| Supplementary Figure 8 | 10 |
| Supplementary Figure 9 | 11 |
| Supplementary Figure 10 | 12 |
| Supplementary Figure 11 | 13 |
| Supplementary Figure 12 | 14-15 |
| Supplementary Figure 13 | 16 |
| Supplementary Figure 14 | 17 |
| Supplementary Figure 15 | 18 |
| Supplementary Figure 16 | 19 |
| Supplementary Figure 17 | 20 |
| Supplementary Figure 18 | 21 |
| Supplementary Figure 19 | 22 |
| Supplementary Figure 20 | 23 |
| Supplementary Figure 21 | 24 |
| Supplementary Figure 22 | 25 |
| Supplementary Figure 23 | 26-29 |
| Supplementary Figure 24 | 30 |
| Supplementary Figure 25 | 31-33 |
| Supplementary Figure 26 | 34 |
| Supplementary Table 1 | 35-37 |
| Supplementary Table 2 | 38 |

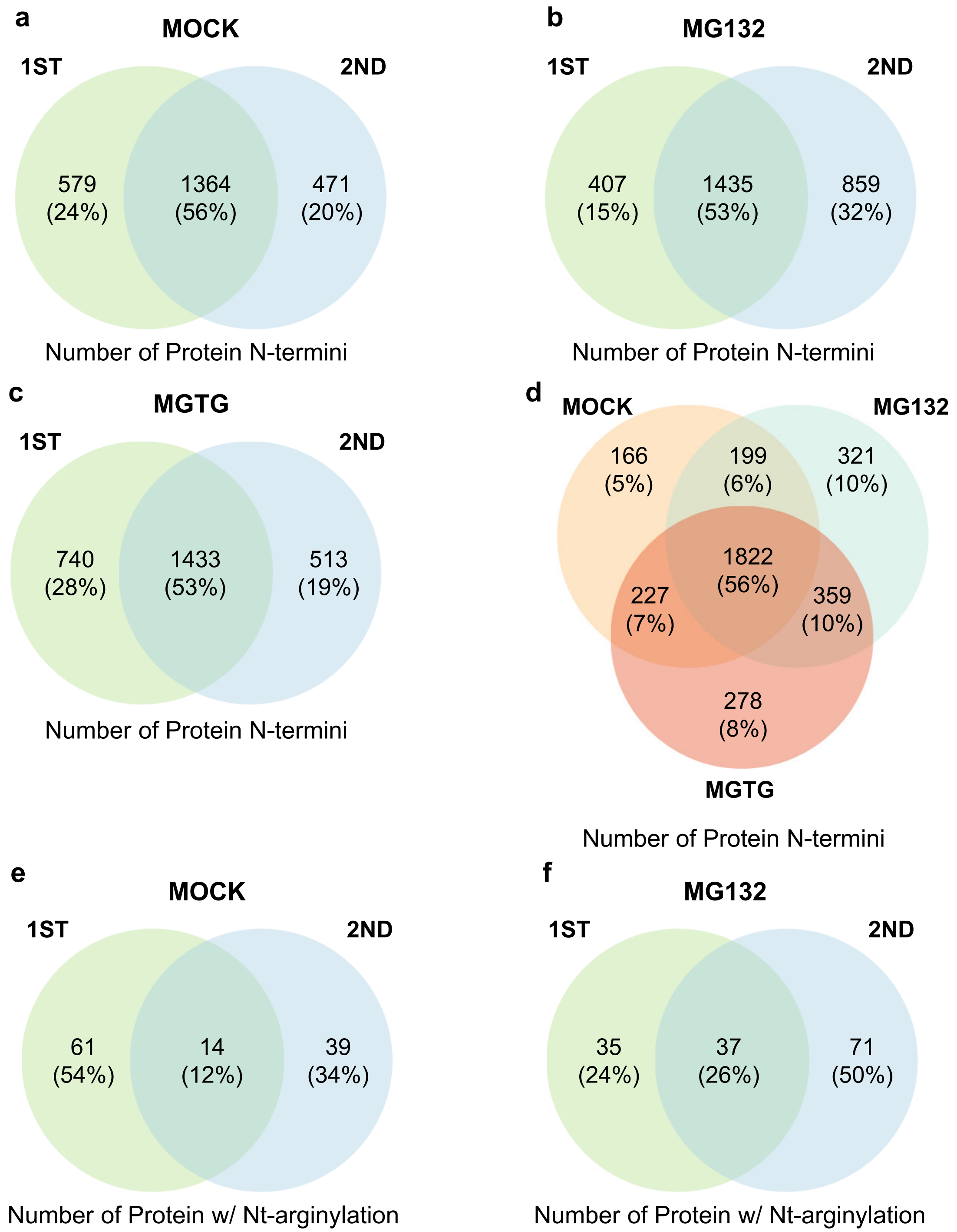


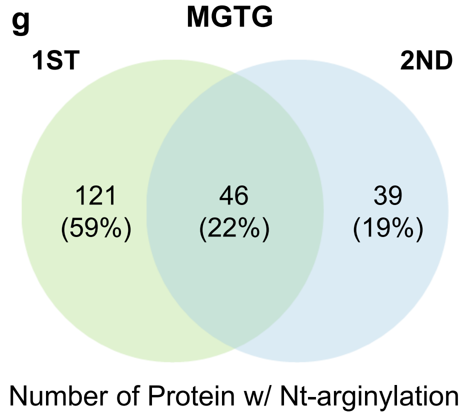


**Supplementary Figure 1. Identified N-terminome in this study**

**a-c,** Venn diagrams showing the number of shared protein N-termini identified in each replicate. The results from the treatment of DMSO (a), MG132 (b), and MG132 and thapsigargin (c). **d,** Shared protein N-termini between three treatments are presented.

**e-f,** Shared proteins with the PSMs searched as Nt-arginylated peptides in each replicate. The results from the treatment of DMSO (e), MG132 (f), and MG132 and thapsigargin (g).

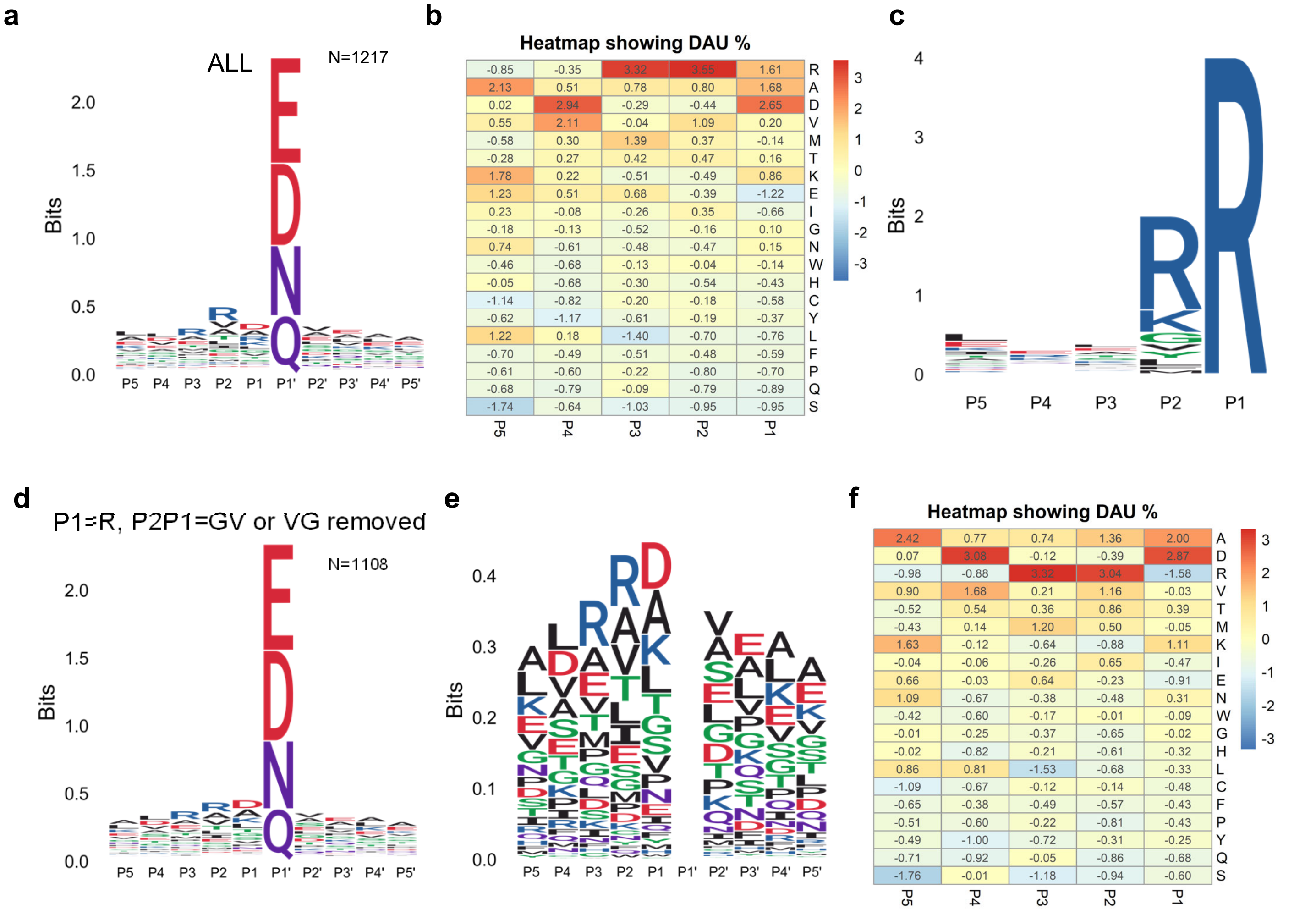


**Supplementary Figure 2. Positional analysis for the putative Nt-arginylated peptides.**
**a**, Logo analysis for the putative Nt-arginylated peptides. **b,** A differential amino acid usage (DAU) analysis for the putative Nt-arginylated peptides. Z-scores of DAU tests for the amino acid residue and the P5 to P1 positions are shown in the heatmap. Z-scores were derived from a Z-test with a background model that was built from randomly sampled sequences from the UniProt human reference proteome database. **c**, Logo analysis for the putative Nt-arginylated peptides that exclusively contain an arginine at the P1 position. **d, e** Logo analysis for the peptides depicted in figure A, specifically focusing on those that do not contain an R at position P1 or GV|VG at positions P2P1 (d) and enlarged figure by omitting residues at P1' (e). **f**, DAU analysis for the PSMs in **(d)**.

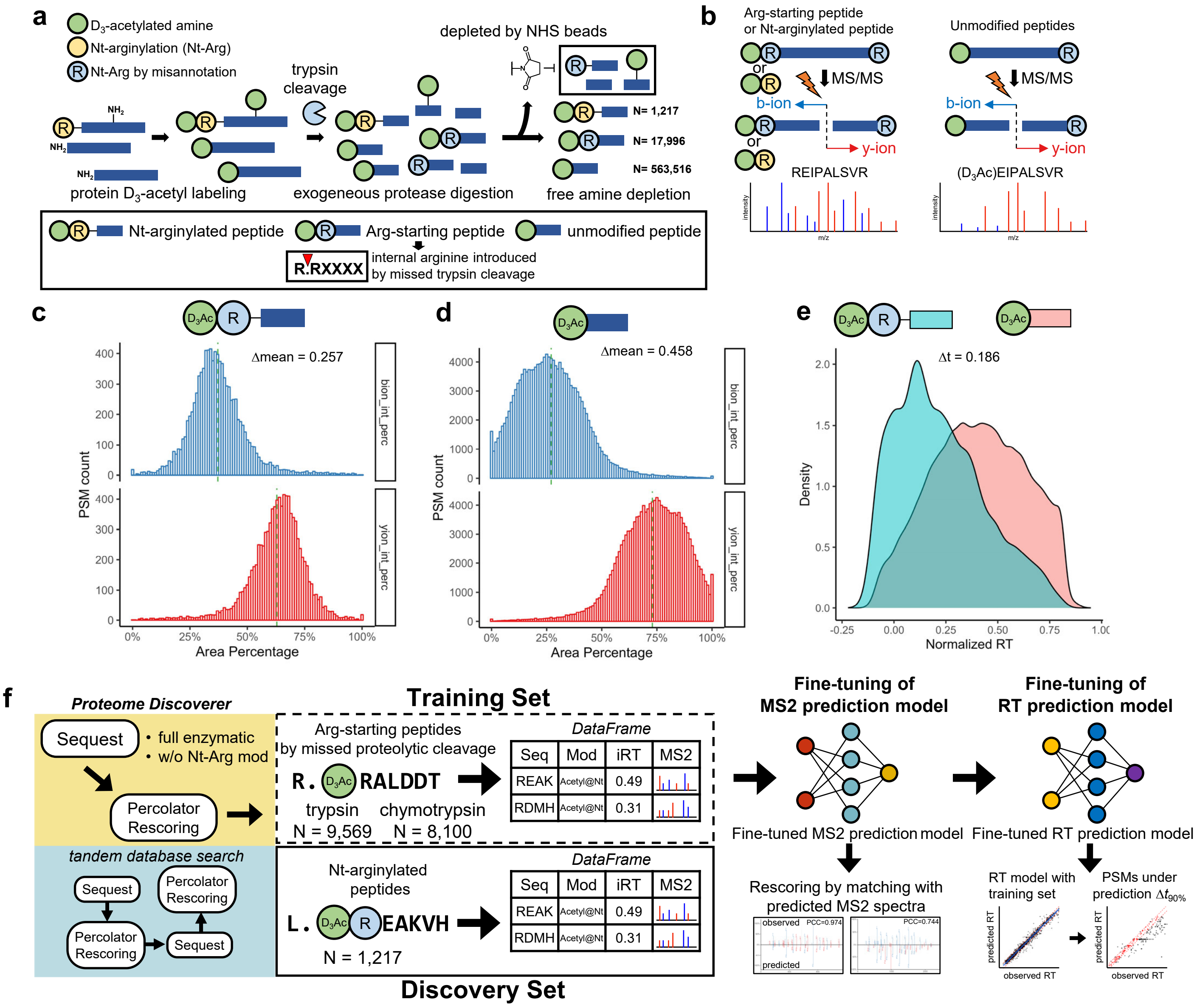


**Supplementary Figure 3. Characteristics of Arg-starting peptides.**
**a**, A schematic representation of the generation of Arg-starting peptides resulting from the missed trypsin cleavage products during an N-terminomic experiment. Numbers (N) indicate the PSMs for each N-terminal peptide species discovered in this study. **b**, The visual representation highlights the physicochemical characteristics of Nt-arginine in modulating b-ion generation subsequent to MS/MS fragmentation. Arginine's significant pKa contributes to its positive charge, which in turn causes fragment ions containing arginine to show greater intensity. **c, d** The total shared intensity distributions of fragment ion species are elucidated for each spectrum. Distributions of spectra from Arg-starting peptides in **(c)** and non-Arg-starting peptides in **(d)**. **e,** RT distribution for Arg-starting peptides and non-Arg-starting peptides. $\Delta t$ indicates a difference in the median RT for two separate groups of peptides. **f,** Schematic representation of the model-training workflow for $MS^2$ spectra and RT prediction. Arg-starting peptides are used as a training set for fine-tuning a pre-trained MS2 or RT prediction model. Nt-arginylated peptides are tested their MS2 spectra and RT with the fine-tuned prediction models.

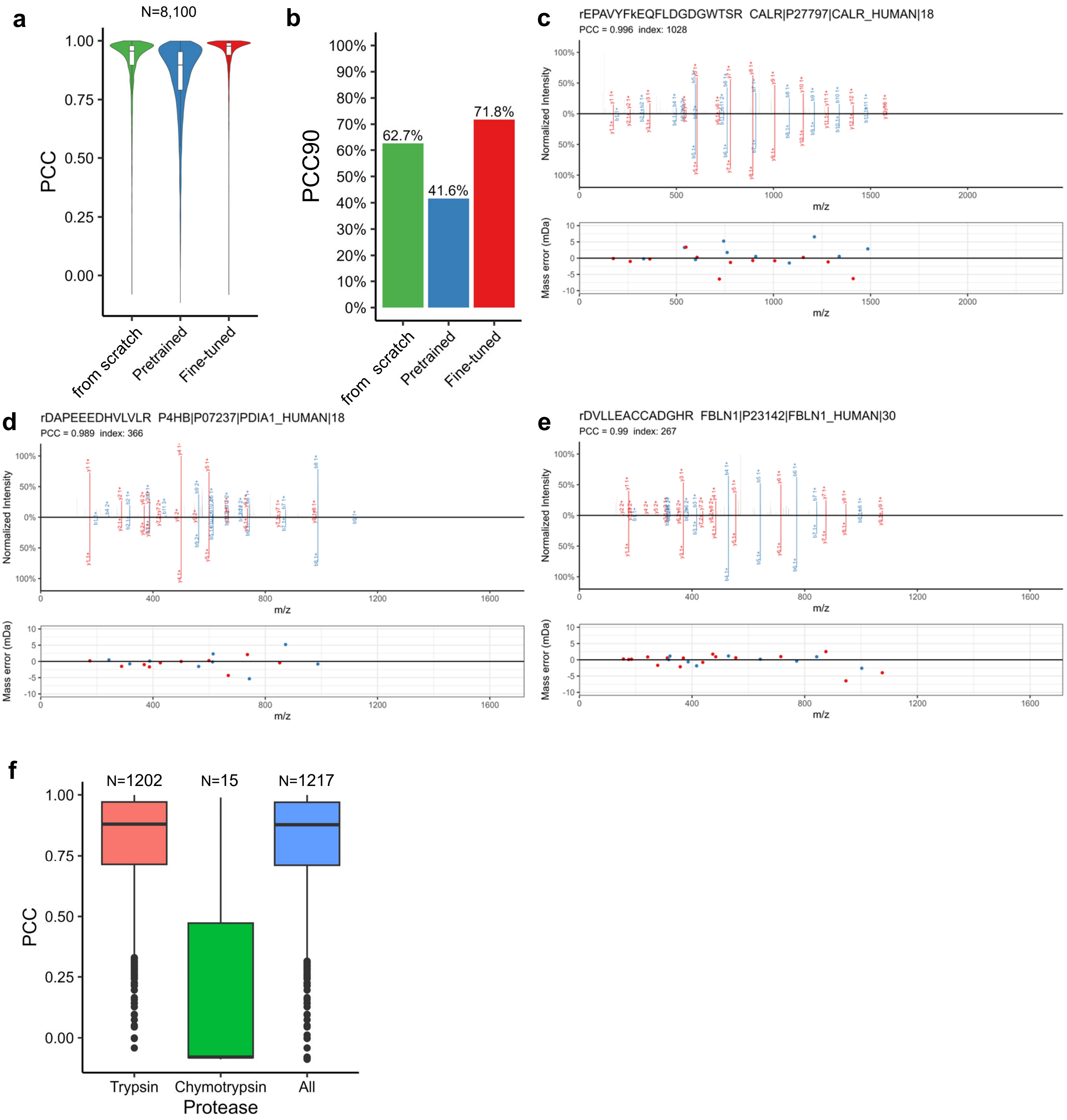


**Supplementary Figure 4. Quality of data from MS2 prediction model.**

**a,b** Assessment of prediction accuracy for MS2 with varying training methods by distribution of Pearson's correlation coefficient (PCC) (a), percentage of PSMs with PCC90 for each prediction model (b). **c-e,** Spectra mirror plots for the selected Nt-arginylated peptides. Mass spectrum shown in upper plane is an observed spectrum, and bottom plane is a predicted spectrum which was derived from a fine-tuned MS2 prediction model. Lower cases in the peptide sequence denote as follows: r; Nt-arginylation, k; trideuteroacetylation. Indentifier description: (gene name)|(UniProt accession)|(UniProt protein name)|(Nt-arginylation site position). **f,** Prediction performance of the protease-specific fine-tuned MS2 prediction models for PSMs searched as Nt-arginylated peptides.

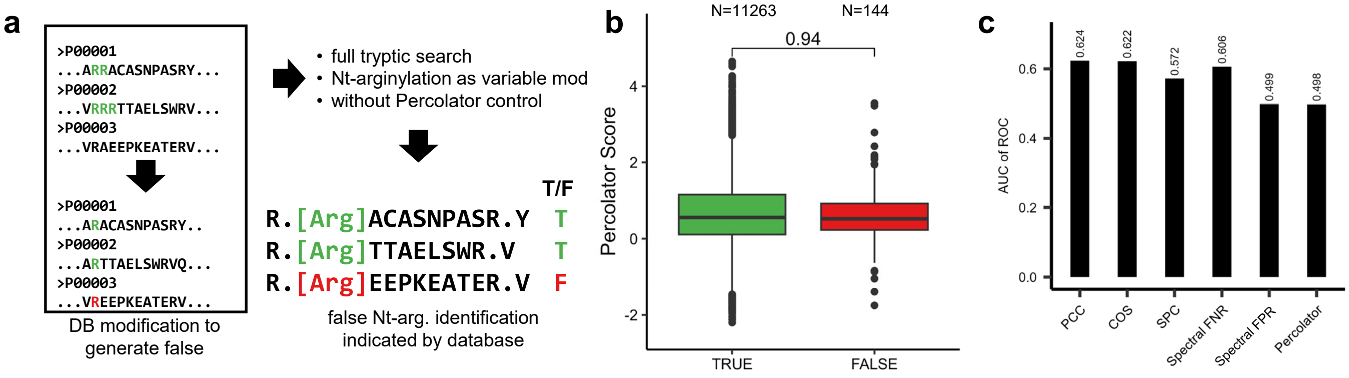


**Supplementary Figure 5. Nt-arginylation specific target-decoy search strategy.**
**a,** Schematic representation of the generation of Nt-arginylation specific target-decoy search strategy. The Nt-arginylated peptides originating from altered location were denoted as true (T), from unaltered location as false (F). **b,** Evaluation of Percolator scores of the Nt-arginylated peptides based on the true or false hit by the decoy database. A two-tailed Student's t-test was used to analyze results. **c,** Evaluation of AUC of ROC analysis based on similarity scoring methods.

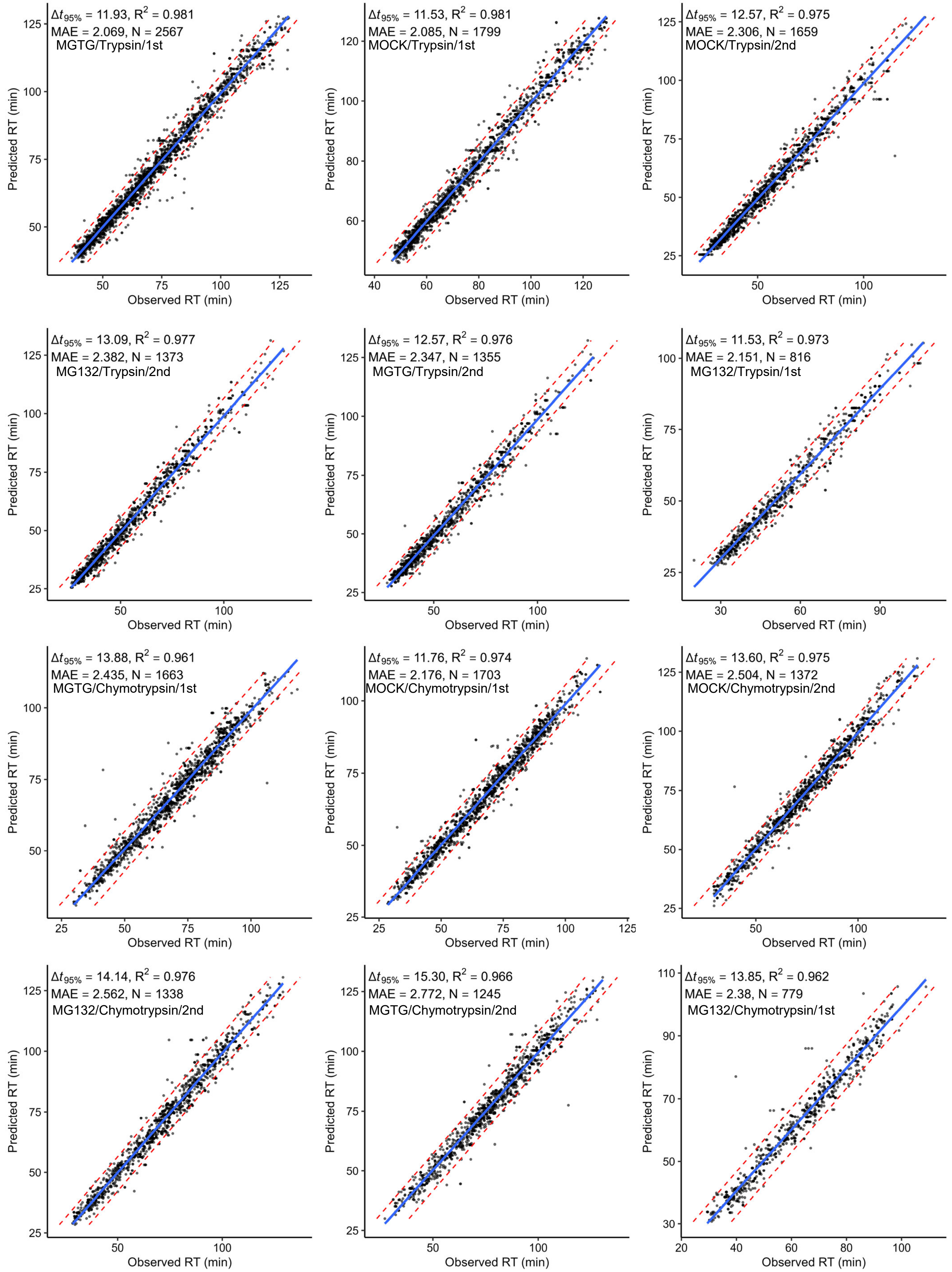


**Supplementary Figure 6. Prediction performance of the RT prediction model.**

For the 12 data sets, the observed RT of Arg-starting peptides is plotted against the predicted RT. The data identification string is denoted as Experiment/Protease/Replicates. Dotted line in red, $\Delta t_{95\%}$ indicating line.

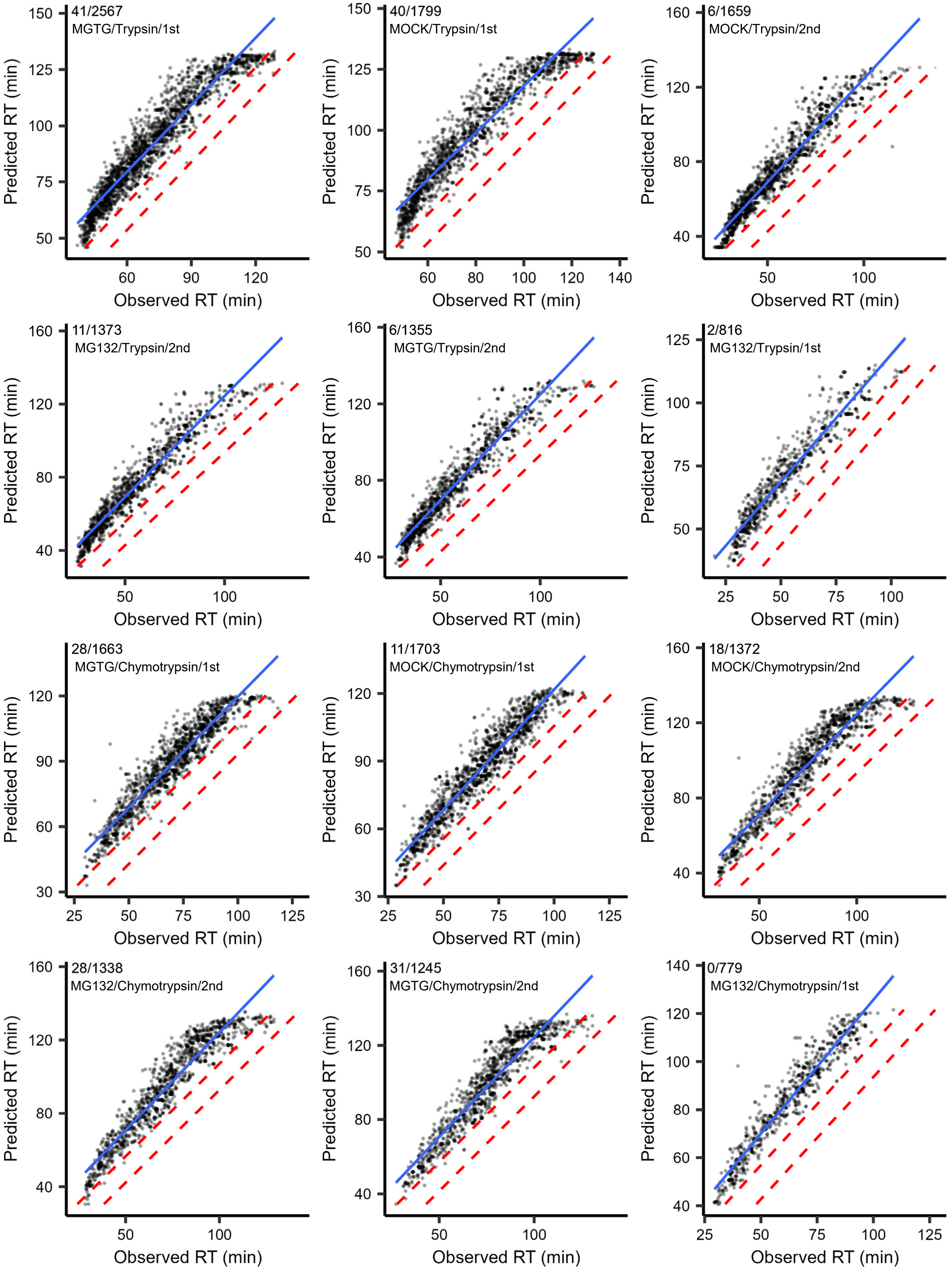


**Supplementary Figure 7. Validation of the RT prediction model.**
For the 12 data sets, scatter plots show the predicted RT as Arg-starting peptide with GV substitution against as the observed RT. The numerator of each fraction indicates the spectral counts of Arg-starting peptides within $\Delta t_{95\%}$ of the RT model. The data identification string is denoted as Experiment/Protease/Replicates. Dotted line in red, $\Delta t_{95\%}$ indicating line.

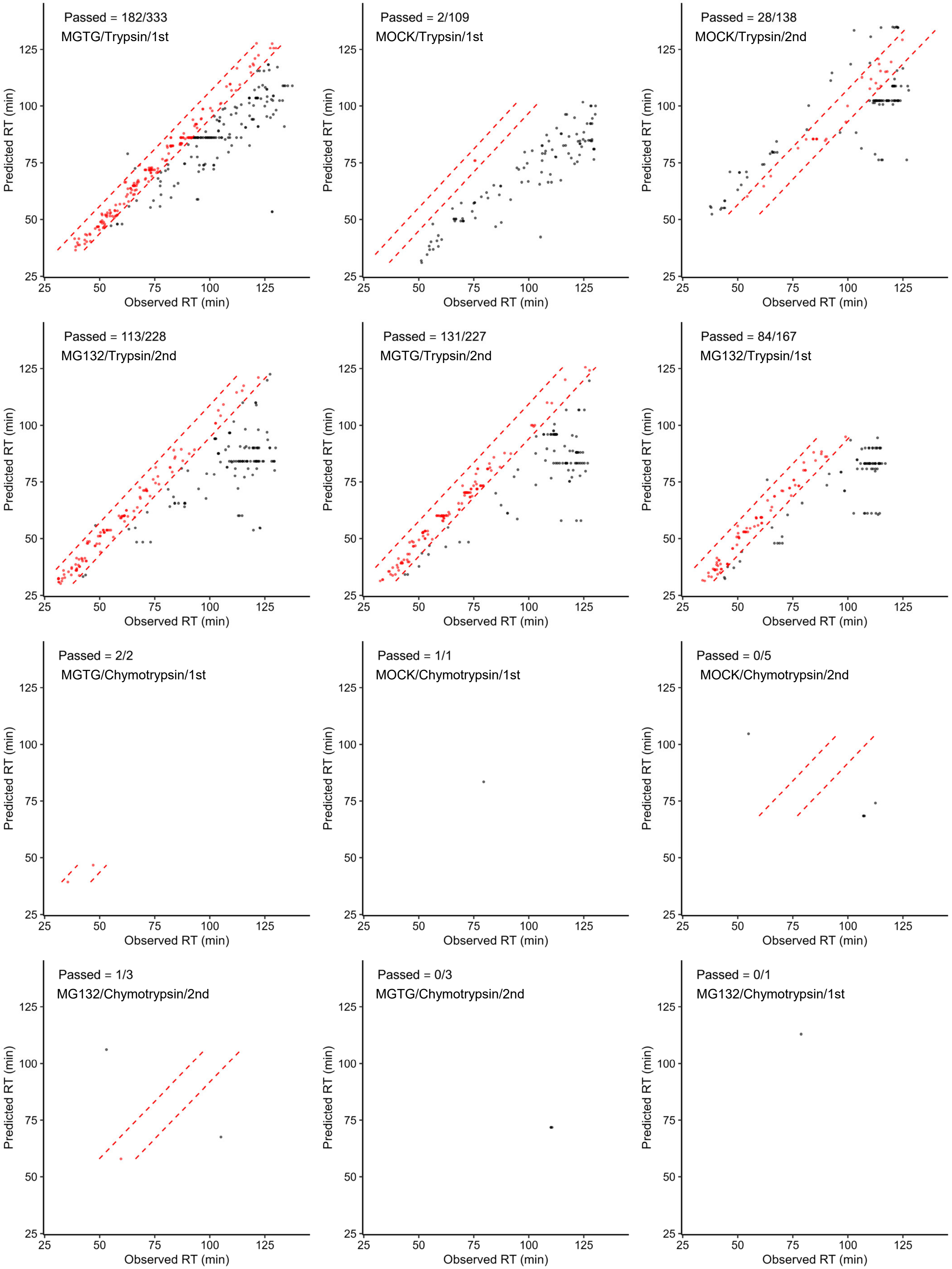


**Supplementary Figure 8. Prediction performance of the RT prediction model.**
For the 12 data sets, the observed RT of the PSM searched as Nt-arginylated peptides is plotted against the predicted RT. The numerator of each fraction indicates the spectral counts of the PSM searched as Nt-arginylated peptides within $\Delta t_{95\%}$ of the RT model. Data identification string is denoted as Experiment/Protease/Replicates. Dotted line in red, $\Delta t_{95\%}$ indicating line.

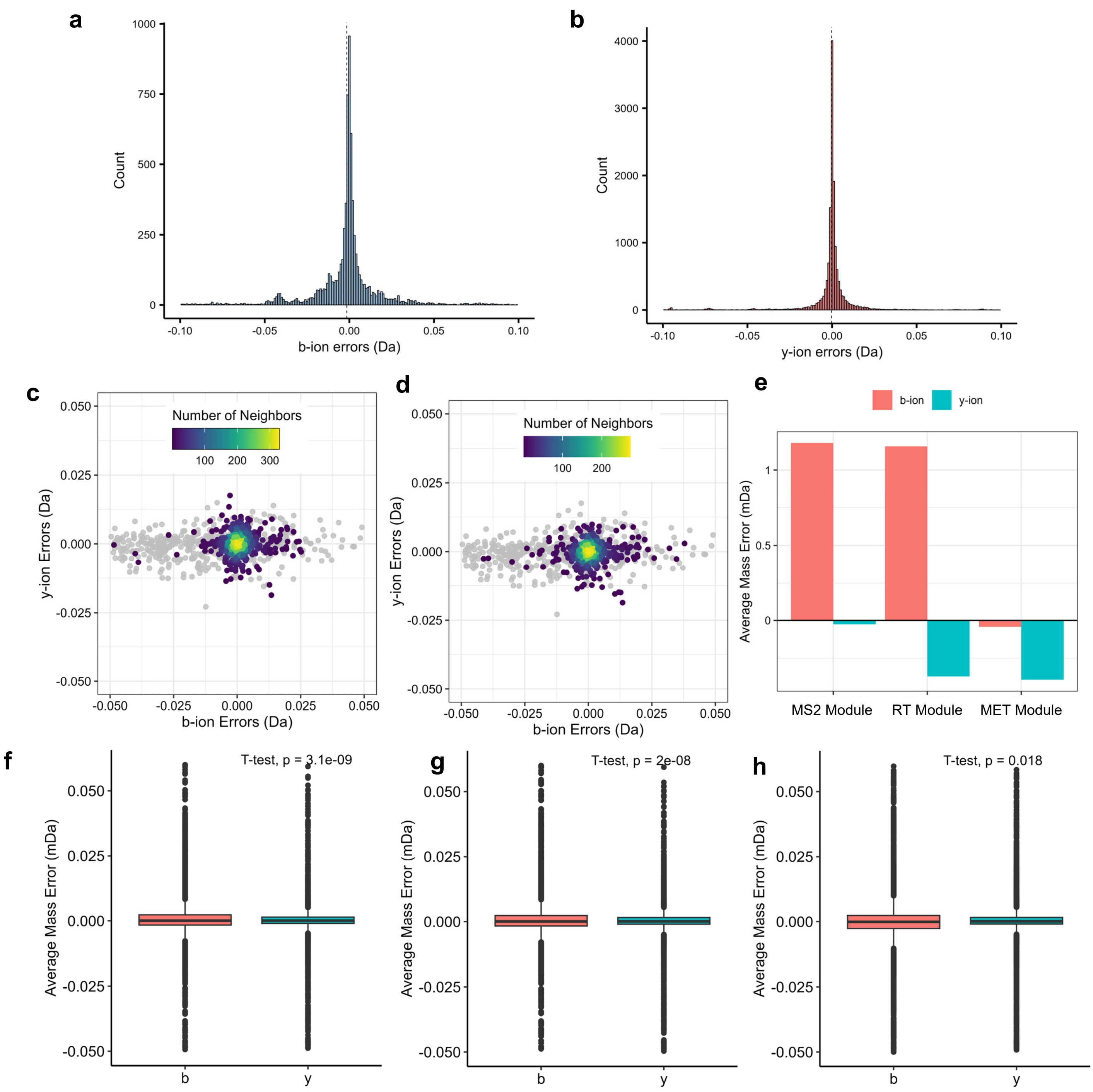


**Supplementary Figure 9. Development of MET module.**
**a,b** Distribution of b-ion errors (**a**) and y-ion errors (**b**) of the putative Nt-arginylated PSMs. Dotted line, median. **c,d** Comparisons of mass errors of fragment ion species. The data from PSMs that passed MS2 module plotted in (**c**) and RT module in (**d**). Gray, removed PSMs through each modules. **e,** Average error for b-ions and y-ions of the passed PSMs through each modules. **f-h**, Box plots of mass error for b-ions and y-ions. A two-tailed Student's t-test was used to analyze results. PSMs that are passed MS2 module (**f**), RT module (**g**), and MET module (**h**).

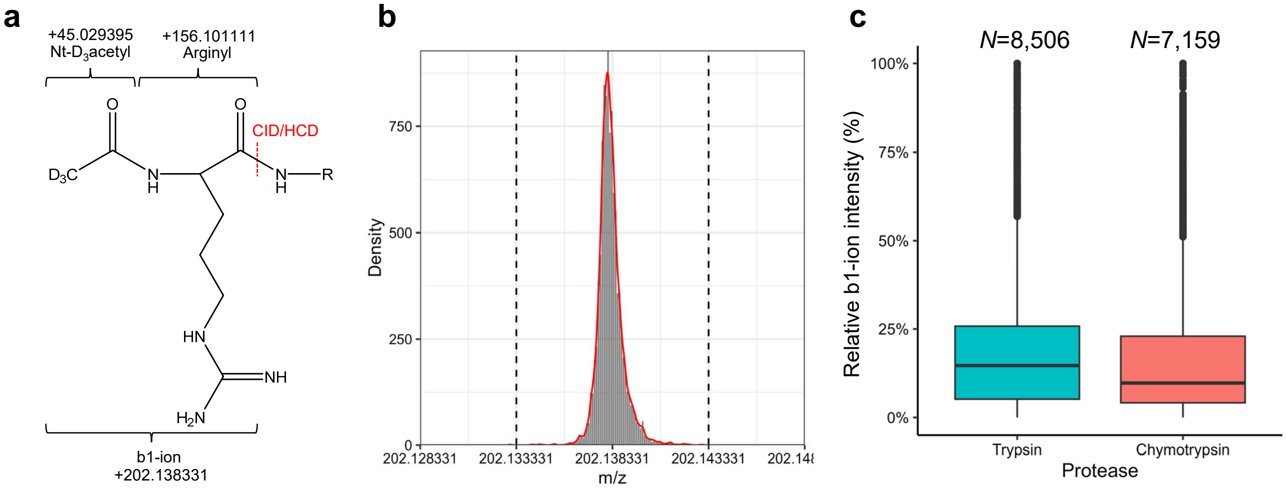


**d**

| Arg-starting peptide | b1-ion found | b1-ion not found | Total |
|---|---|---|---|
| YES | 8,506 (88.9%) | 1,063 (11.1%) | 9,569 |
| NO | 3,296 (1.1%) | 298,682 (98.9%) | 301,978 |
| Total | 11,802 | 299,745 | 311,547 |

| Arg-starting peptide | b1-ion found | b1-ion not found | Total |
|---|---|---|---|
| YES | 7,159 (88.4%) | 941 (11.6%) | 8,100 |
| NO | 3,301 (13.5%) | 240,568 (98.6%) | 243,869 |
| Total | 10,460 | 241,509 | 251,969 |

**Supplementary Figure 10. b1-ion of $D_3$-acetylated arginine.**
**a,** Chemical structure and theoretical mass of a b1-ion of $D_3$-acetylated Arg after fragmentation. Red dashed line shows site of fragmentation. **b,** Mass distribution of the b1-ion in PSMs of Arg-starting peptide. The dotted lines indicate the mass tolerance (±0.005 m/z) for the extracting b1-ions. **c,** Ratio of intensity of each b1-ion relative to total MS2 intensity of each PSMs. **d,** Contingency tables summarizing the result of b1-ion presence in trypsin (left) and chymotrypsin (right) experiments.

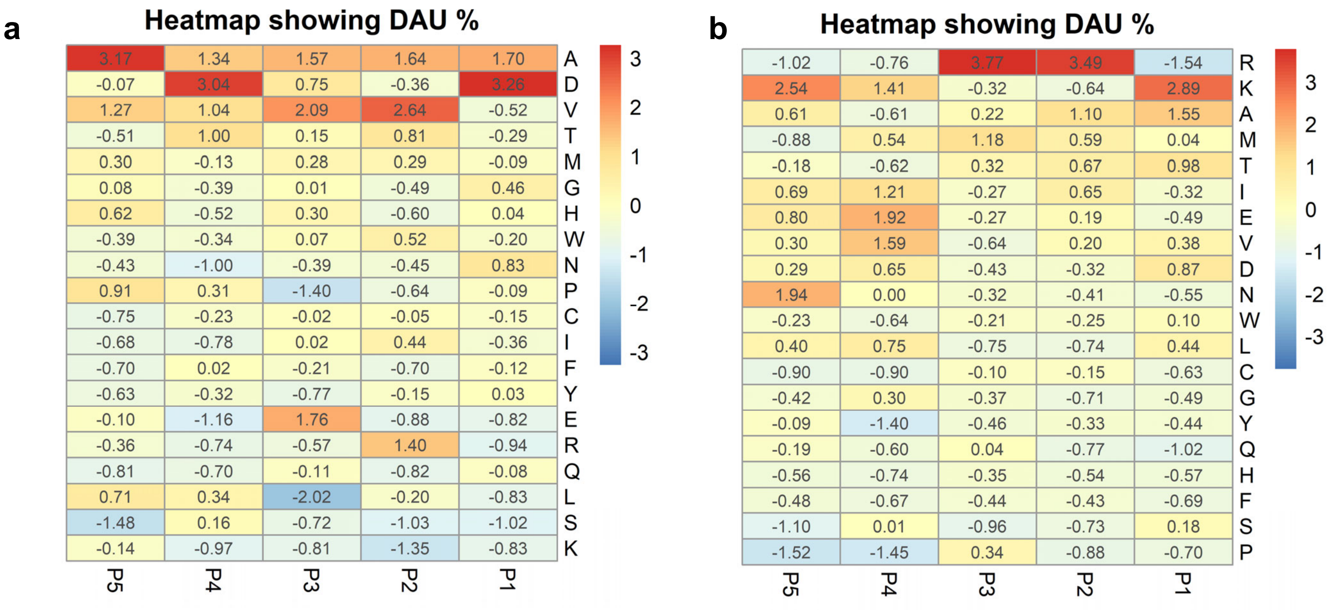


**Supplementary Figure 11. Positional analysis for Nt-arginylated sites through ML-based filtering.**
**a, b** DAU test results for Nt-arginylated PSMs that retained through ML-based filtering (**a**) and discarded (**b**).

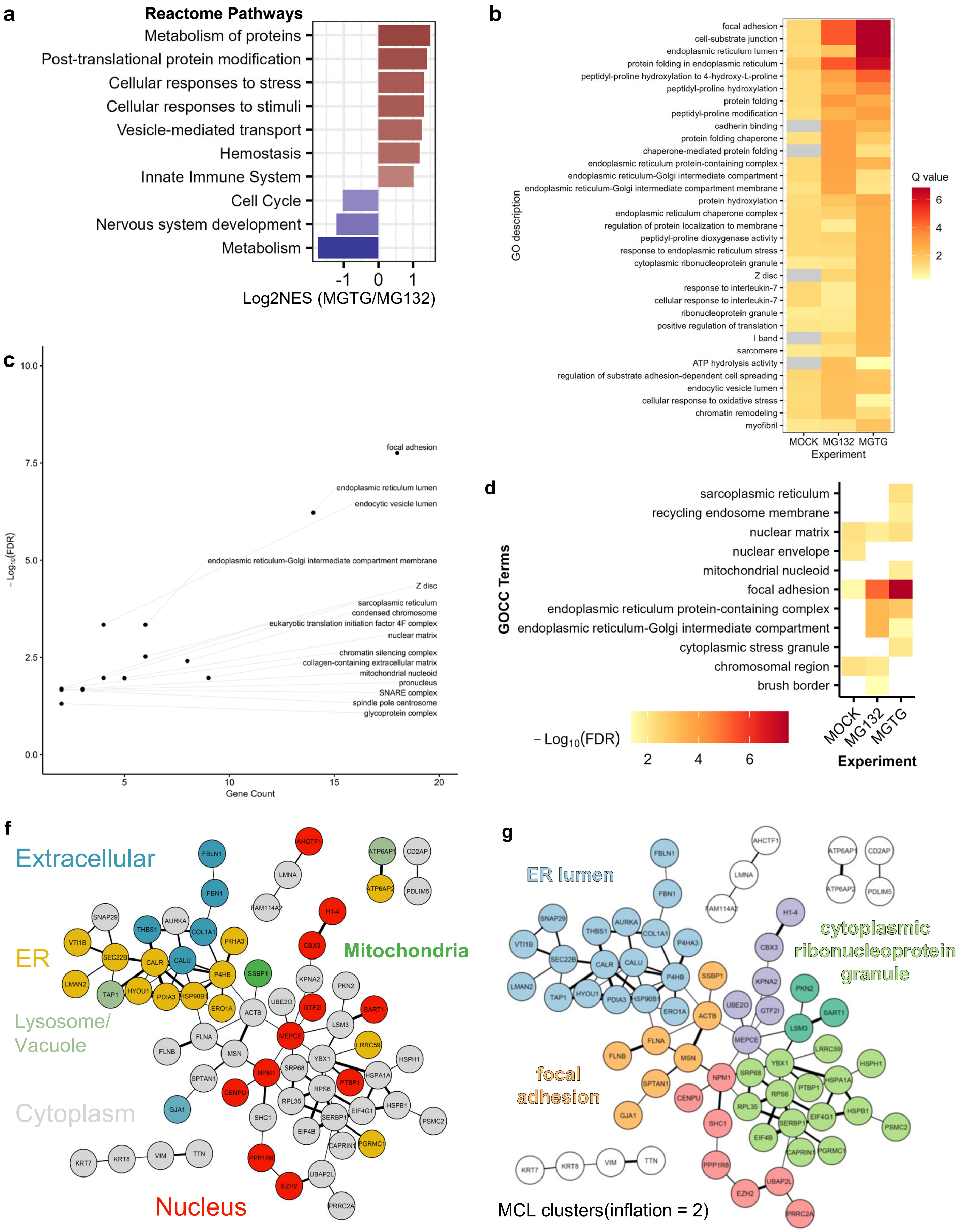
a
Reactome Pathways
Metabolism of proteins
Post-translational protein modification
Cellular responses to stress
Cellular responses to stimuli
Vesicle-mediated transport
Hemostasis
Innate Immune System
Cell Cycle
Nervous system development
Metabolism
-1
0
1
Log2NES (MGTG/MG132)
b
focal adhesion
cell-substrate junction
endoplasmic reticulum lumen
protein folding in endoplasmic reticulum
peptidyl-proline hydroxylation to 4-hydroxy-L-proline
peptidyl-proline hydroxylation
protein folding
peptidyl-proline modification
cadherin binding
protein folding chaperone
chaperone-mediated protein folding
endoplasmic reticulum protein-containing complex
endoplasmic reticulum-Golgi intermediate compartment
endoplasmic reticulum-Golgi intermediate compartment membrane
protein hydroxylation
endoplasmic reticulum chaperone complex
regulation of protein localization to membrane
peptidyl-proline dioxygenase activity
response to endoplasmic reticulum stress
cytoplasmic ribonucleoprotein granule
Z disc
response to interleukin-7
cellular response to interleukin-7
ribonucleoprotein granule
positive regulation of translation
I band
sarcomere
ATP hydrolysis activity
regulation of substrate adhesion-dependent cell spreading
endocytic vesicle lumen
cellular response to oxidative stress
chromatin remodeling
myofibril
GO description
MOCK
MG132
MGTG
Experiment
Q value
6
4
2
c
10.0
7.5
5.0
2.5
0.0
− Log10(FDR)
focal adhesion
endoplasmic reticulum lumen
endocytic vesicle lumen
endoplasmic reticulum-Golgi intermediate compartment membrane
Z disc
sarcoplasmic reticulum
condensed chromosome
eukaryotic translation initiation factor 4F complex
nuclear matrix
chromatin silencing complex
collagen-containing extracellular matrix
mitochondrial nucleoid
pronucleus
SNARE complex
spindle pole centrosome
glycoprotein complex
5
10
15
20
Gene Count
d
sarcoplasmic reticulum
recycling endosome membrane
nuclear matrix
nuclear envelope
mitochondrial nucleoid
focal adhesion
endoplasmic reticulum protein-containing complex
endoplasmic reticulum-Golgi intermediate compartment
cytoplasmic stress granule
chromosomal region
brush border
GOCC Terms
− Log10(FDR)
2
4
6
MOCK
MG132
MGTG
Experiment
f
Extracellular
ER
Lysosome/
Vacuole
Cytoplasm
Mitochondria
Nucleus
FBLN1
FBN1
AHCTF1
LMNA
FAM114A2
ATP6AP1
ATP6AP2
CD2AP
PDLIM5
SNAP29
AURKA
THBS1
COL1A1
P4HA3
H1-4
VTI1B
SEC22B
CALR
CALU
P4HB
SSBP1
CBX3
LMAN2
TAP1
HYOU1
PDIA3
HSP90B1
ERO1A
ACTB
UBE2O
GTF2I
KPNA2
PKN2
SART1
LSM3
FLNA
FLNB
MSN
MEPCE
LRRC59
YBX1
HSPH1
SPTAN1
NPM1
SRP68
HSPA1A
PTBP1
RPS6
GJA1
CENPU
RPL35
SHC1
EIF4G1
HSPB1
SERBP1
PSMC2
EIF4B
CAPRIN1
PGRMC1
KRT7
KRT8
VIM
TTN
PPP1R8
UBAP2L
EZH2
PRRC2A
g
ER lumen
cytoplasmic
ribonucleoprotein
granule
focal
adhesion
MCL clusters(inflation = 2)

**Supplementary Figure 12. Characterization of Nt-arginylome.**

**a,** Reactome pathways enriched by GSEA using $log_2$ fold changes of Nt-arginylation sites comparing MGTG to MG132. NES, normalized enrichment score. **b,** ORA on GO database using gene names of Nt-arginylome. **c,** GO Cellular Compartment (CC) terms in GO analysis result at figure **b** are plotted. **d,** GOCC terms by ORA of proteins with Nt-arginylation. **e,** STRING analysis of Nt-arginylation sites. Protein is depicted by colored circles, with colors indicating their predicted subcellular localization. The confidence in the physical network is visually represented by the varying thickness of the connecting edges. **f**, MCL clustering result from panel **e** are depicted by color, wieht each color corresponding to the most significantly enriched GOCC terms for that cluster.

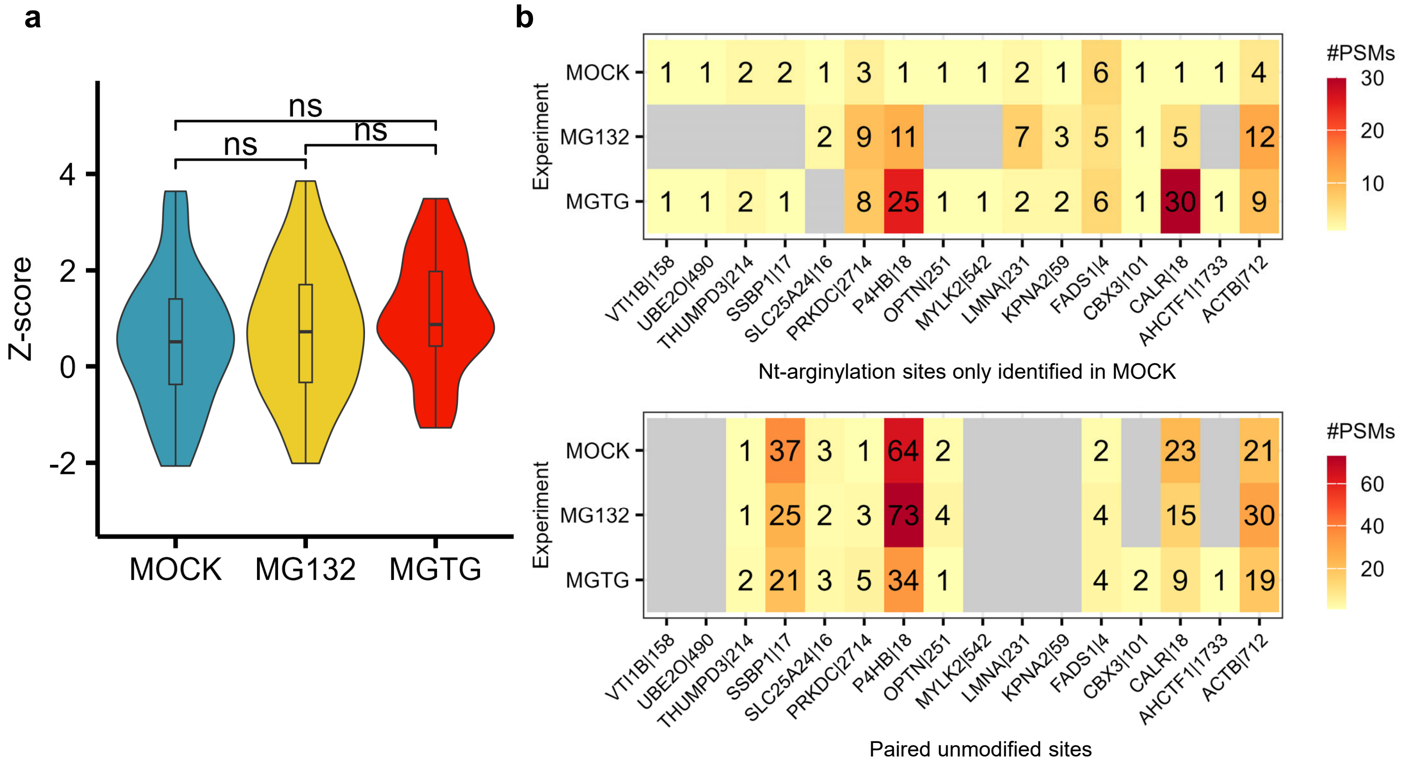


**Supplementary Figure 13. Quantitative analysis of Nt-arginylation sites discovered in MOCK.**
**a,** Violin plots of average LFQ intensities for each Nt-arginylation sites discovered in MOCK experiment across various experimental conditions. A two-tailed Student's t-test was used to analze results. **b,** PSM count of Nt-arginylation sites and their paired unmodified sites discovered in MOCK across all conditions.

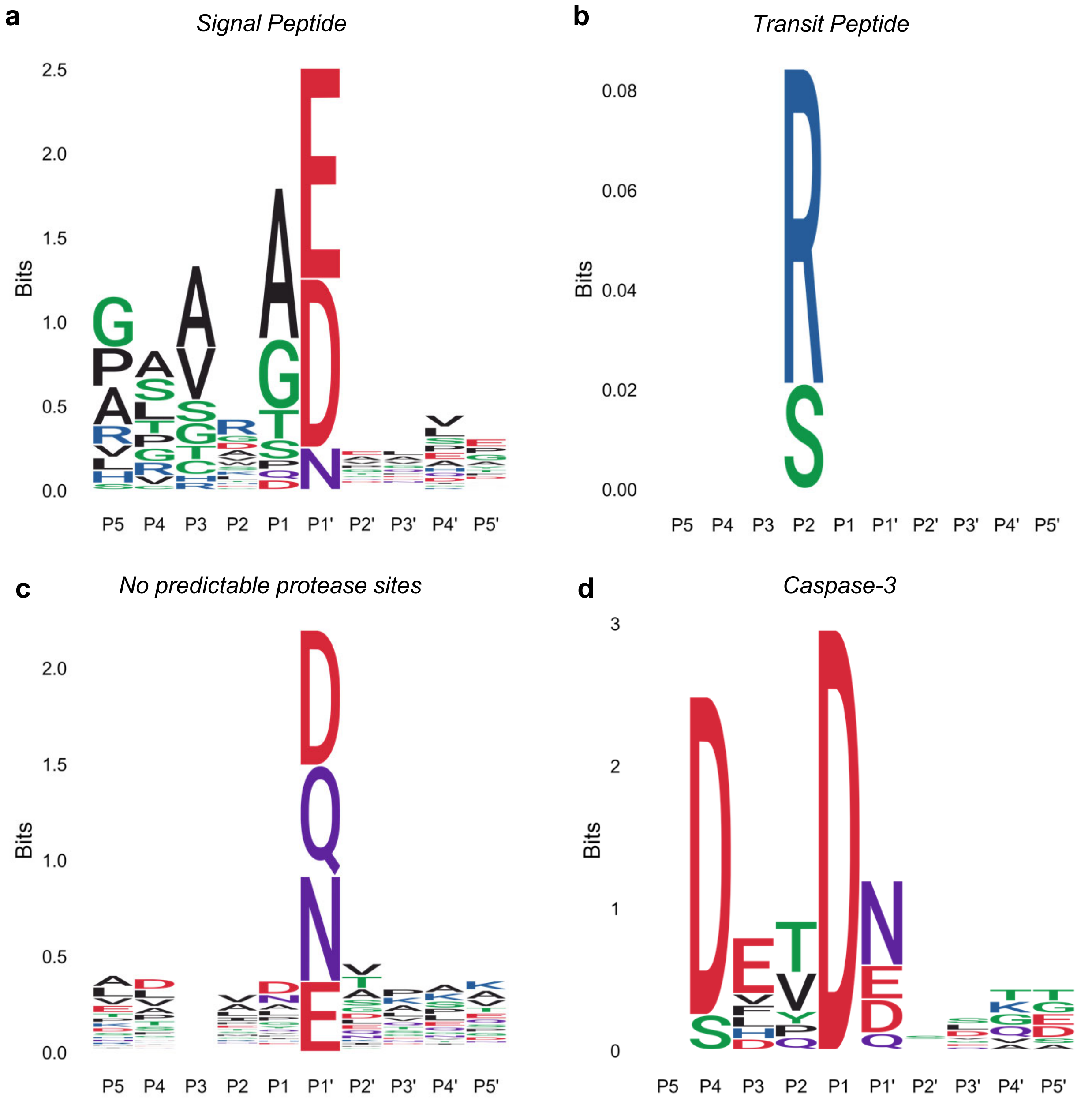


**Supplementary Figure 14. Logo analysis of Nt-arginylome based on the motifs for proteases**
The sequences of Nt-arginylation sites were analyzed for predicting targeting proteases using SignalP (**a**), TargetP (**b**), and Procleave (**c**,**d**).

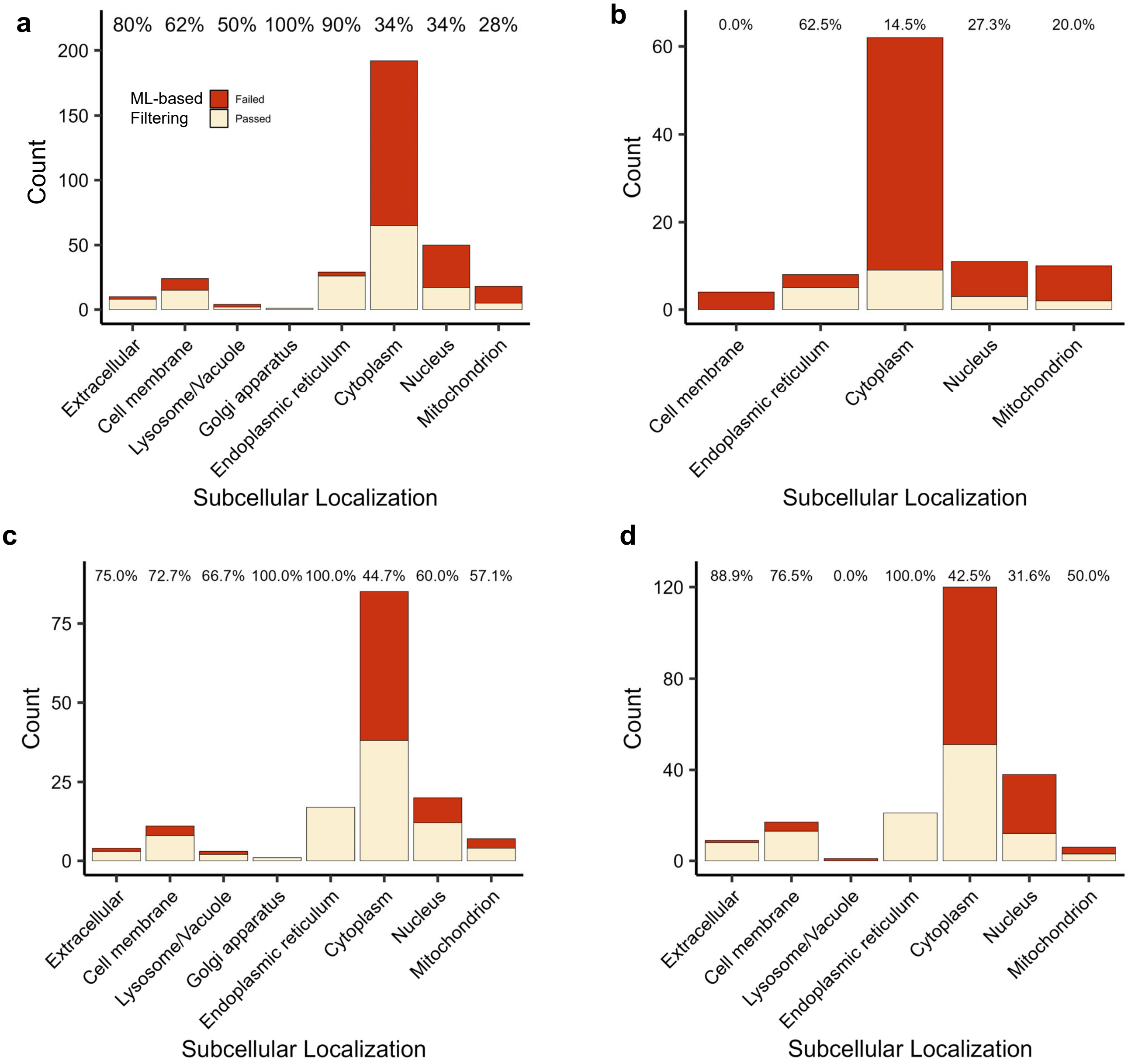


**Supplementary Figure 15. Subcellular localization prediction of Nt-arginylome.**
Predicted subcellular localization for the putative Nt-arginylation sites in experiments of all (**a**), MOCK (**b**), MG132 (**c**), and MGTG (**d**). The count indicates the number of discarded and retained Nt-arginylation sites through ML-based filtering. Percentages represent the proportion of retained sites per each localization.

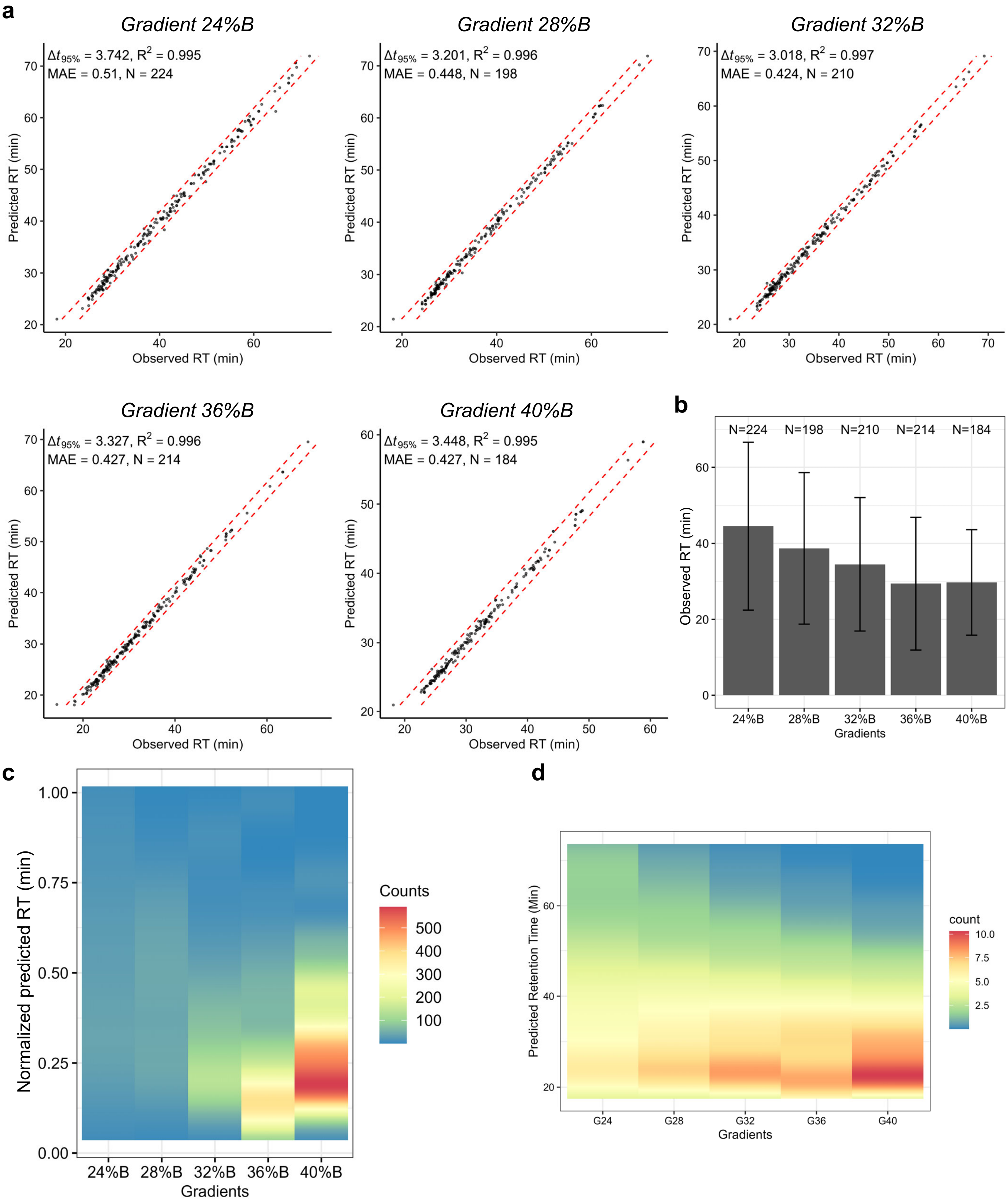


**Supplementary Figure 16. Optimal gradient determined using RT prediction models**
**a,** Performance of RT prediction models using Arg-starting peptides in training set across various LC gradient setups. **b,** Average observed RT of Arg-starting peptides identified in each gradients. Error bar indicates a standard deviation. **c,** Density of normalized predicted RT of Arg-starting peptides across each RT prediction models which were trained for specific gradients. **d,** Density of predicted RT of target Nt-arginylated peptides across each RT prediction models.

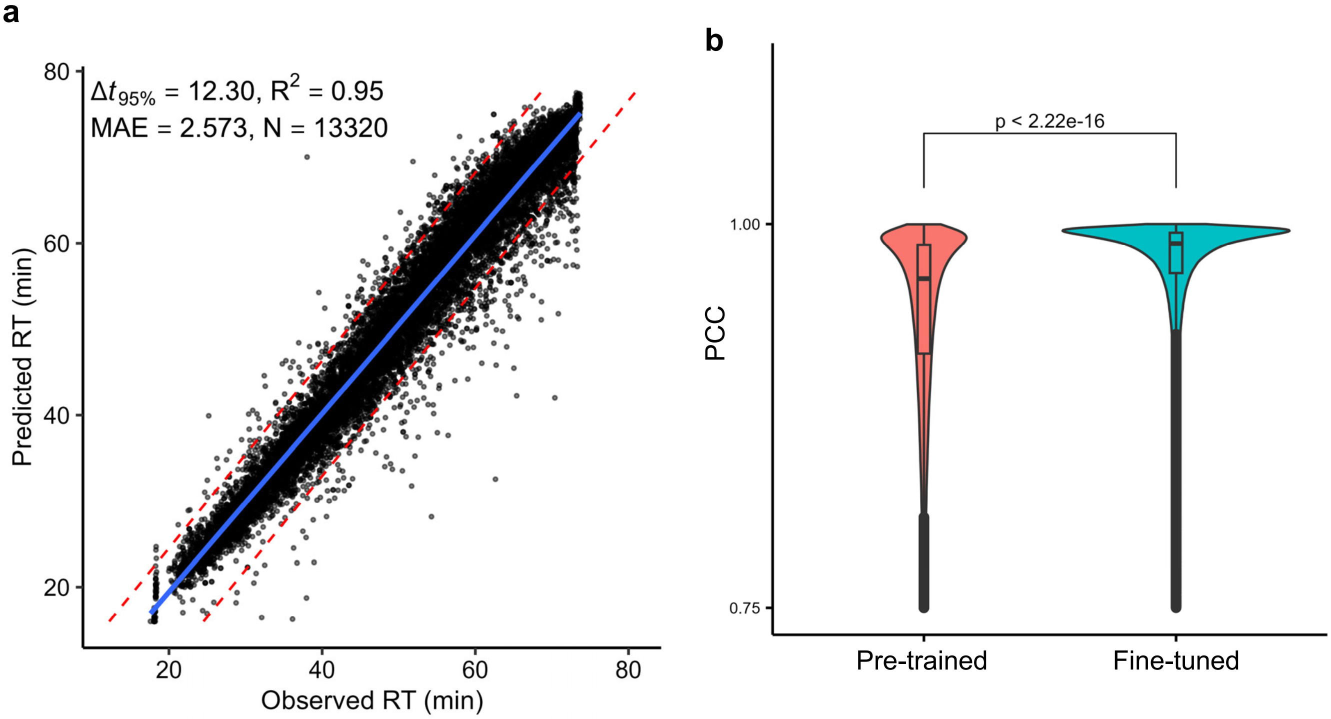


**Supplementary Figure 17. Evaluation of training methods for RT and MS2 prediction model.**
**a,** RT prediction model trained without transfer learning method, using PSMs of HeLa digest profiling data. **b,** Comparison of MS2 prediction model performance with or without transfer learning method, using the same set of PSMs in panel **a**. PCCs were obtained by testing the MS2 prediction model with PSMs used for training the MS2 models. A two-tailed Student's t-test was used to analyze results.

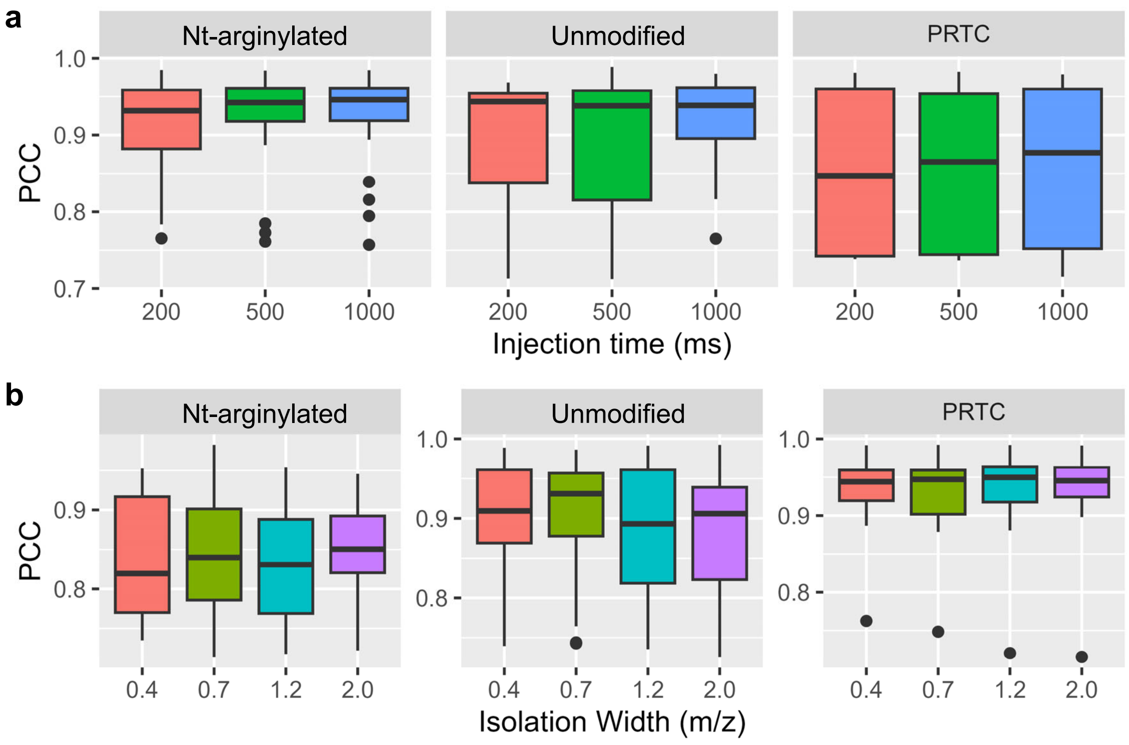


**Supplementary Figure 18. Optimization of ion injection time and isolation width during PRM-MS.**
Optimization results for MS acquisition parameters: injection time (**a**), and isolation width (**b**). PCC values of target peptides obtained by comparing experimental MS2 spectra with predicted MS2 spectra from fine-tuned MS2 prediction models. PRTC refers to peptides of Pierce Retention Time Calibration (PRTC) mixtures.

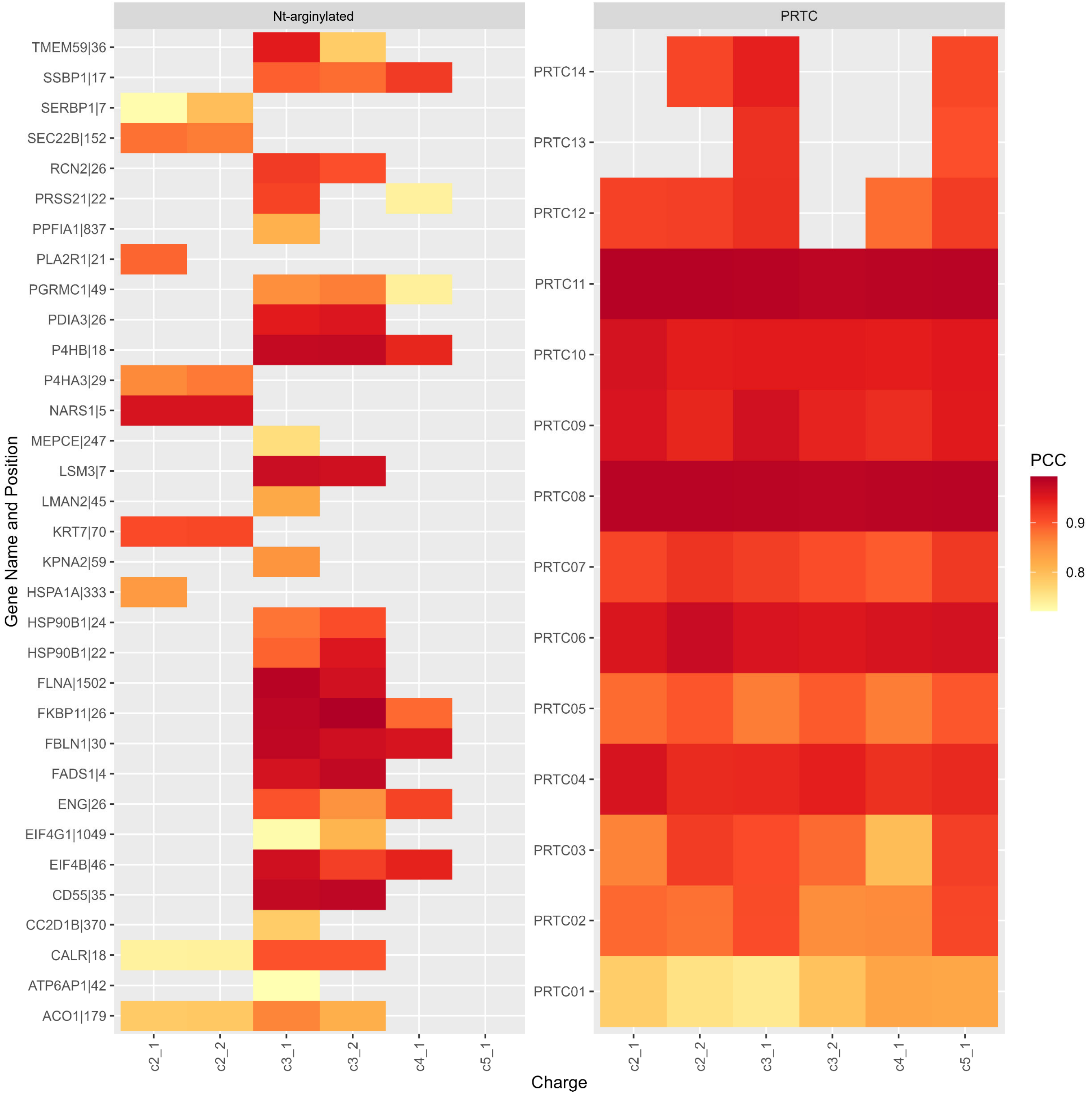


**Supplementary Figure 19. Optimization of charge for target peptides during PRM-MS.**
Heatmap showing PCC values of target peptides, calculated by comparing predicted MS2 spectra from the fine-tuned MS2 prediction models. c2, charge +2; c3, charge +3; c4, charge +4; c5, charge +5.

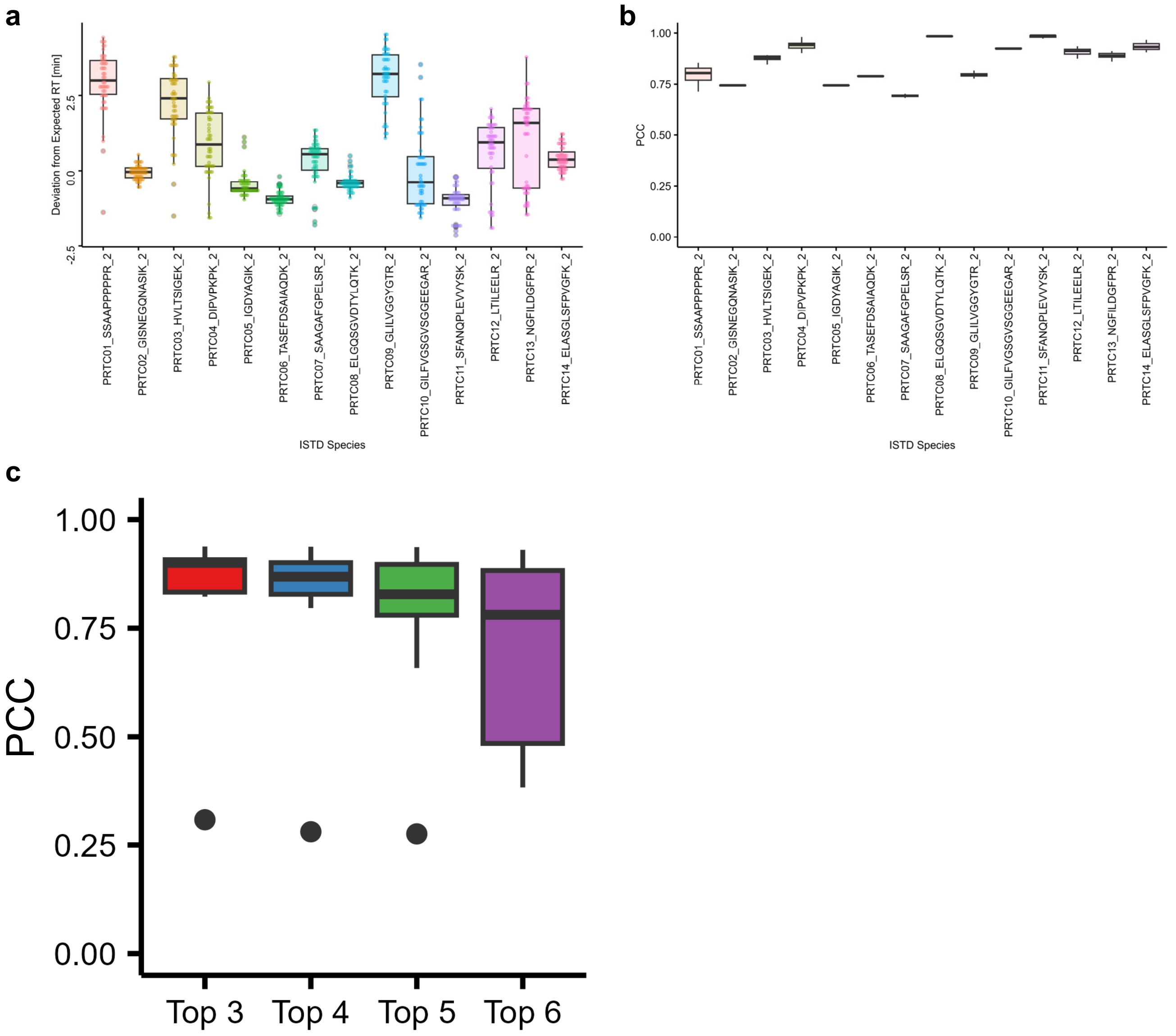


**Supplementary Figure 20. Optimzations of PRM-MS on MS2 matching between the observed and the predicted spectrum.**

**a,b** Comparative box plots showing the deviation in retention time (**a**) and PCC result (**b**) of fragment spectra comparison between predicted and actual ones for PRTC peptides. **c,** Pearson's correlation coefficient (PCC) between LFQ and integrated intensity of Top N matched fragment ions with predicted MS2 spectra.

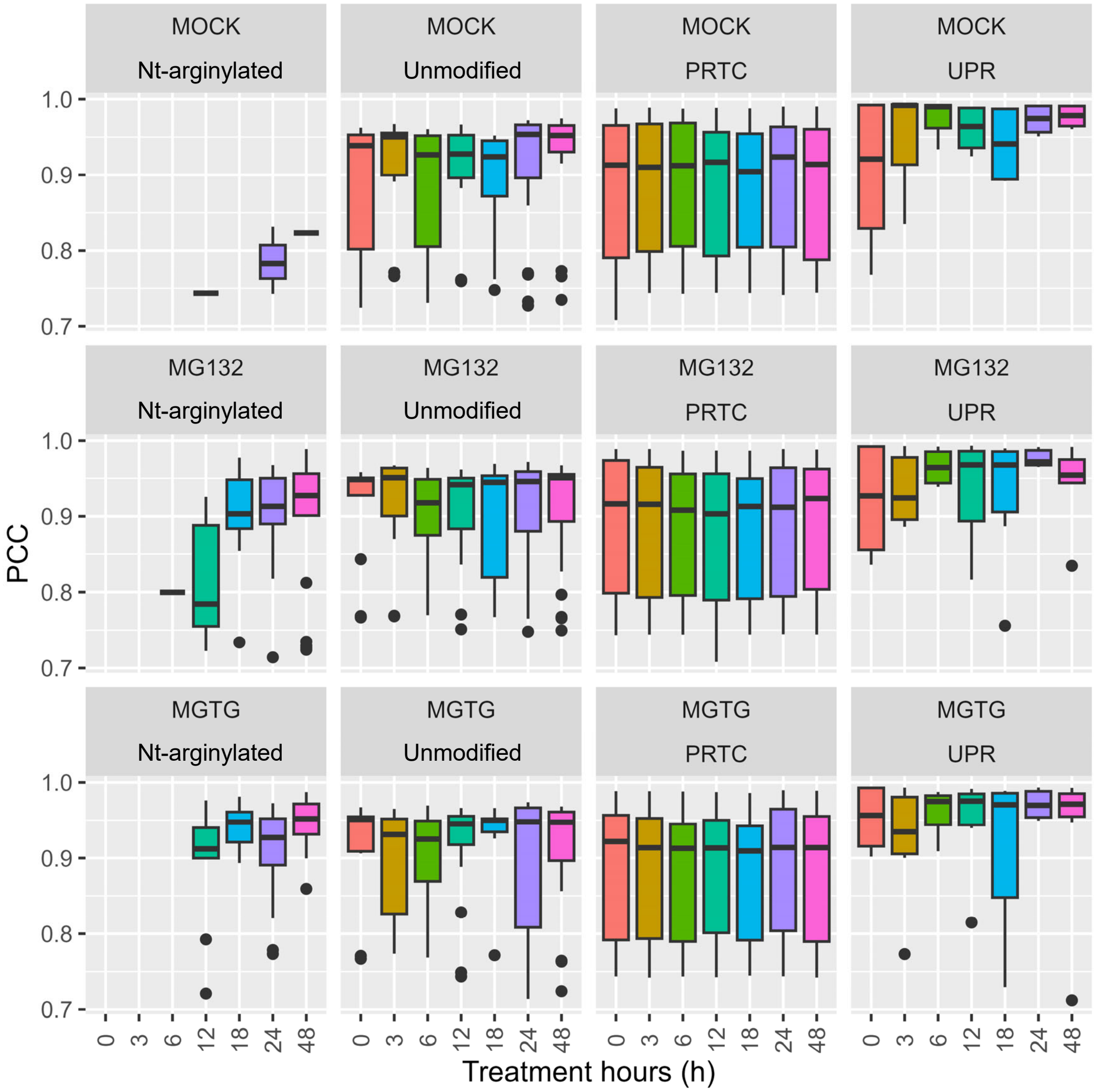


**Supplementary Figure 21. Identification result of PRM-MS experiments.**
PCC values of target peptides obtained by calculating with predicted MS2 spectra from the fine-tuned MS2 prediction. PRTC, peptides of pierce retention time calibration (PRTC) mixtures; UPR, target peptides of indicator proteins of UPR pathway activation.

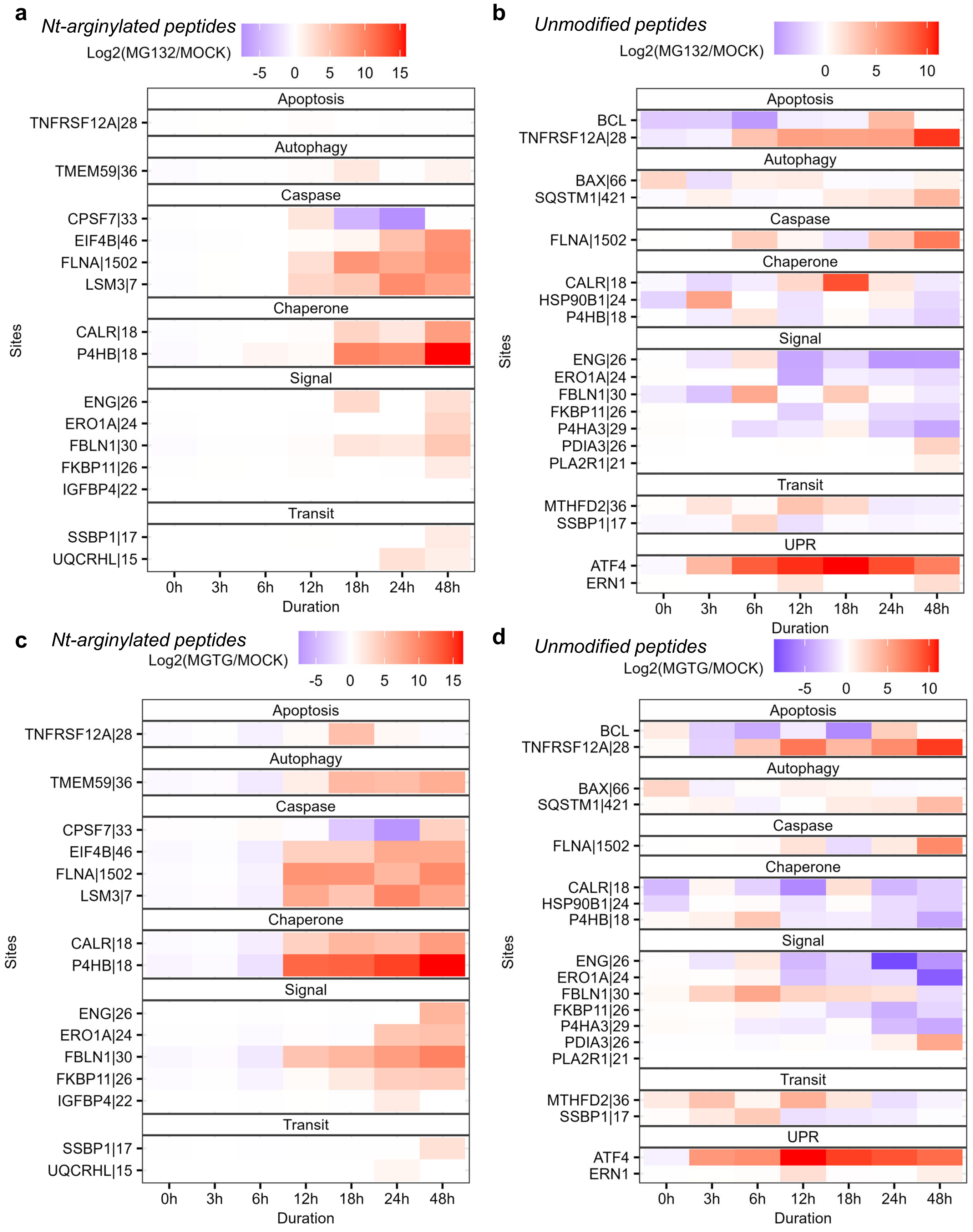


**Supplementary Figure 22. Quantitation result of PRM-MS experiments.**
Average of normalized intensity values of target peptides are shown in heatmap. Apoptosis and autophagy, proteins have significant role with reference in each pathway; caspase, sites predicted caspase-3 substrate; chaperone, proteins with biological function of chaperone; signal, sites known as signal peptide cleavage sites; transit, sites known as transit peptide cleavage sites; UPR, target peptides of indicator proteins of UPR pathway activation. **a,b** Result of MG132 treatment with Nt-arginylated peptides (**a**) and unmodified peptides (**b**). **c,d** Result of MGTG treatment with Nt-arginylated peptides (**c**) and unmodified peptides (**d**).

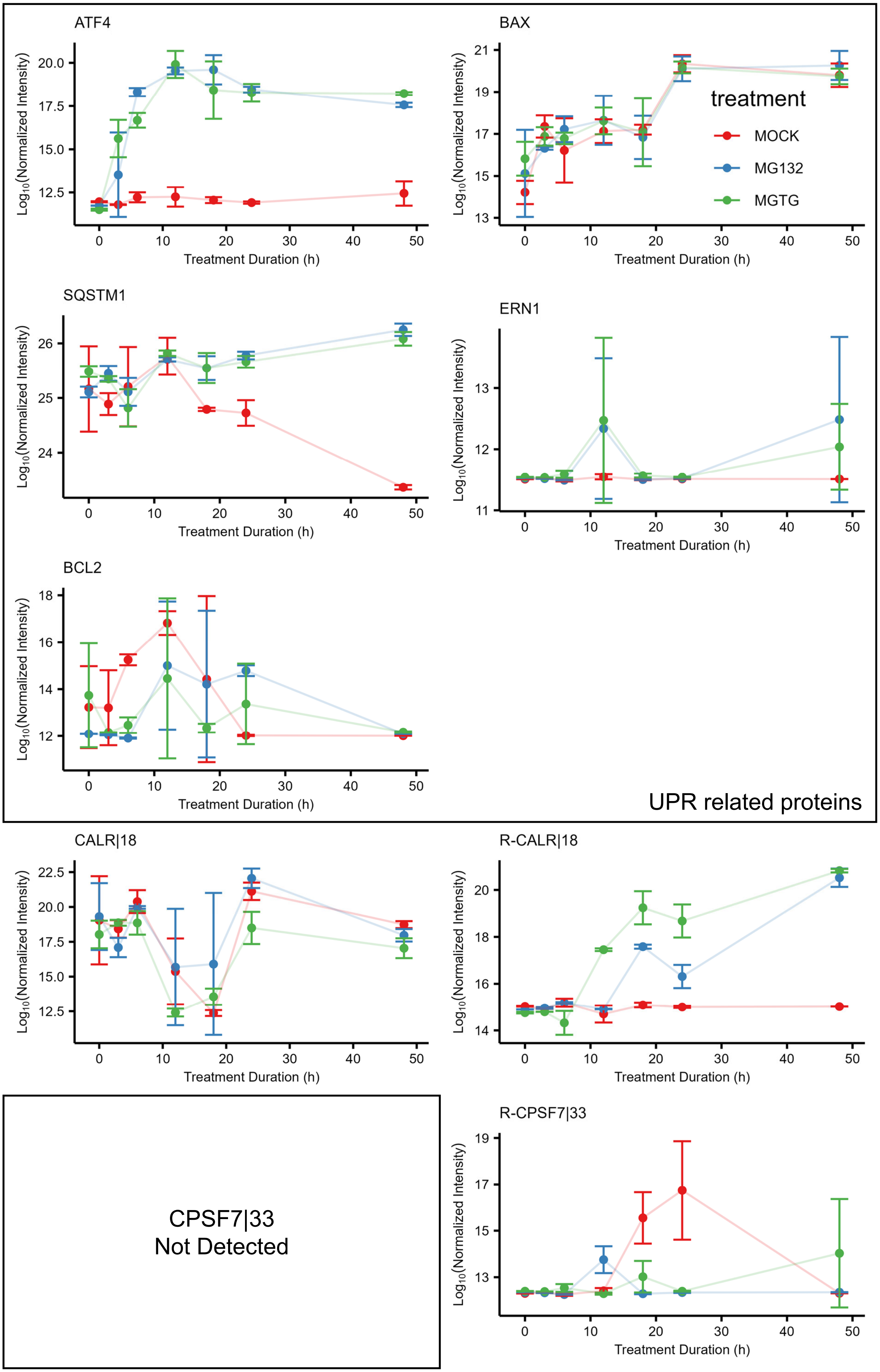
ATF4
BAX
SQSTM1
ERN1
BCL2
UPR related proteins
CALR|18
R-CALR|18
CPSF7|33
Not Detected
R-CPSF7|33
treatment
MOCK
MG132
MGTG
Log10(Normalized Intensity)
Treatment Duration (h)

EIF4B|46
Not Detected

R-EIF4B|46

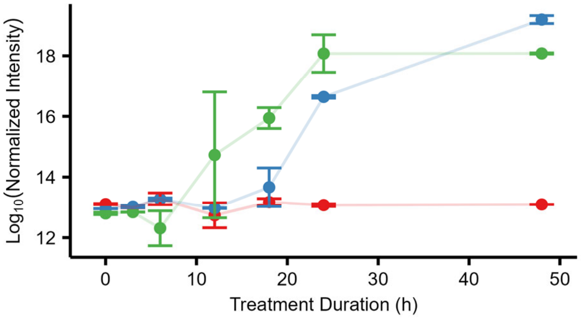


ERO1A|24

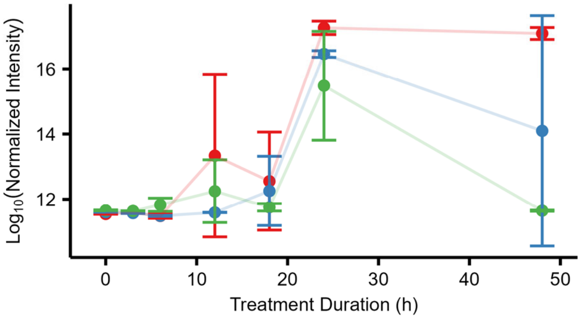


R-ERO1A|24

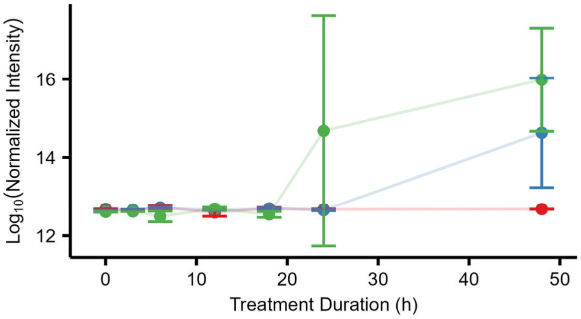


FBLN1|30

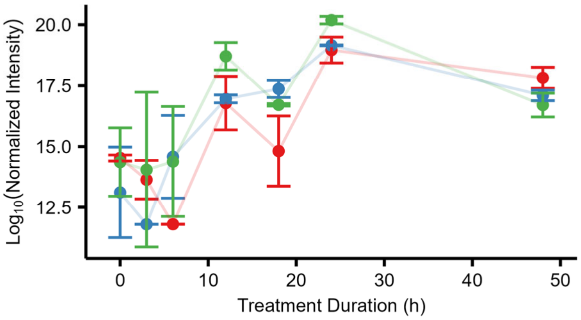


R-FBLN1|30

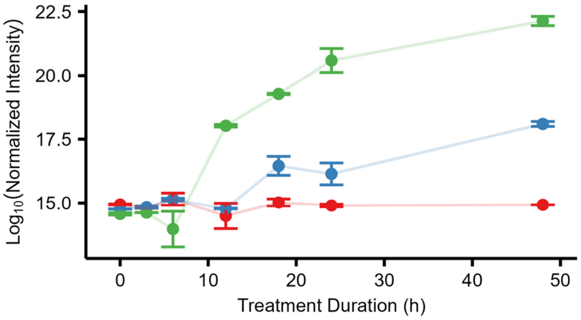


FKBP11|26

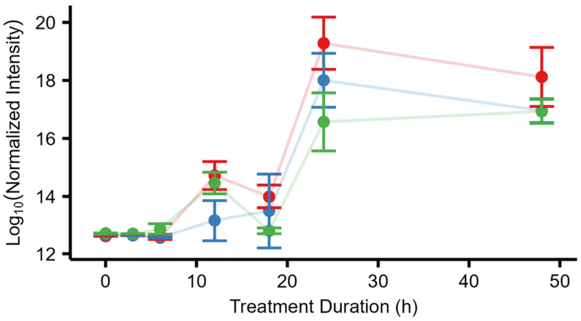


R-FKBP11|26

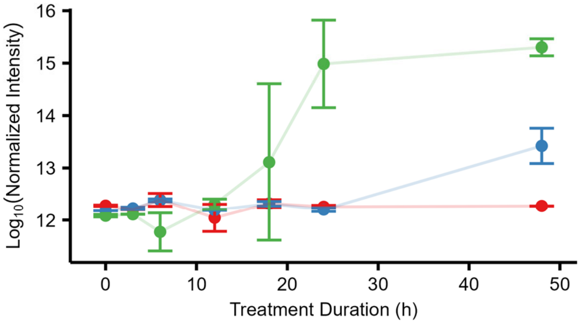


FLNA|1502

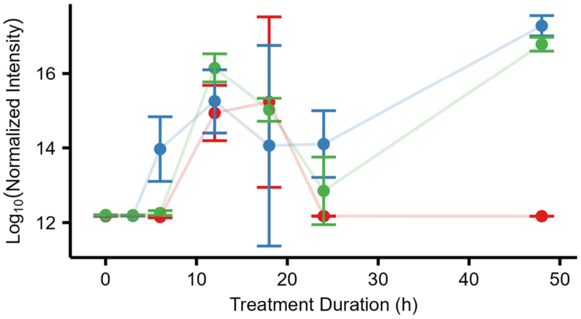


R-FLNA|1502

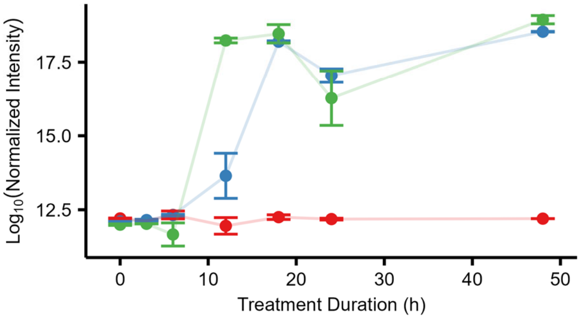

HSP90B1|24

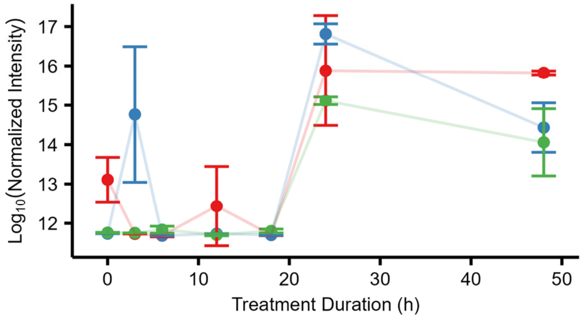


R-HSP90B1|24
Not Detected

IGFBP4|22
Not Detected

R-IGFBP4|22

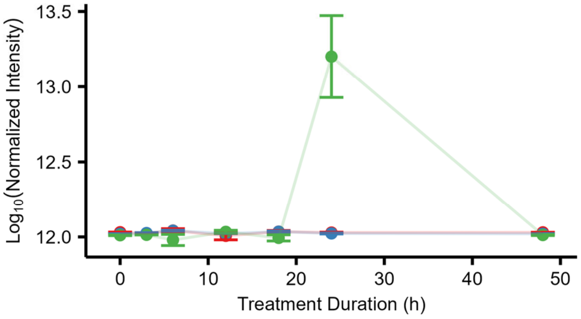


LSM3|22
Not Detected

R-LSM3|7

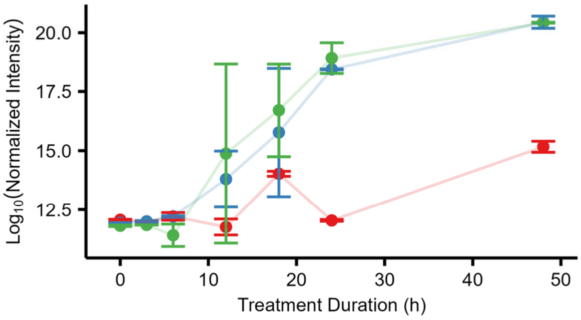


MTHFD2|36

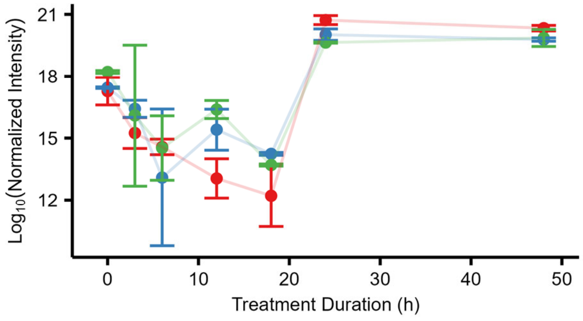


R-MTHFD2|36
Not Detected

P4HB|18

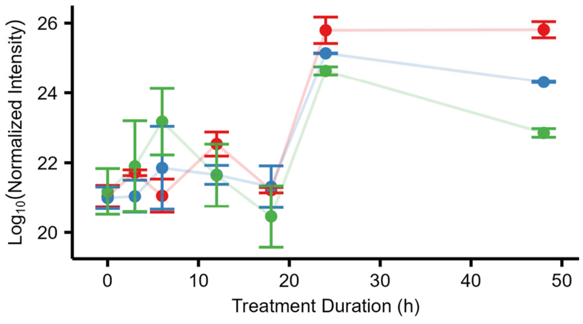


R-P4HB|18

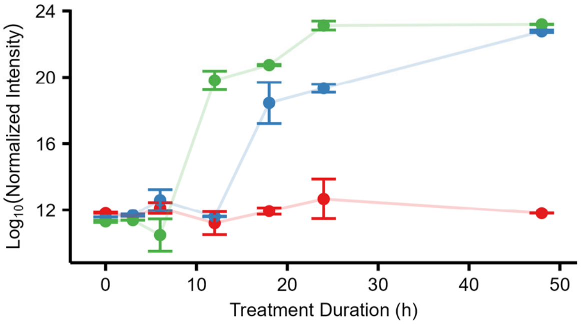

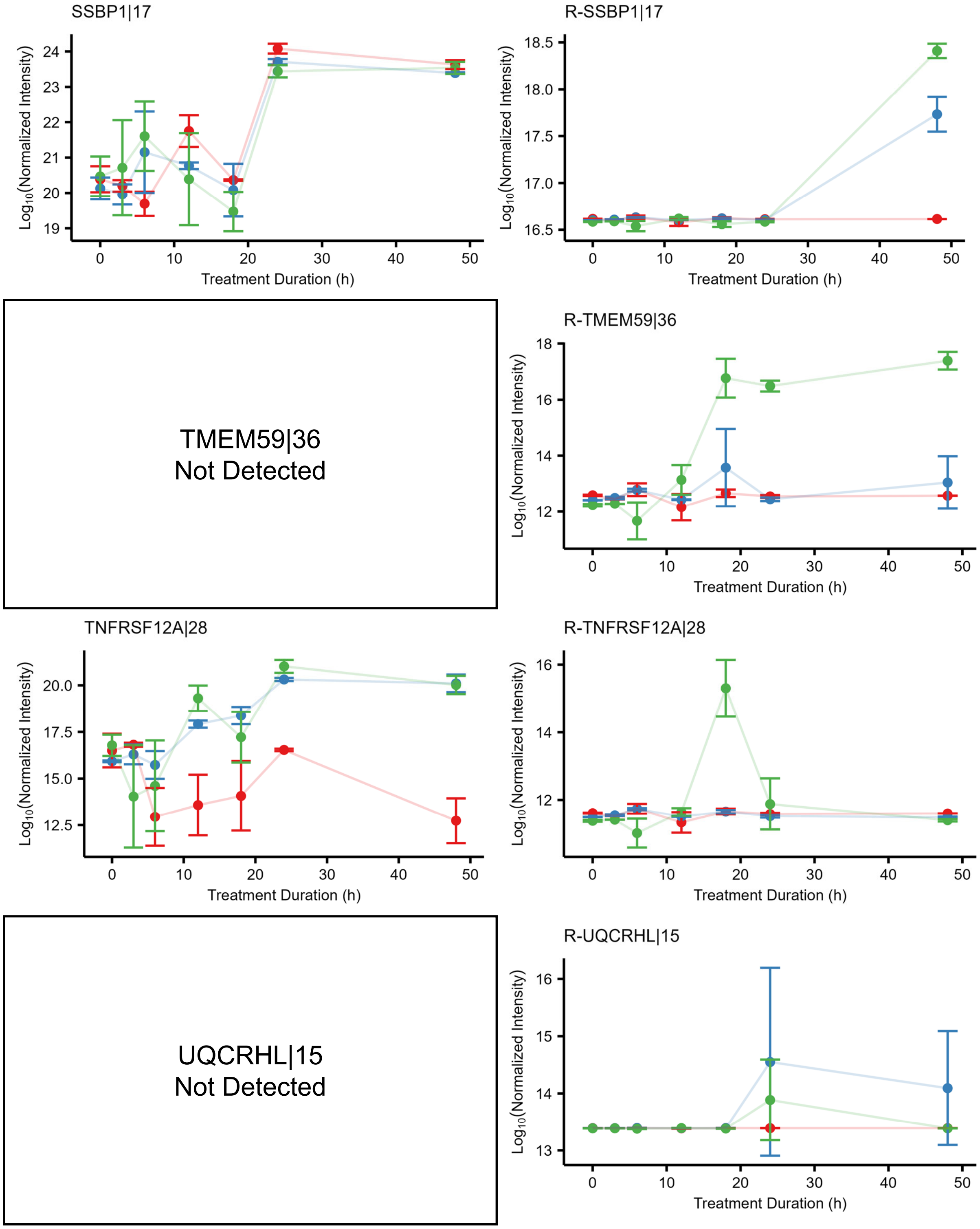


**Supplementary Figure 23. Quantitation result of each target sites.**

Time-course data of average of normalized intensity values of target peptides. Identifiers indicates: (Gene name of protein)|(position). R-, Nt-arginylation. The plots are arranged from left to right, by the presence of N-terminal modification. The plot area was left vacant when we failed to observe any sites related to either Nt-arginylated peptides or unmodified peptides.

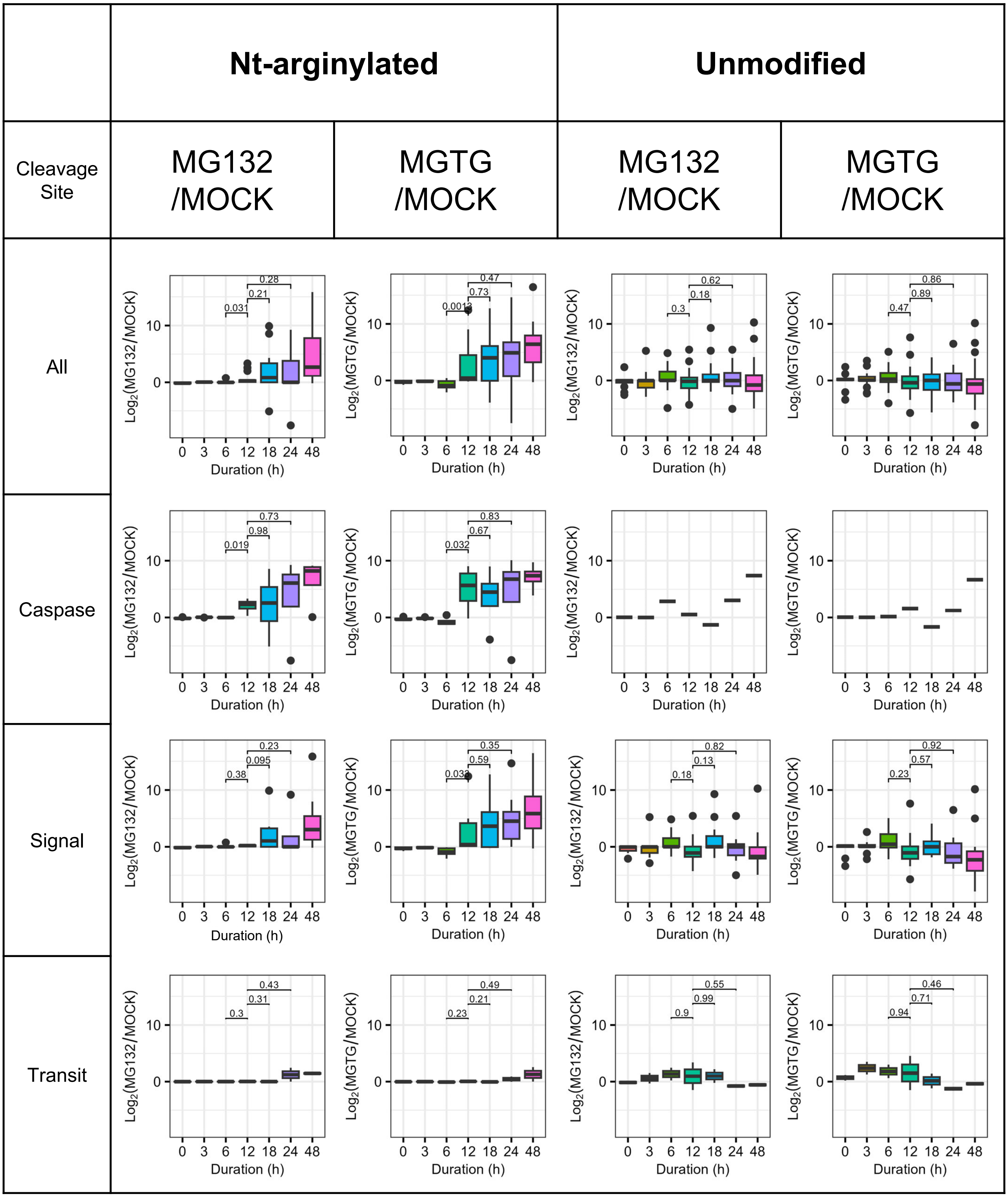


**Supplementary Figure 24. PRM quantification result**
Quantification of Nt-arginylated and unmodified peptides are categorized by protease cleavage sites and treatment group.

Immunoblot images for DMSO

ATF4

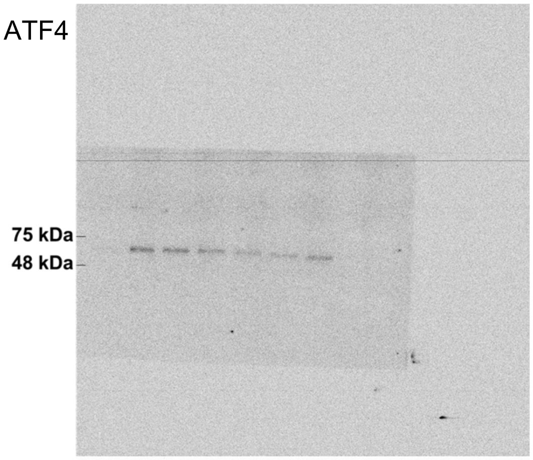


CASP3

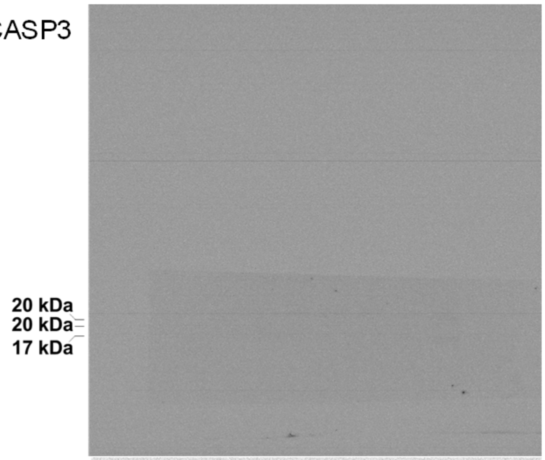


HSPA5

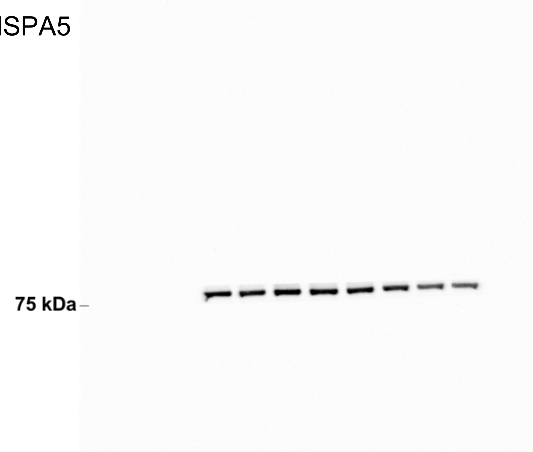


R-HSPA5

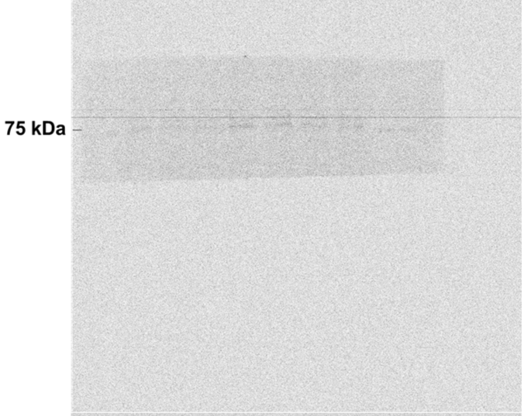


CALR

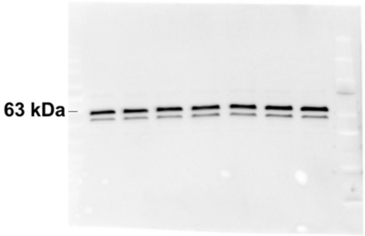


R-CALR

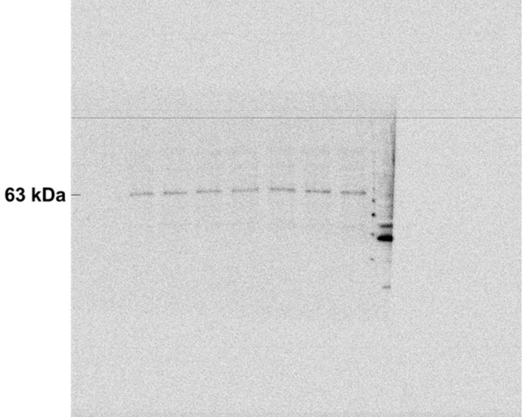


P4HB

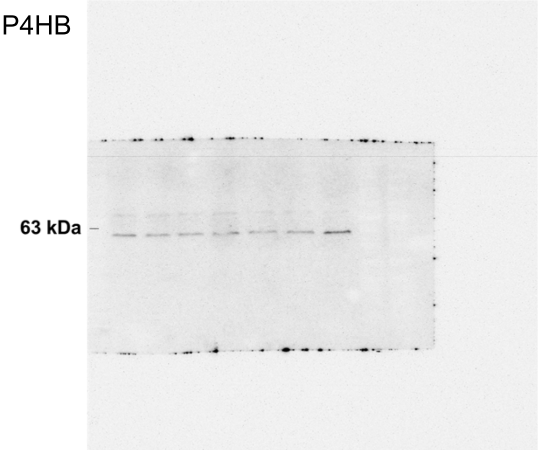


R-P4HB

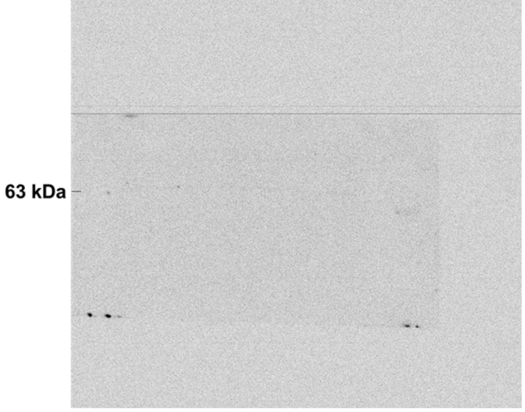

Immunoblot images for MG132

ATF4

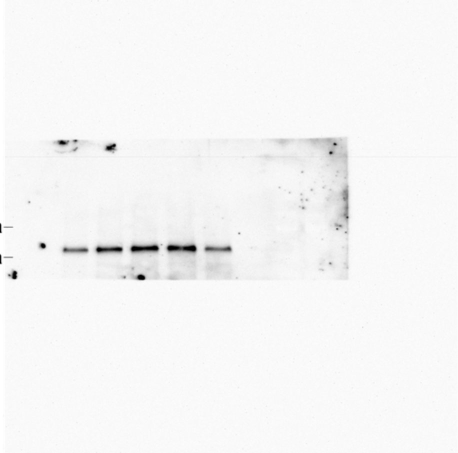


CASP3

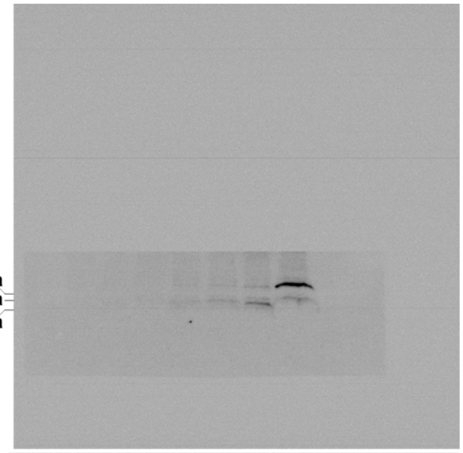


HSPA5

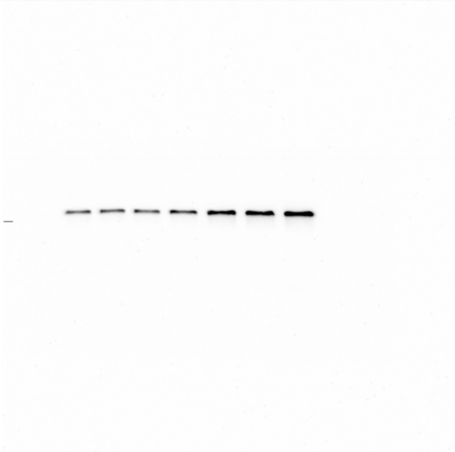


R-HSPA5

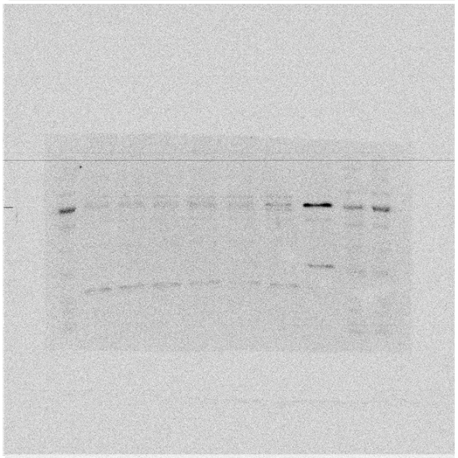


CALR

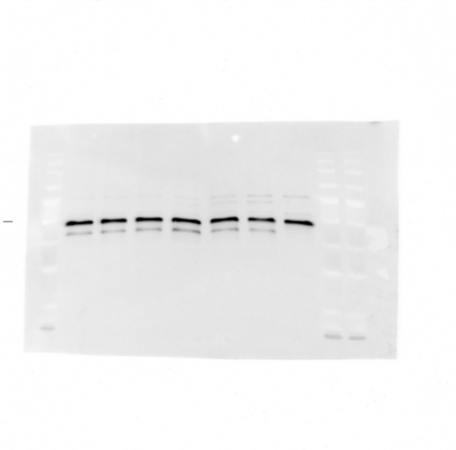


R-CALR

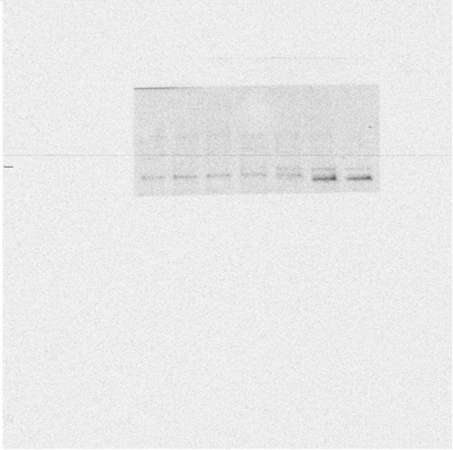


P4HB

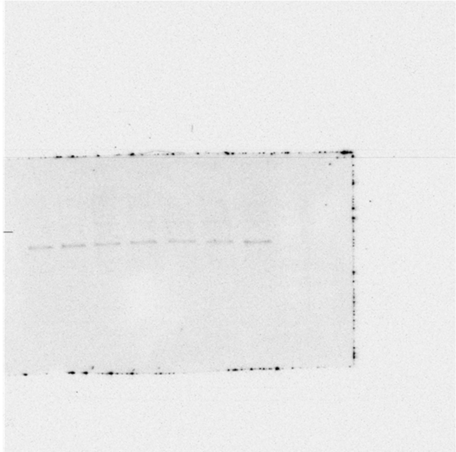


R-P4HB

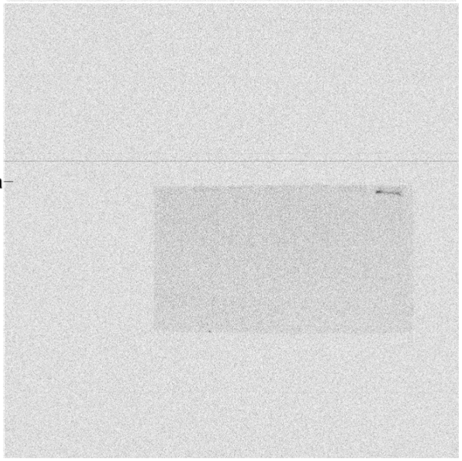

Immunoblot images for MGTG

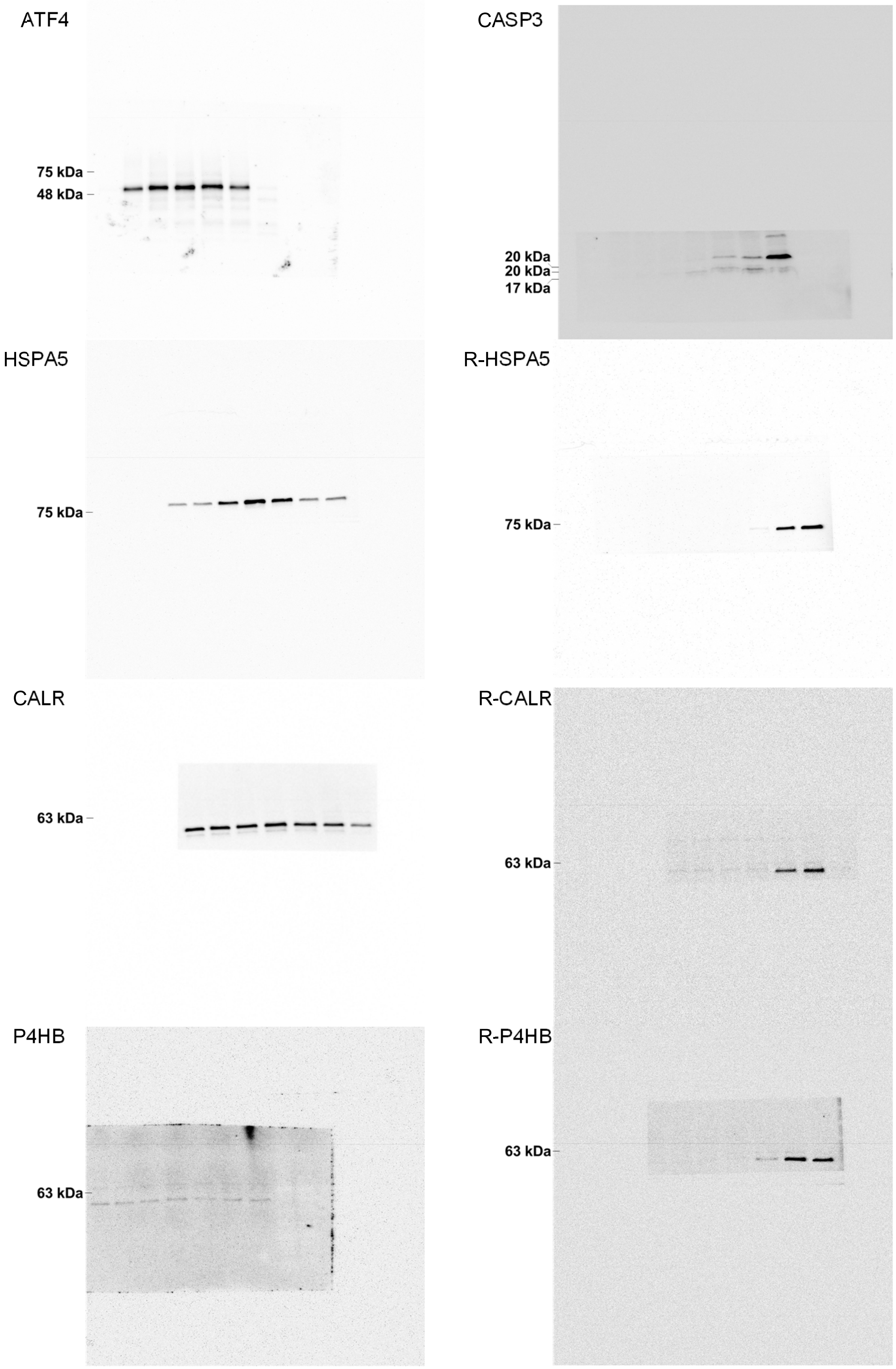


**Supplementary Figure 25. Uncropped scans of immunoblots for figure 6k.**

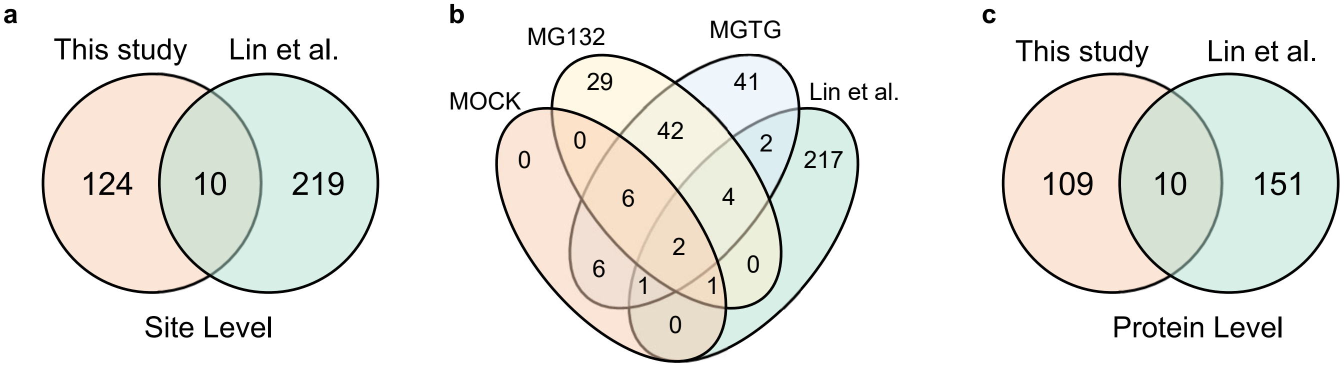


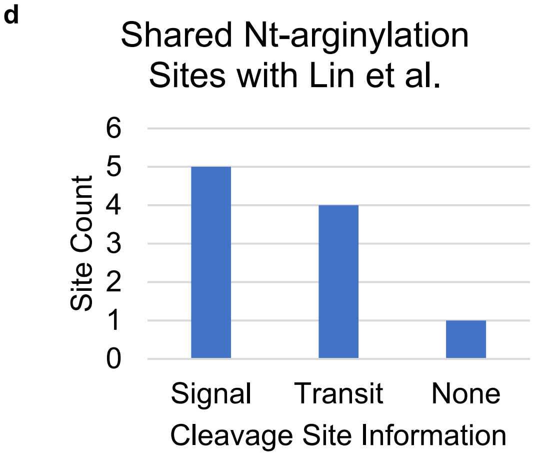


| Nt-arginylation Site | PSM count | SITEID in Lin et al. | cleavage prediction |
|---|---|---|---|
| PGRMC1\|O00264\|PGRC1_HUMAN\|49 | 1 | PGRC1_49 | |
| P4HB\|P07237\|PDIA1_HUMAN\|18 | 37 | PDIA1_18 | Signal |
| MTHFD2\|P13995\|MTDC_HUMAN\|36 | 2 | MTDC_36 | Transit |
| HSP90B1\|P14625\|ENPL_HUMAN\|22 | 64 | ENPL_22 | Signal |
| CALR\|P27797\|CALR_HUMAN\|18 | 36 | CALR_18 | Signal |
| SHMT2\|P34897\|GLYM_HUMAN\|27 | 2 | GLYM_27 | Transit |
| SSBP1\|Q04837\|SSBP_HUMAN\|17 | 3 | SSBP_17 | Transit |
| LMAN2\|Q12907\|LMAN2_HUMAN\|45 | 4 | LMAN2_45 | Signal |
| SLC25A24\|Q6NUK1\|SCMC1_HUMAN\|16 | 3 | SCMC1_16 | Transit |
| ERO1A\|Q96HE7\|ERO1A_HUMAN\|24 | 6 | ERO1A_24 | Signal |

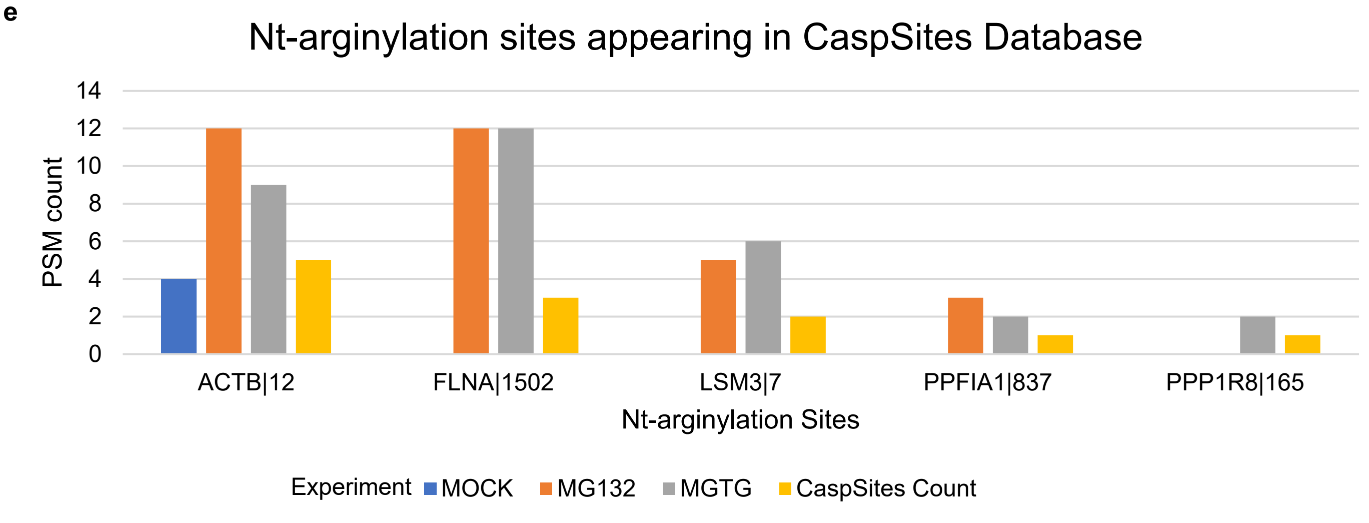


**Supplementary Figure 26. Comparison of discovered Nt-arginylome with the data from Lin et al.**
**a,b** Venn diagram of shared Nt-arginylation sites between this study and Lin et al. The sites discovered in this study are further categorized by experiment in figure **b**. **c,** Shared proteins with Nt-arginylation sites between this study and Lin et la. **d,** Analysis of shared Nt-arginylation sites by cleavage prediction. Table shows a summary of shared Nt-arginylation sites. **e,** Shared Nt-arginylation sites that are annotated in the CaspSites database.

**Supplementary Table 1. List of Nt-arginylome**

| Gene\|Uniprot Accession\|Uniprot Name\|Position | Sequence | cleavage_type | Predicted Localization |
|---|---|---|---|
| ABCD1\|P33897\|ABCD1_HUMAN\|424 | QVFEDVQR | | Cytoplasm |
| ACO1\|P21399\|ACOHC_HUMAN\|179 | QVNLEYLAR | | Cytoplasm |
| ACTB\|P60709\|ACTB_HUMAN\|12 | NGSGMCKAGFAGDDAPR | caspase-1 | Cytoplasm |
| ACTB\|P60709\|ACTB_HUMAN\|280 | NSIMKCDVDIR | cathepsin D | Endoplasmic reticulum |
| AHCTF1\|Q8WYP5\|ELYS_HUMAN\|1733 | DVVSSKTR | | Cytoplasm |
| ATP6AP1\|Q15904\|VAS1_HUMAN\|42 | EQQVPLVLWSSDR | Signal Pepitde | Cytoplasm |
| ATP6AP2\|O75787\|RENR_HUMAN\|282 | EAKQAKNPASPY | | Cytoplasm |
| AURKA\|O14965\|AURKA_HUMAN\|35 | NPLPVNSGQAQR | | Cytoplasm |
| BSG\|P35613\|BASI_HUMAN\|161 | DSATEVTGHR | | Lysosome/Vacuole |
| CALR\|P27797\|CALR_HUMAN\|18 | EPAVYFKEQFLDGDGWTSR | Signal Pepitde | Cytoplasm |
| CALU\|O43852\|CALU_HUMAN\|225 | DEPEWVKTER | | Cytoplasm |
| CAPRIN1\|Q14444\|CAPR1_HUMAN\|101 | EVTNNLEFAKELQR | | Nucleus |
| CAST\|P20810\|ICAL_HUMAN\|77 | DAHNKKAVSR | | Cytoplasm |
| CBLL1\|Q75N03\|HAKAI_HUMAN\|6 | NELQGTNSSGSLGGLDVR | caspase-3 | Cytoplasm |
| CBX3\|Q13185\|CBX3_HUMAN\|101 | DSKSKKKR | | Nucleus |
| CC2D1B\|Q5T0F9\|C2D1B_HUMAN\|370 | DVPATPVAPTESQTVLDALQQR | | Endoplasmic reticulum |
| CD2AP\|Q9Y5K6\|CD2AP_HUMAN\|577 | DVKKNSLDELR | caspase-5 | Endoplasmic reticulum |
| CD55\|P08174\|DAF_HUMAN\|35 | DCGLPPDVPNAQPALEGR | Signal Pepitde | Endoplasmic reticulum |
| CENPU\|Q71F23\|CENPU_HUMAN\|49 | NSDVSSIGR | caspase-3 | Cytoplasm |
| CFDP1\|Q9UEE9\|CFDP1_HUMAN\|44 | QTQKTQGKKR | | Cytoplasm |
| COL1A1\|P02452\|CO1A1_HUMAN\|24 | EEGQVEGQDEDIPPITCVQNGLR | Signal Pepitde | Cytoplasm |
| CPSF7\|Q8N684\|CPSF7_HUMAN\|33 | DVLTATSQPSDDR | caspase-3 | Extracellular |
| DGCR2\|P98153\|IDD_HUMAN\|22 | EPLRPELR | Signal Pepitde | Endoplasmic reticulum |
| EIF4B\|P23588\|IF4B_HUMAN\|46 | DLEGDVSTTWHSNDDDVYR | caspase-3 | Endoplasmic reticulum |
| EIF4G1\|Q04637\|IF4G1_HUMAN\|1049 | DGGWNTVPISKGSRPIDTSR | | Cytoplasm |
| ENG\|P17813\|EGLN_HUMAN\|26 | ETVHCDLQPVGPER | Signal Pepitde | Cell membrane |
| ERO1A\|Q96HE7\|ERO1A_HUMAN\|24 | EEQPPETAAQR | Signal Pepitde | Cytoplasm |
| EZH2\|Q15910\|EZH2_HUMAN\|209 | DKESRPPR | | Cytoplasm |
| FADS1\|O60427\|FADS1_HUMAN\|4 | DPVAAETAAQGPTPR | LAST_MAM peptidase | Cytoplasm |
| FADS1\|O60427\|FADS1_HUMAN\|9 | ETAAQGPTPR | | Cytoplasm |
| FAM114A2\|Q9NRY5\|F1142_HUMAN\|35 | QGAKPESKSEPVVSTR | | Nucleus |
| FBLN1\|P23142\|FBLN1_HUMAN\|30 | DVLLEACCADGHR | Signal Pepitde | Cytoplasm |
| FBN1\|P35555\|FBN1_HUMAN\|28 | NLEAGNVKETR | Signal Pepitde | Mitochondrion |
| FKBP11\|Q9NYL4\|FKB11_HUMAN\|26 | EAGLETESPVR | Signal Pepitde | Cytoplasm |
| FLNA\|P21333\|FLNA_HUMAN\|1502 | NADGTQTVNYVPSR | caspase-3 | Endoplasmic reticulum |
| FLNA\|P21333\|FLNA_HUMAN\|1504 | DGTQTVNYVPSR | | Nucleus |
| FLNB\|O75369\|FLNB_HUMAN\|134 | DDAKKQTPKQR | | Cytoplasm |
| GET3\|O43681\|GET3_HUMAN\|166 | DTAPTGHTLR | | Cytoplasm |
| GJA1\|P17302\|CXA1_HUMAN\|47 | DEQSAFR | | Cytoplasm |
| GPC1\|P35052\|GPC1_HUMAN\|24 | DPASKSR | Signal Pepitde | Nucleus |
| GPC5\|P78333\|GPC5_HUMAN\|25 | EGVQTCEEVR | Signal Pepitde | Endoplasmic reticulum |
| GTF2I\|P78347\|GTF2I_HUMAN\|692 | NNNNPQTSAVR | | Mitochondrion |
| GTF2I\|P78347\|GTF2I_HUMAN\|693 | NNNPQTSAVR | | Nucleus |
| H1-4\|P10412\|H14_HUMAN\|3 | ETAPAAPAAPAPAEKTPVKKKAR | | Nucleus |
| HSP90B1\|P14625\|ENPL_HUMAN\|22 | DDEVDVDGTVEEDLGKSR | Signal Pepitde | Cytoplasm |
| HSP90B1\|P14625\|ENPL_HUMAN\|24 | EVDVDGTVEEDLGKSR | Signal Pepitde | Extracellular |
| HSP90B1\|P14625\|ENPL_HUMAN\|26 | DVDGTVEEDLGKSR | | Cell membrane |
| HSPA1A\|P0DMV8\|HS71A_HUMAN\|333 | DLVLVGGSTR | | Endoplasmic reticulum |

| Gene\|Uniprot Accession\|Uniprot Name\|Position | Sequence | cleavage_type | Predicted Localization |
|---|---|---|---|
| HSPB1\|P04792\|HSPB1_HUMAN\|64 | ESPAVAAPAYSR | | Cytoplasm |
| HSPH1\|Q92598\|HS105_HUMAN\|466 | NVSAQKDGEKSR | | Cytoplasm |
| HYOU1\|Q9Y4L1\|HYOU1_HUMAN\|713 | EDKLAQSVQKLQDLTLR | | Cell membrane |
| IGFBP4\|P22692\|IBP4_HUMAN\|22 | DEAIHCPPCSEEKLAR | Signal Pepitde | Cytoplasm |
| JMJD8\|Q96S16\|JMJD8_HUMAN\|24 | EGDGGWRPGGPGAVAEEER | Signal Pepitde | Cytoplasm |
| KANK2\|Q63ZY3\|KANK2_HUMAN\|257 | EDPVALETR | | Golgi apparatus |
| TAP1\|Q03518\|TAP1_HUMAN\|638 | EAGSQLSGGQR | caspase-5 | Cytoplasm |
| TCTN2\|Q96GX1\|TECT2_HUMAN\|26 | DLAFIPPFIR | Signal Pepitde | Extracellular |
| KRT7\|P08729\|K2C7_HUMAN\|70 | QSLLAPLR | | Endoplasmic reticulum |
| KRT8\|P05787\|K2C8_HUMAN\|69 | NQSLLSPLVLEVDPNIQAVR | | Cell membrane |
| KRT8\|P05787\|K2C8_HUMAN\|70 | QSLLSPLVLEVDPNIQAVR | | Lysosome/Vacuole |
| LAMA4\|Q16363\|LAMA4_HUMAN\|28 | DDNAFPFDIEGSSAVGR | Signal Pepitde | Cytoplasm |
| LMAN2\|Q12907\|LMAN2_HUMAN\|45 | DITDGNSEHLKR | Signal Pepitde | Endoplasmic reticulum |
| LMNA\|P02545\|LMNA_HUMAN\|231 | NGKQREFESR | caspase-5 | Endoplasmic reticulum |
| LRRC59\|Q96AG4\|LRC59_HUMAN\|223 | QAPKSKSGSRPR | | Cytoplasm |
| LSM3\|P62310\|LSM3_HUMAN\|7 | QQQTTNTVEEPLDLIR | caspase-3 | Cell membrane |
| LSM3\|P62310\|LSM3_HUMAN\|9 | QTTNTVEEPLDLIR | | Cytoplasm |
| LY6K\|Q17RY6\|LY6K_HUMAN\|18 | DANLTAR | Signal Pepitde | Cytoplasm |
| MAP4\|P27816\|MAP4_HUMAN\|12 | EPSPDIEGEIKR | | Endoplasmic reticulum |
| MEPCE\|Q7L2J0\|MEPCE_HUMAN\|247 | EGHVVLASPLKTGR | | Endoplasmic reticulum |
| MSN\|P26038\|MOES_HUMAN\|485 | ENGAEASADLR | caspase-3 | Endoplasmic reticulum |
| MTHFD2\|P13995\|MTDC_HUMAN\|36 | EAVVISGR | Transit Peptide | Mitochondrion |
| MYLK2\|Q9H1R3\|MYLK2_HUMAN\|542 | NLAEKAKR | | Endoplasmic reticulum |
| NARS1\|O43776\|SYNC_HUMAN\|5 | ELYVSDR | | Cytoplasm |
| NPM1\|P06748\|NPM_HUMAN\|209 | QNGKDSKPSSTPR | | Cytoplasm |
| OLFML2A\|Q68BL7\|OLM2A_HUMAN\|28 | DSKVFGDLDQVR | Signal Pepitde | Cytoplasm |
| OPTN\|Q96CV9\|OPTN_HUMAN\|251 | EVALKEAKER | | Cytoplasm |
| P4HA3\|Q7Z4N8\|P4HA3_HUMAN\|29 | DTFSALTSVAR | Signal Pepitde | Extracellular |
| P4HB\|P07237\|PDIA1_HUMAN\|18 | DAPEEEDHVLVLR | Signal Pepitde | Cytoplasm |
| P4HTM\|Q9NXG6\|P4HTM_HUMAN\|88 | DESSDPGPQHR | | Endoplasmic reticulum |
| PCNP\|Q8WW12\|PCNP_HUMAN\|3 | DGKAGDEKPEKSQR | | Endoplasmic reticulum |
| PCNP\|Q8WW12\|PCNP_HUMAN\|8 | DEKPEKSQR | | Cell membrane |
| PDIA3\|P30101\|PDIA3_HUMAN\|26 | DVLELTDDNFESR | Signal Pepitde | Cytoplasm |
| PDLIM5\|Q96HC4\|PDLI5_HUMAN\|302 | NTKKANNSQEPSPQLASSVASTR | | Cell membrane |
| PGRMC1\|O00264\|PGRC1_HUMAN\|49 | DQPAASGDSDDDEPPPLPR | | Cell membrane |
| PKN2\|Q16513\|PKN2_HUMAN\|207 | NAKPVISPLELR | | Cytoplasm |
| PLA2R1\|Q13018\|PLA2R_HUMAN\|21 | EGVAAALTPER | Signal Pepitde | Cell membrane |
| PPFIA1\|Q13136\|LIPA1_HUMAN\|222 | DINHEQENTPSTSGKR | | Endoplasmic reticulum |
| PPFIA1\|Q13136\|LIPA1_HUMAN\|837 | NSSQDALGLSKLGGQAEKNR | caspase-3 | Nucleus |
| PPP1R8\|Q12972\|PP1R8_HUMAN\|165 | NLTEFNTAHNKR | granzyme B | Cytoplasm |
| PRKDC\|P78527\|PRKDC_HUMAN\|2714 | NKVKGAAGR | caspase-3 | Cytoplasm |
| PRRC2A\|P48634\|PRC2A_HUMAN\|1466 | DQVIHSNPAGIQQALAQLSSR | | Cytoplasm |
| PRSS21\|Q9Y6M0\|TEST_HUMAN\|22 | ESQEAAPLSGPCGR | Signal Pepitde | Endoplasmic reticulum |
| PSMC2\|P35998\|PRS7_HUMAN\|3 | DYLGADQR | | Nucleus |
| PTBP1\|P26599\|PTBP1_HUMAN\|43 | DSKKFKGDSR | | Cytoplasm |
| RANBP1\|P43487\|RANG_HUMAN\|131 | ECPKPELLAIR | | Cell membrane |
| RCN2\|Q14257\|RCN2_HUMAN\|26 | EELHYPLGER | Signal Pepitde | Cell membrane |
| RPL35\|P42766\|RL35_HUMAN\|23 | DLKVELSQLR | meprin alpha subunit | Cytoplasm |
| RPL35\|P42766\|RL35_HUMAN\|62 | NQTQKENLR | | Extracellular |

| Gene\|Uniprot Accession\|Uniprot Name\|Position | Sequence | cleavage_type | Predicted Localization |
|---|---|---|---|
| RPL35\|P42766\|RL35_HUMAN\|63 | QTQKENLR | | Cytoplasm |
| RPS6\|P62753\|RS6_HUMAN\|110 | NLVIVKKGEKDIPGLTDTTVPR | | Extracellular |
| SART1\|O43290\|SNUT1_HUMAN\|646 | QNKGLLETTVQKVAR | | Cytoplasm |
| SCAMP3\|O14828\|SCAM3_HUMAN\|22 | QDPAVIQHRPSR | | Cytoplasm |
| SCD\|O00767\|SCD_HUMAN\|142 | NTMAFQNDVYEWAR | | Cytoplasm |
| SCD\|O00767\|SCD_HUMAN\|7 | QDDISSSYTTTTTITAPPSR | | Cytoplasm |
| SEC22B\|O75396\|SC22B_HUMAN\|152 | NIEEVLQR | | Cytoplasm |
| SERBP1\|Q8NC51\|PAIRB_HUMAN\|7 | EGFGCVVTNR | meprin beta subunit | Cytoplasm |
| SHC1\|P29353\|SHC1_HUMAN\|112 | NKLSGGGGRR | | Cytoplasm |
| SHMT2\|P34897\|GLYM_HUMAN\|27 | QHSNAAQTQTGEANR | Transit Peptide | Cell membrane |
| SHOC2\|Q9UQ13\|SHOC2_HUMAN\|70 | NTIKRPNPAPGTR | | Cytoplasm |
| SLC25A24\|Q6NUK1\|SCMC1_HUMAN\|16 | QDAEQPTRYETLFQALDR | | Cytoplasm |
| SMC1B\|Q8NDV3\|SMC1B_HUMAN\|1110 | NCVAPGKR | | Nucleus |
| TSPYL2\|Q9H2G4\|TSYL2_HUMAN\|6 | EGPPAKTRR | | Cell membrane |
| SMC2\|O95347\|SMC2_HUMAN\|238 | QFLLAEDTKVR | | Cell membrane |
| SMC2\|O95347\|SMC2_HUMAN\|989 | NVLTEAEER | | Nucleus |
| SNAP29\|O95721\|SNP29_HUMAN\|36 | DAPADRQQYLR | | Nucleus |
| SPTAN1\|Q13813\|SPTN1_HUMAN\|678 | NQQQQFNR | | Cytoplasm |
| SREBF2\|Q12772\|SRBP2_HUMAN\|801 | QAFCKNLLER | | Nucleus |
| SRP68\|Q9UHB9\|SRP68_HUMAN\|6 | QVPGGGGGGGSGGGGGSGGG GSGGGR | | Endoplasmic reticulum |
| SSBP1\|Q04837\|SSBP_HUMAN\|17 | ESETTTSLVLER | Transit Peptide | Extracellular |
| TAP1\|Q03518\|TAP1_HUMAN\|638 | EAGSQLSGGQR | caspase-5 | Cytoplasm |
| TCTN2\|Q96GX1\|TECT2_HUMAN\|26 | DLAFIPPFIR | Signal Pepitde | Extracellular |
| THBS1\|P07996\|TSP1_HUMAN\|19 | NRIPESGGDNSVFDIFELTGAAR | Signal Pepitde | Nucleus |
| THUMPD3\|Q9BV44\|THUM3_HUMAN\|214 | ESSKEETEPQVLKFR | caspase-3 | Endoplasmic reticulum |
| TM9SF2\|Q99805\|TM9S2_HUMAN\|166 | DVEDGQRF | | Cytoplasm |
| TMEM59\|Q9BXS4\|TMM59_HUMAN\|36 | EAFDSVLGDTASCHR | Signal Pepitde | Nucleus |
| TMEM9\|Q9P0T7\|TMEM9_HUMAN\|21 | NKSSEDIR | Signal Pepitde | Cytoplasm |
| TNFRSF12A\|Q9NP84\|TNR12_HUMAN\|28 | EQAPGTAPCSR | Signal Pepitde | Endoplasmic reticulum |
| TSPYL2\|Q9H2G4\|TSYL2_HUMAN\|6 | EGPPAKTRR | | Cell membrane |
| TTN\|Q8WZ42\|TITIN_HUMAN\|3157 | QVIEKQR | | Endoplasmic reticulum |
| UBAP2L\|Q14157\|UBP2L_HUMAN\|22 | QTQHKQRPQATAEQIR | | Nucleus |
| UBE2O\|Q9C0C9\|UBE2O_HUMAN\|490 | DTSSVTSSASSTTSSQSGSGTSR | | Endoplasmic reticulum |
| UBR4\|Q5T4S7\|UBR4_HUMAN\|2313 | NAQQIKHR | | Cytoplasm |
| UQCRHL\|A0A096LP55\|QCR6L_HUMAN\|15 | DPEEEEEEEEELVDPLTTVR | Transit Peptide | Mitochondrion |
| VIM\|P08670\|VIME_HUMAN\|259 | DVSKPDLTAALR | meprin beta subunit | Cytoplasm |
| VIM\|P08670\|VIME_HUMAN\|93 | NTEFKNTR | | Mitochondrion |
| VTI1B\|Q9UEU0\|VTI1B_HUMAN\|158 | QIGSEIIEELGEQR | | Cytoplasm |
| YBX1\|P67809\|YBOX1_HUMAN\|223 | NQGAGEQGRPVR | meprin alpha subunit | Extracellular |

### Supplementary Table 2. Primer list of Nt-arginylation protein candidates

| Gene | Primer | Sequences of primer pairs |
|---|---|---|
| ERO1A | Forward | 5' - CCGCTCGAGGCCACCATGGGCCGCGGCTGGGGATTC - 3' |
| | Reverse | 5' - GCTCTAGATTATGAATATTCTGTAACAAGTTCCTGAAGTTTTCTAATTC - 3' |
| THBS1 | Forward | 5' - CCGCTCGAGGCCACCATGGGGCTGGCCTGGGGACTAGG - 3' |
| | Reverse | 5' - GCTCTAGATTGGGATCTCTACATTCGTATTTCAGGTCAGAGAAGAACACC - 3' |
| CD55 | Forward | 5' - CCGCTCGAGGCCACCATGACCGTCGCGCGGCCGAGCG - 3' |
| | Reverse | 5' - GCTCTAGATTAGAAACTAGGAACAGTCTGTATACTTGTGTGTATTTTCTTC - 3' |
| PDIA3 | Forward | 5' - CCGCTCGAGGCCACCATGCGCCTCCGCCGCCTAGCGC - 3' |
| | Reverse | 5' - GCTCTAGATTGAGATCCTCCTGTGCCTTCTTCTTCTTCTTGGG - 3' |
| SLC25A24 | Forward | 5' - CCGCTCGAGGCCACCATGTTGCGCTGGCTGCGGGAC - 3' |
| | Reverse | 5' - GCTCTAGATTTTTCTGGGTTACTCCTAAAGTTTGCTTCATATTTTC - 3' |
| MTHFD2 | Forward | 5' - CCGCTCGAGGCCACCATGAAACCAGCTTCAATTTCAGAGG - 3' |
| | Reverse | 5' - GCTCTAGATTATTAGTGGCTACCCCAAGCTCTTTAGACTTCAGCACTTC - 3' |
| PPP1R8 | Forward | 5' - CCGCTCGAGGCCACCATGGGTGGAGAGGATGATGAACTCAAGGG - 3' |
| | Reverse | 5' - GCTCTAGATTAATCAGCAAGGAAGGTGTGGGCTTCTTGCCTGGCC - 3' |
| HSP90B1 | Forward | 5' - CCGCTCGAGGCCACCATGAGGGCCCTGTGGGTGCTGGG - 3' |
| | Reverse | 5' - GCTCTAGATTCAATTCATCTTTTTCAGCTGTAGATTCCTTTGC - 3' |
| EIF4B | Forward | 5' - GATATCGCCACCATGGCGGCCTCAGCAAAAAAGAAGAATAAG - 3' |
| | Reverse | 5' - ATAGTTTAGCGGCCGCTTTCGGCATAATCTTCTCCCTCATTTTCATCTTCACC - 3' |
| CALU | Forward | 5' - GATATCGCCACCATGAAGGAAACTGATCTAATTATCATGG - 3' |
| | Reverse | 5' - ATAGTTTAGCGGCCGCTGAACTCATCATGCCGTACTAAGGCC - 3' |
| LMAN2 | Forward | 5' - CCGCTCGAGGCCACCATGGCGGCGGAAGGCTGGATTTGG - 3' |
| | Reverse | 5' - GCTCTAGATTGTAGAAGCGCTTGTTCCGCTCCTGCCGCTTCTGG - 3' |
| NPM1 | Forward | 5' - CCGCTCGAGGCCACCATGGAAGATTCGATGGACATGG - 3' |
| | Reverse | 5' - GCTCTAGATTAAGAGACTTCCTCCACTGCCAGAGATCTTGAATAGCC - 3' |
| RGS4 | Forward | 5' - AAAGAATTCGCCACCATGTATAATATGATGCTTCTAATCCAAAAGAGG - 3' |
| | Reverse | 5' - TTTCTCGAGGGCACACTGAGGGACCAGGGAAGCAC - 3' |
| R-HSPA5 | Forward | 5' - AAAAAGCTTGCCACCATGAAGCTCTCCCTGGTGGCCGCGATGC - 3' |
| | Reverse | 5' - TTTGGATCCCAACTCATCTTTTTCTGCTGTATCCTCTTCACCAGTTGGG - 3' |
